# A flexible, modular and versatile functional part assembly toolkit for gene cluster engineering in *Streptomyces*

**DOI:** 10.1101/2023.09.20.558588

**Authors:** Xuejin Zhao, Yeqing Zong, Qiuli Lou, Chenrui Qin, Chunbo Lou

**Affiliations:** CAS Key Laboratory of Microbial Physiological and Metabolic Engineering, Institute of Microbiology, Chinese Academy of Sciences, Beijing, 100101, China; CAS Key Laboratory of Pathogen Microbiology and Immunology, Institute of Microbiology, Chinese Academy of Sciences, Beijing, 100101, China; Center for Cell and Gene Circuit Design, Key Laboratory of Quantitative Synthetic Biology, Shenzhen Institute of Synthetic Biology, Shenzhen Institutes of Advanced Technology, Chinese Academy of Sciences, 1068 Xueyuan Avenue, University Town, Nanshan, Shenzhen 518055, China; Peking-Tsinghua Joint Center for Life Sciences, Peking University, Beijing 100871, China; Center for Quantitative Biology, Academy for Advanced Interdisciplinary Studies, School of Physics, Peking University, Beijing 100871, China; College of Life Sciences, University of Chinese Academy of Sciences, Beijing, 100149, China

**Keywords:** *Streptomyces*, standard toolkit, gene cluster, heterologous expression, synthetic biology

## Abstract

*Streptomyces* has enormous potential to produce novel natural products (NPs) as it harbors a huge reservoir of uncharacterized and silent natural product biosynthetic gene clusters (BGCs). However, the lack of efficient gene cluster engineering strategies has hampered the pace of new drug discovery. Here, we developed an easy-to-use, highly flexible DNA assembly toolkit for gene cluster engineering. The DNA assembly toolkit are compatible with various DNA assembling approaches including Biobrick, Golden Gate, CATCH, yeast homologous recombination-based DNA assembly and homing endonuclease-mediated assembly. This compatibility offers great flexibility in handling multiple genetic parts or refactoring large gene clusters. To demonstrate the utility of this toolkit, we quantified a library of modular regulatory parts, and engineered a gene cluster (*act*) using characterized promoters that led to increased production. Overall, this work provides a powerful part assembly toolkit that can be used for natural product discovery and optimization in *Streptomyces*.

## INTRODUCTION

*Streptomyces* is one of the most industrial genera used to produce natural products, such as antitumor drugs, immunosuppressants, herbicides and especially antibiotics (Demain, 2014). Recent advances in genome sequencing have revealed a vast unexploited resource of gene clusters (BGCs) within *Streptomyces*, holding great potential for the discovery of novel bioactive compounds (Liu et al., 2021). Strategies to activate these gene clusters involve cloning and refactoring, and heterologous expression in well-characterized *Streptomyces* hosts, aiming to overcome limitations posed by individual activation methods (Wang et al., 2021; Zhuo et al., 2017). However, this approach is currently limited by the lack of standard and versatile toolkits and assembly technologies. With the rapid development of synthetic biology, the construction of complex synthetic biological systems has developed from monocistronic gene expression to multi-step pathways or even chromosomes (Gibson et al., 2009; Ostrov et al., 2016). Accordingly, the demands of DNA assembly have shifted from single fragments to more intricate multi-fragments or large gene clusters. To address this need, several modern assembly techniques have been developed, including BioBrick (Smolke, 2009), BglBrick (Lee et al., 2011), Golden Gate (Engler et al., 2009) and Gibson assembly(Gibson et al., 2009). Notably, Cas9-Assisted Targeting of CHromosome segments (CATCH) (Jiang et al., 2015) and Transformation-Associated Recombination (TAR) (Yamanaka et al., 2014) have proven effective for cloning large gene clusters. Moreover, yeast homologous recombination-based methods such as mCRISTAR(Kang et al., 2016), miCASTAR(Kim et al., 2019) and mpCRISTAR(Kim et al., 2020) have been developed for multi-site gene cluster editing.

Although genetic engineering approaches have long been employed to boost the secondary metabolite production in *Streptomyces* for decades, and several vector platforms are commonly used, such as the pIJ family (Bierman et al., 1992). However, these vector systems are primarily limited to operate single gene, which limits their application in multi-gene biosynthetic pathways. In addition, these platforms are incompatible with the advanced standard and modular assembly approaches such as Biobrick and Golden Gate. Additionally, their reliance on ϕC31or ϕBT1 integrated system restricts efficient combinatorial engineering in *Streptomyces*. Moreover, the size of many BGCs of natural products exceeds the capacity to process them. Although several derivative vectors such as pStreptoBAC(Miao et al., 2005)and pSBAC(Liu et al., 2009) have been developed for cloning large gene clusters, they are not standard and modular vectors, and present challenges for subsequent modification. Recent efforts have led to the development of a suite of 45 orthogonal integration vectors based on different site-specific integration systems for heterologous biosynthetic pathways in *Streptomyces venezuelae* (Phelan et al., 2017). However, these vectors are designed around traditional restriction sites and pMB1 *Escherichiacoli* replicon, making them unsuitable for assembling large gene clusters. Similarly, a set of 12 standardized modular plasmids has been designed to facilitate the assembly of natural product BGCs, but only allow the iterative assembly of genes (or gene cassettes) using the BioBrick assembly method. In addition, they are all based on the *E. coli* p15A replicon, which limits the capacity of the cloned natural product BGCs (Aubry et al., 2019).

In this work, we design a flexible and modular DNA assembly strategy that is easily to exchange plasmid copy number, selection marker gene, integration site, regulatory and catalytic parts of gene cluster. More importantly, it enables cloning and editing of different-sized gene clusters using various cloning methods such as CATCH and yeast homologous recombination-based DNA assembly. Base on the assembly toolkit, we have developed a high-throughput experimental pipeline for testing regulatory elements, and identified a library of modular promoters. In addition, we have demonstrated the advantage of the assembly strategy to enhance the production of a large gene cluster.

## RESULTS AND DISCUSSION

### Design and construction the modular DNA assembly toolkit

One of the barriers to activate the silent gene clusters is the lack of ability to manipulate the multiple regulatory and multiple catalytic parts in a cross-species manner. Here, we developed a flexible, modular and versatile DNA assembly toolkit for gene cluster engineering. All vectors in the assembly toolkit include a cargo part (part 1) and an essential part (part 2). The cargo part (part 1) was composed by the cloning sites for the regulatory parts and catalytic pats of the gene clusters, while the essential part (part 2) was responsible for the replication origins of the plasmid, selection marker genes in *E. coli*, yeast and *Streptomyces* (Figure 1).

**Figure 1.**
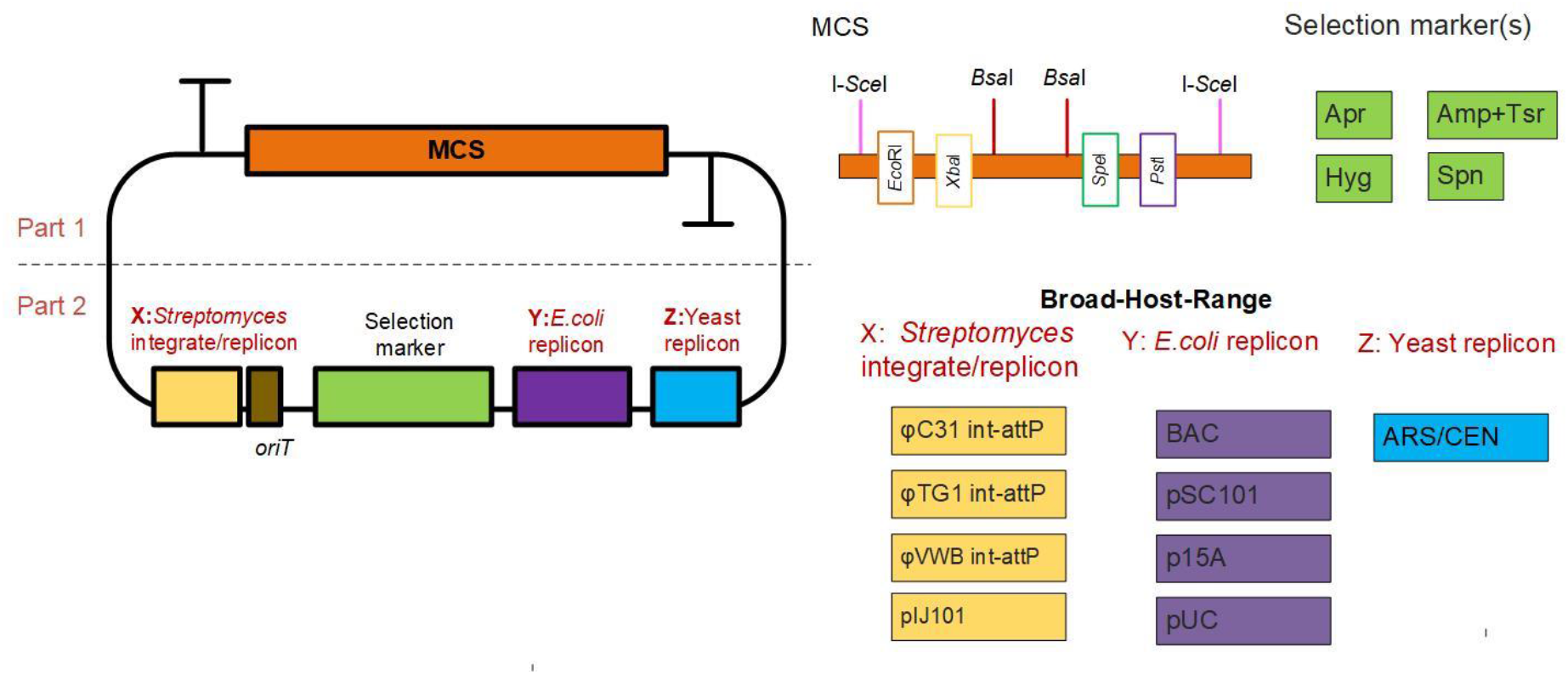
The modular DNA assembly toolkit. Left part shows the structural organization of the vectors in the toolkit. The right part depicts the details of the MCS, selection marker(s), *Streptomyces* integrate/replicon, *E. coli* and yeast replicon. Apr, apramycin; Amp, ampicillin; Hyg, hygromycin; Spn, spectinomycin; Tsr, thiostrepton.

The cargo part contains a multiple cloning site (MCS) and two transcription terminators (T1 and T2). In order to be compatible with a number of standard cloning systems, the MCS was designed as follows:I-*Sce*I–*Eco*RI-–*Xba*I–*Bsa*I-*Bsa*I–*Spe*I–*Pst*I–I-*Sce*I. The *Eco*RI, *Spe*I, *Xba*I and *Pst*I sites allow biobrick assembly, and two *Bsa*I sites allow Golden Gate assembly. The two homing endonucleases I*-Sce*I sites, which recognize long DNA sequences (18 bp) are suitable for exchanging large fragments. To avoid unwanted transcriptional read-through into other elements of the backbone, two transcriptional terminators, T1 and T2 were inserted into the up and down of MCS (Figure 1).

The essential part consists of four modules, which are arranged in the same order in all vectors (1) integration/replication module in *Streptomyces*; (2) antibiotic markers; (3) replication module in *E. coli*; (4) replication module in yeast (Figure 1). To ensure the ability to accommodate large gene clusters, the vectors are mainly based on medium or low copy number replicons, including p15A, pSC101 and BAC in *E. coli*, and ARS/CEN origin in yeast. For the hierarchical construction of gene clusters, more than one vector should be used in a single host. Therefore, four antibiotic marker cassettes (Apr, Hyg, Spn, and Tsr+Amp), four integration/ replicon elements (ϕC31, TG1, VWB and pIJ101) for four commonly used *Streptomyces* hosts (*Streptomyces coelicolor, Streptomyces lividans, Streptomyces albus* and *S. venezuelae*) were included (Figure 1, Figure S1).

As a result, a total of 10 basic vectors were constructed (Table 1). In general, large DNA gene clusters are more stable in a low-copy vector. In our toolkit, the pPAB with BAC replicon is primarily designed for assembling large sized gene clusters, especially > 50kb, the pPAS, pVHS and pTHS with pSC101 replicon are designed for the assemble medium sized gene clusters (<50 kb), the pPAP, pVSP, pTSP and pIATP with p15A replicon are used to assemble <30 kb gene clusters, and the pIATU with pUC replicon is suable to small size gene clusters (<10 kb). The detailed information of all vectors maps and sequences can be found in Figure S2 and Table S1.

**Table 1.**
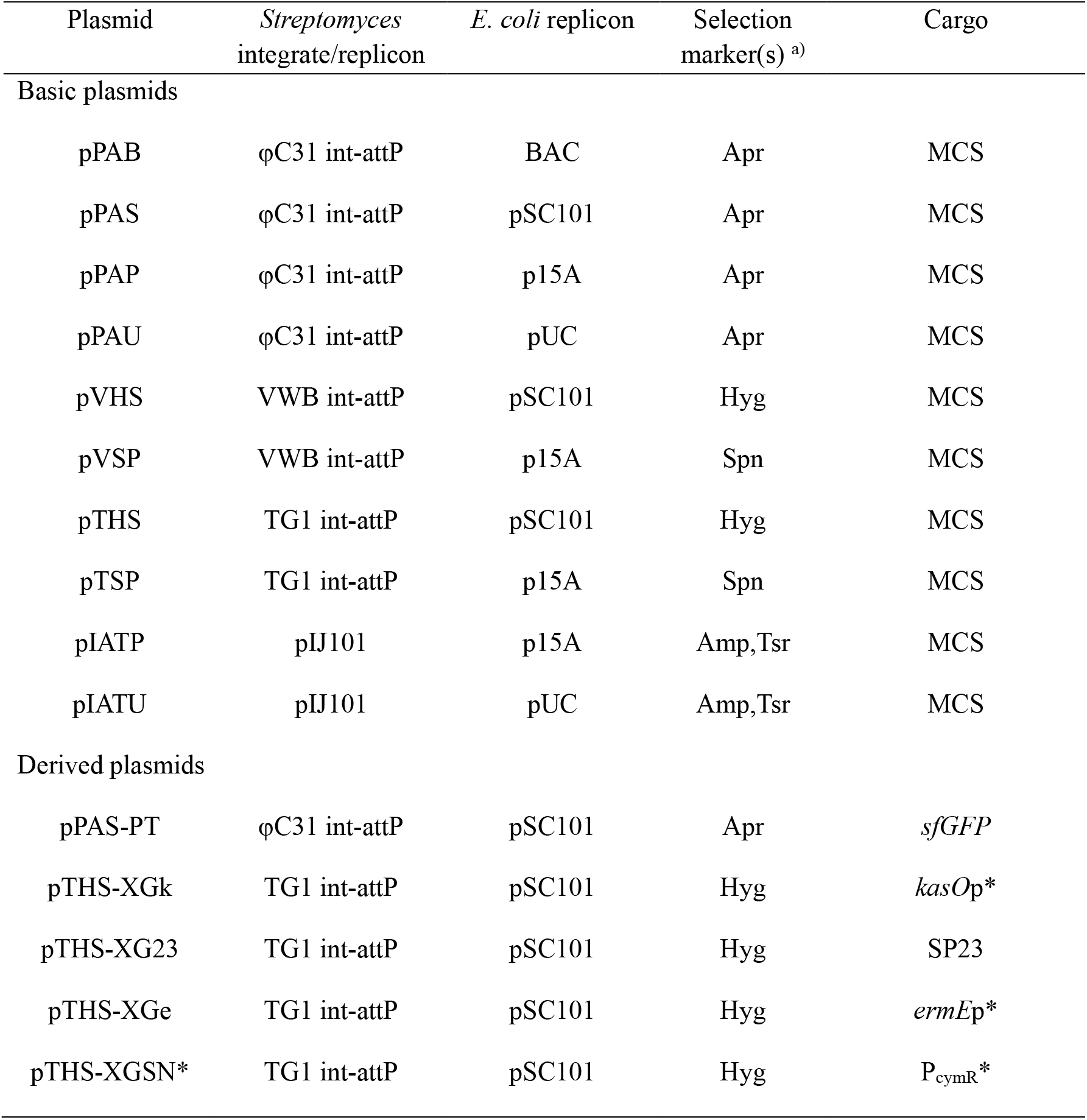
List of 10 basic plasmids and 5 derivatives.

The 10 basic vectors were then conjugated from *E. coli* into *S. coelicolor, S. lividans, S. albus* and *S. venezuelae*. At the same time, commonly used plasmid pIJ8660 was used as a positive control. As shown in Figure 2, the conjugal frequency of the majority of the plasmids was greater than 10^-6^, with the exception of the following cases. (1) *S. lividans* was found to be resistant to spectinomycin. Therefore, the plasmids with the spectinomycin antibiotic marker cannot be used in it. (2) *S. albus* has a high level of tolerance to spectinomycin. Therefore, the concentration of spectinomycin used in *S. albus* is higher than that used in other hosts.

**Figure 2.**
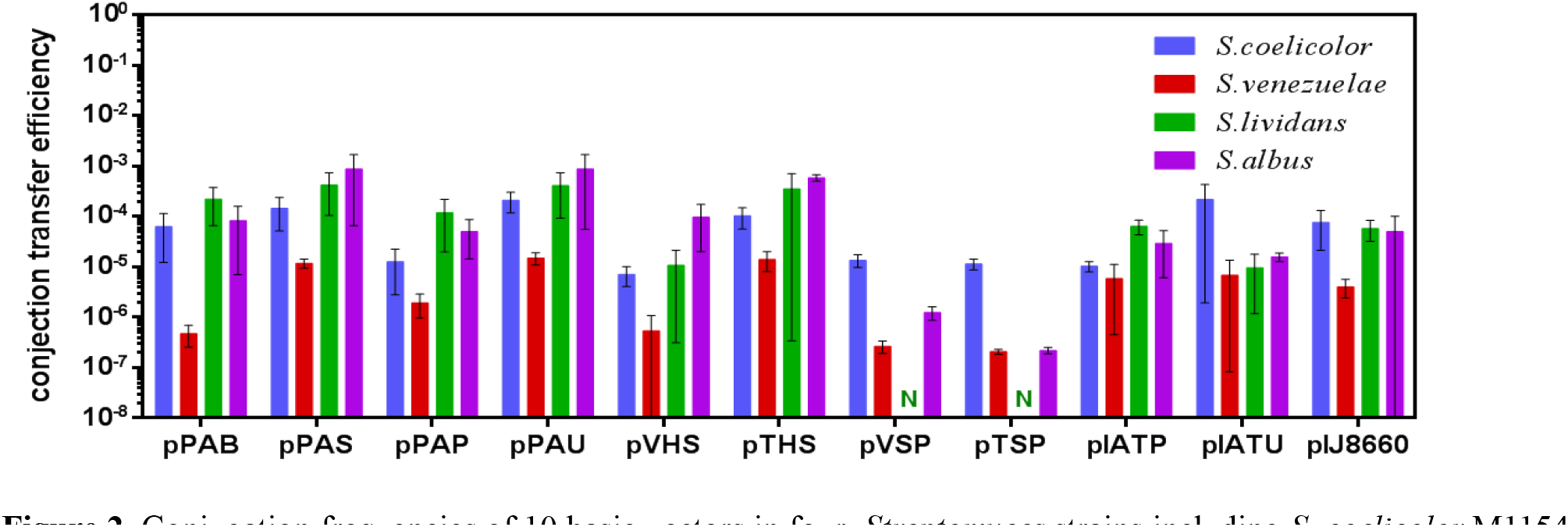
Conjugation frequencies of 10 basic vectors in four *Streptomyces* strains including *S. coelicolor* M1154, *S. venezuelae* ISP5230, *S. lividans* TK24 and *S. albus* J1074.

### Demonstration of compatible molecular operations

The BioBrick and Golden Gate are the most widely used DNA assembly methods due to their robustness and modularity. BioBrick provides a straightforward way to combine standardized biological components, and Golden Gate assembly allows scarless, multi-part DNA assembly. However, they have the limitation that all the parts used must be free of the enzymes used, which makes them unsuitable for assembling gene clusters due to the frequent presence of restriction sites in the target sequence. Therefore, some CRISPR/Cas9-based methods developed to clone and refactor large gene clusters. In addition,I-*Sce*I-based assembly is suitable for exchanging large gene clusters due to the long recognition sequence 8 bp).

The assembly toolkit in our work is compatible with a wide range of DNA assembly approaches mentioned above. All vectors in the toolkit support the assembly of standard BioBrick parts according to the procedure shown in Figure 3A. For example, an anther fragment can be easily ligated to the standard plasmids containing a fragment as follows: The first step is to digest the donor part with *Spe*I and *Pst*I and the target vector with *Eco*RI and *Xba*I. Then, the donor part and the target vector are ligated using T4 DNA ligase to generate the recombinant plasmid. For Golden Gate assembly, PCR products or fragment inserts in donor vectors flanked by specific four-base overhangs and *Bsa*I restriction sites, or double-stranded DNA oligos with appropriate overhangs could be assembled into the vector backbone containing two *Bsa*I restriction sites (Figure 3B). In addition, the plasmid toolkit allows rapid assembly of gene clusters using the CATCH method. As shown in Figure 3C, the first step is to introduce homologous arms corresponding to the two ends of the gene cluster into the vector backbone using the Golden Gate assembly. The resulting plasmid was then linearized with *Aar*I. Finally, the linearized capture vector was assembled with the gene cluster fragment from Cas9-treated bacterial genomic DNA by Gibson assembly. Furthermore, all vectors contain a yeast ARS/CEN origin to support gene cluster editing in yeast (Figure 3D). Recombinant plasmids carrying gene clusters are easily edited in yeast using homologous recombination-based methods such as mCRISTAR, miCASTAR or mpCRISTAR. Moreover, all vectors carry two I-*Sce*I recognition sites. This means that a gene cluster in one vector can be easily assembled into another vector as follows: the gene cluster was digested with I-*Sce*I from original plasmid, and the destination vector was digested with *Bsa*I. The digested gene cluster was then assembled into the linearized destination vector using Gibson assembly (Figure 3E).

**Figure 3.**
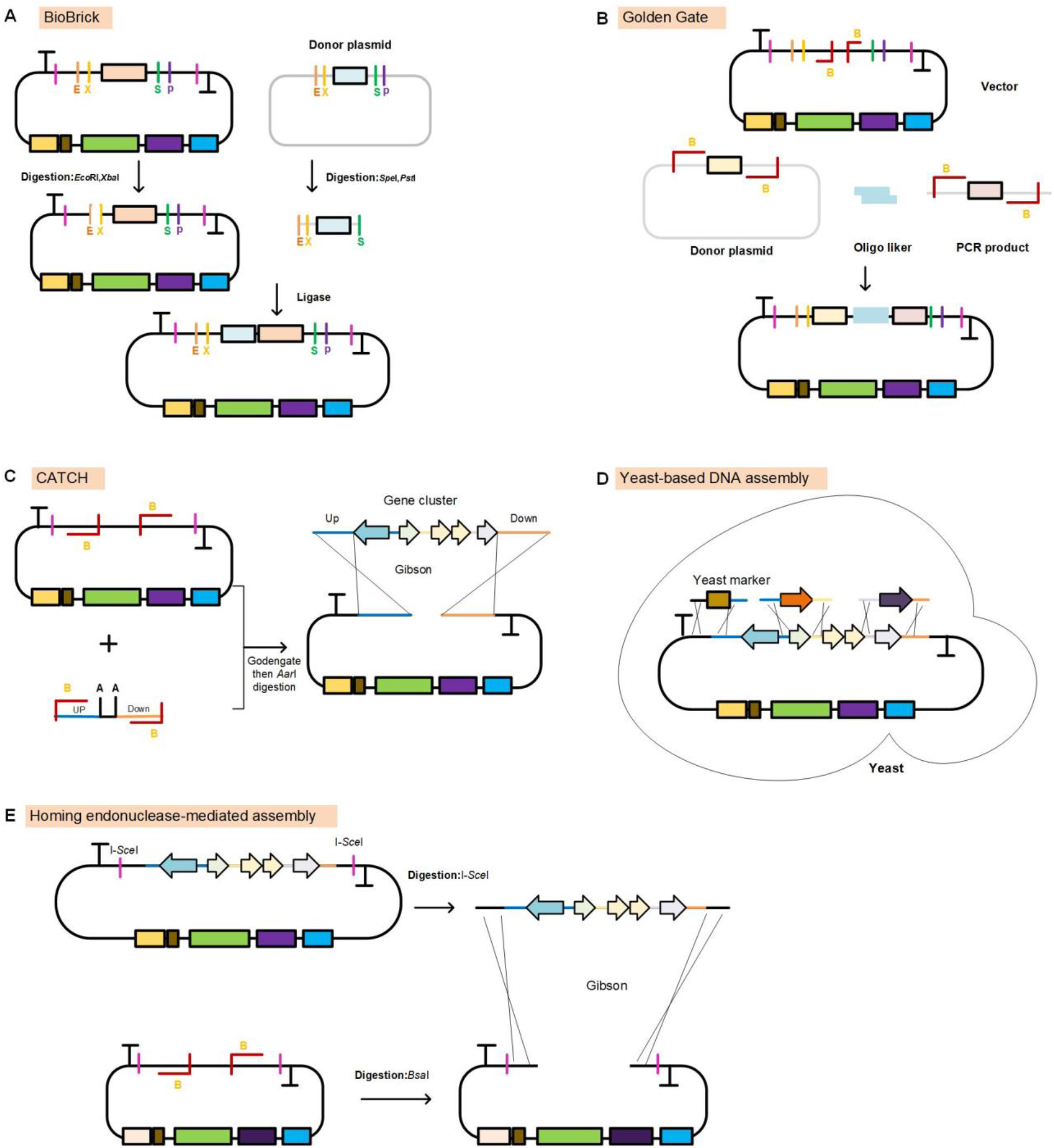
Demonstration of compatible molecular operations. A. Biobrick standard assembly . B. Golden Gate assembly.C. Direct cloning gene cluster using CATCH method. D. Schematic of homologous recombination-based gene cluster editing in yeast. E. Vector exchange for large gene cluster based on I-*Sce*I-based assembly.

### Development of an extensible assembly toolkit and parts library for gene cluster engineering

Controlled gene expression is import for improvement of secondary metabolite production. However, there is currently no easy-to-use DNA assembly toolbox that enables controlled gene expression in *Streptomyces*. In order to facilitate optimization gene expression at the desired level, we developed a rapid, simple, and standardized gene expression assembly vectors based on pTHS, including three constitutive expression vectors and one inducible expression vector compatible with the Goden gate cloning method. Of these, pTHS-XGe, pTHS-XGk and pTHS-XG23 carry constitutive promoters with different strengths (*ermE*p*, *KasO*p*and SP23) (Bai et al., 2015), and pTHS-XGSN* contains a tight cumate-inducible promoter P_cymR*_ (Zhao et al., 2022) (Figure 4A).

**Figure 4.**
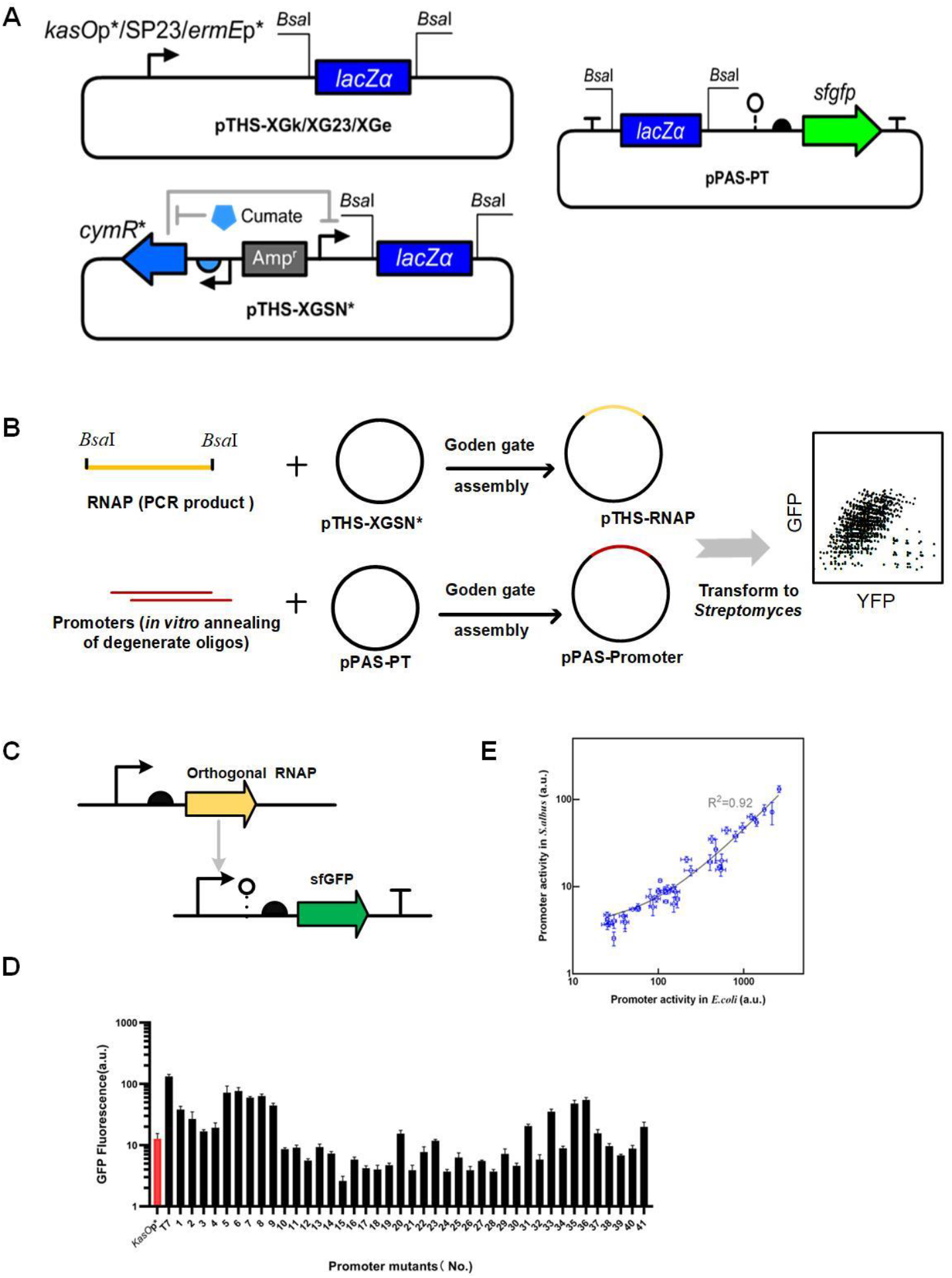
Derived vectors and their application for promoter activity analysis. A. The maps of three constitute expression plasmids, one inducible expression plasmid and one reporter plasmid. B.promoter library was constructed on the sfGFP reporter vector, and RNAP was constructed on modularized expression vector using Golden gate assembly. C. The structure of genetic circuit used for orthogonal transcription system characterizing. D. Analysis of fluorescence intensities of the T7 promoter library using flow cytometry-based quantitative method. *KasO*p* was used as the the positive control(red). F. Linearity between expression in *S. albus* and *E*.*coli* transformed with T7 RNAP* and T7 promoters.

In order to fine-tune and enable combinatorial expression each gene (operon) of gene cluster, regulatory elements with a broad range of strengths need to be implemented. To rapid assembly and characterizing of regulatory elements, we constructed a sfGFP reporter vector pPAS-PT (Figure 4A). Using pTHS-XGSN* and pPAS-PT, we identified a panel of modular T7 promoters, where pTHS-XGSN* expresses T7 RNA polymerase (T7 RNAP*), and pPAS-PT allows easy assessment of T7 promoters strength. The T7 RNAP* encoding gene was cloned into pTHS-XGSN* using Golden gate assembly, resulting in pTHS-T7RNAP*. As well as, 41 chosen T7 promoters (Zong et al., 2017), were cloned into pPAS-PT using Golden gate assembly, resulting in pPAS-P_T7(X)_ (Figure 4B, C). pTHS-T7RNAP* was first introduced into *S*.*albus*, resulting in *S*.*a/*T7RNAP* .The pPAS-P_T7(X)_ plasmids were then introduced into *S*.*a/*T7RNAP*. The promoter activities were quantified by flow cytometry. As shown in Figure 4D, the T7 promoter exhibits a very strong activity in *S*.*albus*, and 15 promoters were found to be stronger than the commonly used strong promoter *kas*Op*. Particularly, the wild-type T7 promoter was the strongest of these promoters, reaching approximately 10-fold *kas*Op* (Figure 4D). At the same time, we found that the activities of of T7 promoters in *Streptomyces* showed a good linear correlation with the expression level in *E. coli* host (R^2^=0.92), demonstrating the highly modular feature of the T7 promoters (Figure 4E). These results indicated that our toolkit is effective for high-through analysis of regulatory elements, and the corresponding characterized modular promoters could be used to express genes or gene clusters at different levels as required in *Streptomyces*.

### Applying the assembly toolkit to clone and refactor gene cluster

To illustrate the powerful capabilities of our toolkit for gene clusters engineering, we cloned the 26-Kb actinorhodin (*act*) gene cluster in pPAB vector using the CATCH method as follows. First, two 30 bp DNA fragments from each side of the *act* were cloned into pPAB to generate pPAB-HS. Then, the 26 kb *act* BGC digested from *S. coelicolor* M145 genome by CRISPR/Cas9 was cloned directly into the pPAB-HS using Gibson, resulting in pPAB-act (Figure 5A). In order to improve the production of actinorhodin, the regulatory network of *act* gene cluster was refactored by inserting well-characterized promoters (P1-P2) to control the actVA and *act*II operons, P3 promoter to control the *act*?-orf2, and (P4-P5) to control the *act*III and *act*I operons (Figure 5B). For *act* gene cluster editing, the pPAB-act was first digested at the native promoter sites of gene cluster with Cas9 and sgRNAs i*n vitro*. Then the digested fragments, the DNA cassettes containing synthetic promoters with homology arms matching each digestion site in the gene cluster, and the URA marker were co-transformed into yeast. Transformants were plated on yeast synthetic dropout media missing uracil, and verified by PCR. The corrected recombinant plasmid pPAB-act2 and original plasmid pPAB-act were introduced into *S. albus* to generate *S*.*a*/*act2* and *S*.*a*/*act*, respectively. As shown in Figure 5B, the production of actinorhodin by *S*.*a*/*act2* resulted in a more intense blue pigment than that of *S*.*a*/*act*. These results demonstrate that our assembly toolkit is effective for gene cluster engineering, and shows excellent performance achieved by engineering gene clusters using desired regulatory elements.

**Figure 5.**
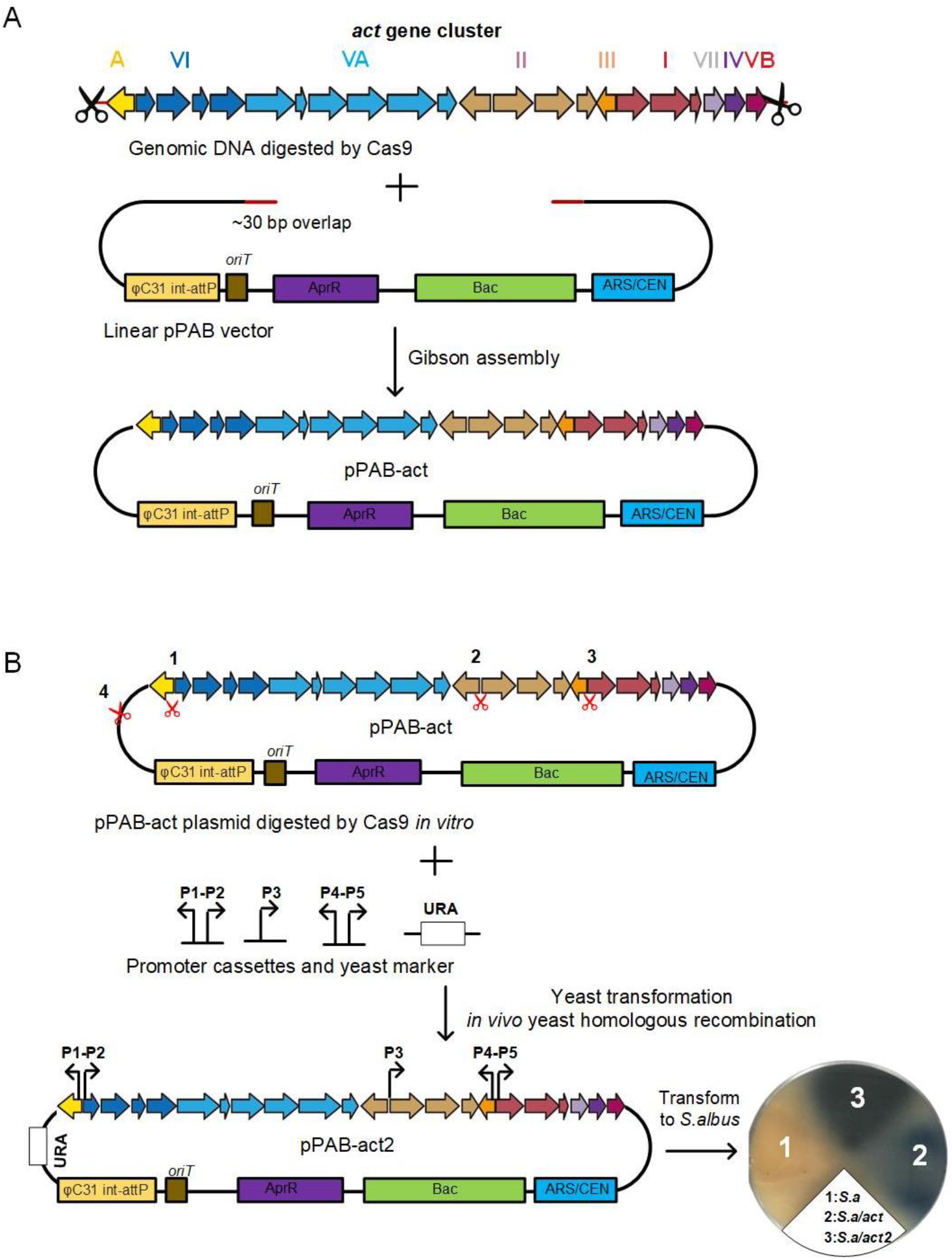
Cloning and editing of the *act* gene cluster. A.The *act* gene cluster was cloned into pPAB vector using CATCH method. B. The *act* gene cluster was reconstructed using well characterized promoters.

## COUCLUSION

In this work, we have developed an attractive assembly toolkit for engineering of gene cluster. Using this toolkit, a modular promoter library for tune gene expression was characterized by a rapid and efficient method. In addition, the assembly toolkit allows manipulation and optimization of large gene clusters and has been successfully used to clone and refactor the *act* BGC, resulting in improved actinorhodin production. Taken together, the modular plasmid toolkit provides a complete solution for manipulating gene clusters, and expands the plasmid toolkit for synthetic biology research in *Streptomyces*. The sequence of all the vectors can be found in the Supplementary Material. We hope that this will help to unlock the potential of the huge gene clusters from *Streptomyces* through synthetic biology and biotechnology.

## MATERIALS AND METHODS

### Strains, plasmids, and media

*E. coli* strains were grown at 37°C in LB medium (10 g/L tryptone, 5 g/L yeast extract and 10 g/L NaCl) with the corresponding antibiotic. *S. coelicolor, S. albus and S. lividans* strains were culture on the MS medium (20 g/L soybean flour, 20 g/L mannitol, and 20 g/L agar) for sporulation and conjugation. *S. venezuelae* strains were culture on the MYM medium (4.2 g/L (D-(+)-maltose monohydrate, 4 g/L yeast extract, 4 g/L malt extract, and 20 g/L agar) for sporulation and conjugation. The antibiotics used in this work were ampicillin (100 μg/ml for *E. coli*), thiostrepton (30 g/ml for *Streptomyces*), apramycin (50 μg/mL for *E. coli*, 50 μg/mL for *Streptomyces*), hygromycin (150 μg/mL for *E. coli*, 50 μg/mL for *Streptomyces*), spectinomycin (80 μg/mL for *E. coli*, 120 μg/ mL for *S. coelicolor* and *S. venezuelae*, and 200 μg/mL for *S. albus*).

### Conjugation of the vectors in *Streptomyces* species

All the vectors constructed were introduced into four *Streptomyces species* by conjugation according to the standard procedure(Flett et al., 1997). Exconjugants were selected on MS or MYM agar containing the appropriate antibiotic and 25 μg/mL nalidixic acid.

### Characterization of modular promoters

To test the T7 promoters, two plasmids pTHS-XGSN* and pPAS-PT were used as basic vectors to carry the T7 RNA polymerase and T7 promoters. To efficiently express the T7 RNAP in *Streptomyces*, the four rare TTA _leu_ codons were replaced by CTC _leu_, and the mutated T7 RNAP* sequence was listed in Table S3. The T7 RNAP* fragment amplified with T7 RNAP*-F and T7 RNAP*-R primers was cloned into pTHS-XGSN* using the Goden Gate method. All primers and oligonucleotides were synthesized by Generay Biotechnology (Table S4).The pPAS-P_T7(X)_ series plasmids were constructed via Oligonucleotide Linkers-Mediated Assemble (OLMA) approach developed in our lab (Zhang et al., 2015). Promoter DNA fragments were acquired by annealed and phosphorylated oligonucleotides, which have sticky ends complemented with the backbone pPAS-PT.

The pTHS-T7 RNAP* cassette was integrated into the chromosome of *S. albus*, and the pPAS-P_T7_ series plasmids carrying the T7 promoters were introduced into the resulting strain. The *kasO*p* promoter was inserted into pPAP-PT to generate the pPAS-*kasO*p* plasmid as a positive control. Quantitation of sfGFP expression was performed as described previously (Bai et al., 2015).

### Cloning and refactoring the *act* gene cluster

The *act* gene cluster from *S. coelicolor* M145 was cloned directly using a previously described method(Jiang et al., 2015). Briefly, mycelia of *S. coelicolor* M145 were collected after 2 days of cultivation. The preparation of the genomic DNA plugs was carried out according to the CHEF genomic DNA plug kit (Bio-Rad, Hercules, CA, USA). The DNA templates of sgRNA-actF and sgRNA-actR were generated by overlap extension of sgRNA-actF/sgRNA-actR and guide RNA-F plus guide RNA-R, respectively (Table S4). *In vitro* transcription of sgRNAs was performed using the HiScribe™ T7 Quick High Yield RNA Synthesis Kit (NEB). Then, the DNA plugs were digested with 500 ng Cas9 enzyme, 500 ng sgRNA-F and 500 ng sgRNA-R at 37°C for 2 hours. The digested DNA was then precipitated with ethanol and resuspended in 20 µL DNase-free water. The linearized capture vector pPAB-HR was constructed to introduce two ~30 bp overlaps with the corresponding ends of the *act* gene cluster fragment and digested with *Aar*I. Approximately 50 ng of the pPAB-HR backbone and 1 µg of digested genomic fragments were assembled using Gibson assembly and introduced into *E. coli* EPI300 by electroporation. The correct recombinant plasmid was verified by PCR using the PF-1 & PR-1, PF-2 & PR-2 primers (Table S4) (Figure S3). The recombinant plasmid was further confirmed by restriction enzyme digestion with I-*Sce*I (Figure S3).

For *act* gene cluster editing, four CRISPR target sequences were selected from three promoter regions (1, 2 and 3) in the *act* cluster, and one target sequence (4) for insertion an auxotrophic marker in the backbone of pPAB (Figure 5B). sgRNA-1, sgRNA-2, sgRNA-3, sgRNA-4were generated as described above. pPAB-act plasmid (10 µg) was digested with Cas9 guided by sgRNAs. After digestion, DNA was precipitated with ethanol and resuspended in 50µL water. Promoter cassettes

(Table S5) were synthesized by Generay Biotechnology. The yeast autotrophic marker (URA) and promoters were amplified by PCR using primers URA-F and URA-R (Table S4). Purified PCR products (150-300 ng) and the digested pPAB-act fragments (1 µg) were transformed into *<u>Saccharomyces</u> cerevisiae* VL6-48(Yamanaka et al., 2014) using the Frozen-EZ Yeast Transformation II Kit (Zymo Research). The correct promoter insertion was screened by PCR using primers upstream and downstream of the target site (Table S4). The correct plasmids were isolated and transferred into *S. albus* by conjugation.

## Compliance and ethics

*The author(s) declare that they have no conflict of interest*

## Acknowledgements

This work was supported by the National Key Research and Development Program of China [2020YFA0906900, 2018YFA0900700]; Natural Science Foundation of China [31500069], the Chinese Academy of Sciences [No. QYZDB-SSW-SMC050, No. No. XDB0480000 of the Strategic Priority Research Program], CAS Youth Interdisciplinary Team and the Shenzhen Science and Technology Innovation Committee [No. JCYJ20180507182241844, JCHZ20200005, DWKF20190009].

## Notes

### Competing Interest Statement

The authors have declared no competing interest.

## References

Aubry, C., Pernodet, J.L., and Lautru, S. (2019). Modular and Integrative Vectors for Synthetic Biology Applications in Streptomyces spp. Appl Environ Microbiol 85.

Bai, C., Zhang, Y., Zhao, X., Hu, Y., Xiang, S., Miao, J., Lou, C., and Zhang, L. (2015). Exploiting a precise design of universal synthetic modular regulatory elements to unlock the microbial natural products in Streptomyces. Proc Natl Acad Sci U S A 112, 12181–12186.

Bierman, M., Logan, R., O’Brien, K., Seno, E.T., Rao, R.N., and Schoner, B.E. (1992). Plasmid cloning vectors for the conjugal transfer of DNA from Escherichia coli to Streptomyces spp. Gene 116, 43–49.

Chater, K.F., and Wilde, L.C. (1976). Restriction of a bacteriophage of Streptomyces albus G involving endonuclease SalI. J Bacteriol 128, 644–650.

Demain, A.L. (2014). Importance of microbial natural products and the need to revitalize their discovery. J Ind Microbiol Biotechnol 41, 185–201.

Engler, C., Gruetzner, R., Kandzia, R., and Marillonnet, S. (2009). Golden gate shuffling: a one-pot DNA shuffling method based on type IIs restriction enzymes. PLoS One 4, e5553.

Flett, F., Mersinias, V., and Smith, C.P. (1997). High efficiency intergeneric conjugal transfer of plasmid DNA from Escherichia coli to methyl DNA-restricting streptomycetes. FEMS Microbiol Lett 155, 223–229.

Gibson, D.G., Young, L., Chuang, R.Y., Venter, J.C., Hutchison, C.A., 3rd, and Smith, H.O. (2009). Enzymatic assembly of DNA molecules up to several hundred kilobases. Nat Methods 6, 343–345.

Gomez-Escribano, J.P., and Bibb, M.J. (2011). Engineering Streptomyces coelicolor for heterologous expression of secondary metabolite gene clusters. Microb Biotechnol 4, 207–215.

Hopwood, D.A., Kieser, T., Wright, H.M., and Bibb, M.J. (1983). Plasmids, recombination and chromosome mapping in Streptomyces lividans 66. J Gen Microbiol 129, 2257–2269.

Jiang, W.J., Zhao, X.J., Gabrieli, T., Lou, C.B., Ebenstein, Y., and Zhu, T.F. (2015). Cas9-Assisted Targeting of CHromosome segments CATCH enables one-step targeted cloning of large gene clusters. Nature Communications 6.

Kang, H.S., Charlop-Powers, Z., and Brady, S.F. (2016). Multiplexed CRISPR/Cas9- and TAR-Mediated Promoter Engineering of Natural Product Biosynthetic Gene Clusters in Yeast. ACS Synth Biol 5, 1002–1010.

Kim, H., Ji, C.H., Je, H.W., Kim, J.P., and Kang, H.S. (2020). mpCRISTAR: Multiple Plasmid Approach for CRISPR/Cas9 and TAR-Mediated Multiplexed Refactoring of Natural Product Biosynthetic Gene Clusters. ACS Synth Biol 9, 175–180.

Kim, S.H., Lu, W., Ahmadi, M.K., Montiel, D., Ternei, M.A., and Brady, S.F. (2019). Atolypenes, Tricyclic Bacterial Sesterterpenes Discovered Using a Multiplexed In Vitro Cas9-TAR Gene Cluster Refactoring Approach. ACS Synth Biol 8, 109–118.

Lee, T.S., Krupa, R.A., Zhang, F., Hajimorad, M., Holtz, W.J., Prasad, N., Lee, S.K., and Keasling, J.D. (2011). BglBrick vectors and datasheets: A synthetic biology platform for gene expression. J Biol Eng 5, 12.

Liu, H., Jiang, H., Haltli, B., Kulowski, K., Muszynska, E., Feng, X., Summers, M., Young, M., Graziani, E., Koehn, F., Carter, G.T., and He, M. (2009). Rapid cloning and heterologous expression of the meridamycin biosynthetic gene cluster using a versatile Escherichia coli-streptomyces artificial chromosome vector, pSBAC. J Nat Prod 72, 389–395.

Liu, Z., Zhao, Y., Huang, C., and Luo, Y. (2021). Recent Advances in Silent Gene Cluster Activation in Streptomyces. Front Bioeng Biotechnol 9, 632230.

Miao, V., Coeffet-LeGal, M.F., Brian, P., Brost, R., Penn, J., Whiting, A., Martin, S., Ford, R., Parr, I., Bouchard, M., Silva, C.J., Wrigley, S.K., and Baltz, R.H. (2005). Daptomycin biosynthesis in Streptomyces roseosporus: cloning and analysis of the gene cluster and revision of peptide stereochemistry. Microbiology (Reading) 151, 1507–1523.

Ostrov, N., Landon, M., Guell, M., Kuznetsov, G., Teramoto, J., Cervantes, N., Zhou, M., Singh, K., Napolitano, M.G., Moosburner, M., Shrock, E., Pruitt, B.W., Conway, N., Goodman, D.B., Gardner, C.L., Tyree, G., Gonzales, A., Wanner, B.L., Norville, J.E., Lajoie, M.J., and Church, G.M. (2016). Design, synthesis, and testing toward a 57-codon genome. Science 353, 819–822.

Phelan, R.M., Sachs, D., Petkiewicz, S.J., Barajas, J.F., Blake-Hedges, J.M., Thompson, M.G., Reider Apel, A., Rasor, B.J., Katz, L., and Keasling, J.D. (2017). Development of Next Generation Synthetic Biology Tools for Use in Streptomyces venezuelae. ACS Synth Biol 6, 159–166.

Smolke, C.D. (2009). Building outside of the box: iGEM and the BioBricks Foundation. Nat Biotechnol 27, 1099–1102.

Sun, J.H., Kelemen, G.H., Fernandez-Abalos, J.M., and Bibb, M.J. (1999). Green fluorescent protein as a reporter for spatial and temporal gene expression in Streptomyces coelicolor A3(2). Microbiol-Uk 145, 2221–2227.

Wang, W., Zheng, G., and Lu, Y. (2021). Recent Advances in Strategies for the Cloning of Natural Product Biosynthetic Gene Clusters. Front Bioeng Biotechnol 9, 692797.

Yamanaka, K., Reynolds, K.A., Kersten, R.D., Ryan, K.S., Gonzalez, D.J., Nizet, V., Dorrestein, P.C., and Moore, B.S. (2014). Direct cloning and refactoring of a silent lipopeptide biosynthetic gene cluster yields the antibiotic taromycin A. Proc Natl Acad Sci U S A 111, 1957–1962.

Yang, K., Han, L., and Vining, L.C. (1995). Regulation of jadomycin B production in Streptomyces venezuelae ISP5230: involvement of a repressor gene, jadR2. J Bacteriol 177, 6111–6117.

Zhang, S., Zhao, X., Tao, Y., and Lou, C. (2015). A novel approach for metabolic pathway optimization: Oligo-linker mediated assembly (OLMA) method. J Biol Eng 9, 23.

Zhao, X., Wei, W., Zong, Y., Bai, C., Guo, X., Zhu, H., and Lou, C. (2022). Novel switchable ECF sigma factor transcription system for improving thaxtomin A production in Streptomyces. Synth Syst Biotechnol 7, 972–981.

Zhuo, J., Ma, B., Xu, J., Hu, W., Zhang, J., Tan, H., and Tian, Y. (2017). Reconstruction of a hybrid nucleoside antibiotic gene cluster based on scarless modification of large DNA fragments. Sci China Life Sci 60, 968–979.

Zong, Y., Zhang, H.M., Lyu, C., Ji, X., Hou, J., Guo, X., Ouyang, Q., and Lou, C. (2017). Insulated transcriptional elements enable precise design of genetic circuits. Nat Commun 8, 52.

