## Supplemental for "A flexible, modular and versatile functional part assembly toolkit for gene cluster engineering in *Streptomyces*"

### SUPPORTION INFORMATION

**Figure S1** Alternative *attP-attB* integration sites in four *Streptomyces* species.

**Figure S2** Schematic maps of the 10 basic vectors.

**Figure S3** Confirmation of the plasmid pPAB-act.

**Table S1** The sequences of plasmids.

**Table S2** The sequences of $P_{T7}$ promoter variants with corresponding sfGFP intensity.

**Table S3** The sequence of T7 RNAP*.

**Table S4** Primers used in this study.

**Table S5** The list of promoter cassettes and URA marker.

| Integrase | attP |
|---|---|
| ϕC31 | CCCCAACTGGGGTAACCTTTGAGTTCTCTCAGTTGGGGG |
| TG1 | TCCAGCCCAACAGTGTTAGTCTTTGCTCTTACCCAGTTGGGCGGGA |
| VWB | GACTCACTGAGGCTCATGATCGCTTTACGTTCTCTCCTAAAGCGG |

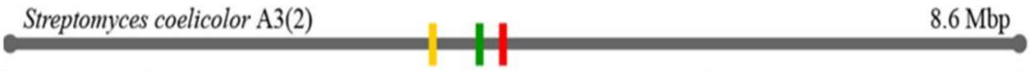


| Integrase | attB | Gene ID |
|---|---|---|
| ϕC31 | GTGCCAGGGCGTGCCCTTGGGCTCCCCGGGCGCG | SCO3798 |
| TG1 | TCCAGCCCAACAGTGTTAGTCTTTGCTCTTACCCAGTTGGGCGGGA | SCO3658 |
| VWB | CATCGGGTCGGCCGGCAAGACGTTCTCGTTCACCGGCTGGAAGGTC | SCOt27 |

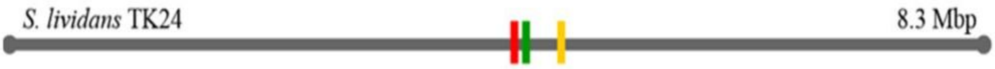


| Integrase | attB | Gene ID |
|---|---|---|
| ϕC31 | GTGCCAGGGCGTGCCCTTGGGCTCCCCGGGCGCG | SLIV_RS19325 |
| TG1 | TCCAGCCCAACAGTGTTAGTCTTTGCTCTCTTACCCAGTTGGGCGGGA | SLIV_RS20010 |
| VWB | CATCGGGTCGGCCGGCAAGACGTTCTCGTTCACCGGCTGGAAGGTC | SLIV_RS21445 |

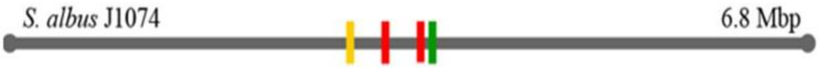


| Integrase | attB | Gene ID |
|---|---|---|
| ϕC31 | GGGTGCCAGGGAGAGCCGTACGTCTCGCCCTGGCACCC | Between XNR_2791 and XNR_2792 |
| | GTGCCAGGGCGTGCCCTTGGGCTCCCCGGGCGCG | XNR_3086 |
| TG1 | TCCAGCCCAACAGTGTTAGTCTTTGCTCTCTTACCCAGTTGGGCGGGA | XNR_3189 |
| VWB | CATCGGGTCGGCCGGCAAGACGTTCTCGTTCACCGGCTGGAAGGTC | XNR_2176 |

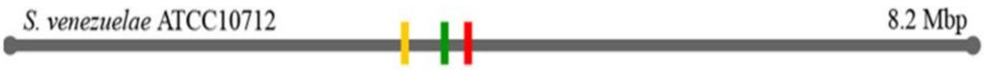


| Integrase | attB | Gene ID |
|---|---|---|
| ϕC31 | GTGCCAGGGCGTGCCCTTGGGCTCCCCGGGCGCG | SVEN_RS17625 |
| TG1 | TCCAGCCCAACAGTGTTAGTCTTTGCTCTCTTACCCAGTTGGGCGGGA | SVEN_RS16920 |
| VWB | CATCGGGTCGGCCGGCAAGACGTTCTCGTTCACCGGCTGGAAGGTC | SVEN_RS15185 |

**Figure S1** Alternative *attP*-*attB* integration sites in four *Streptomyces* chromosomes. The short lines with different colors indicate the locations of *attB* sites on chromosomes.

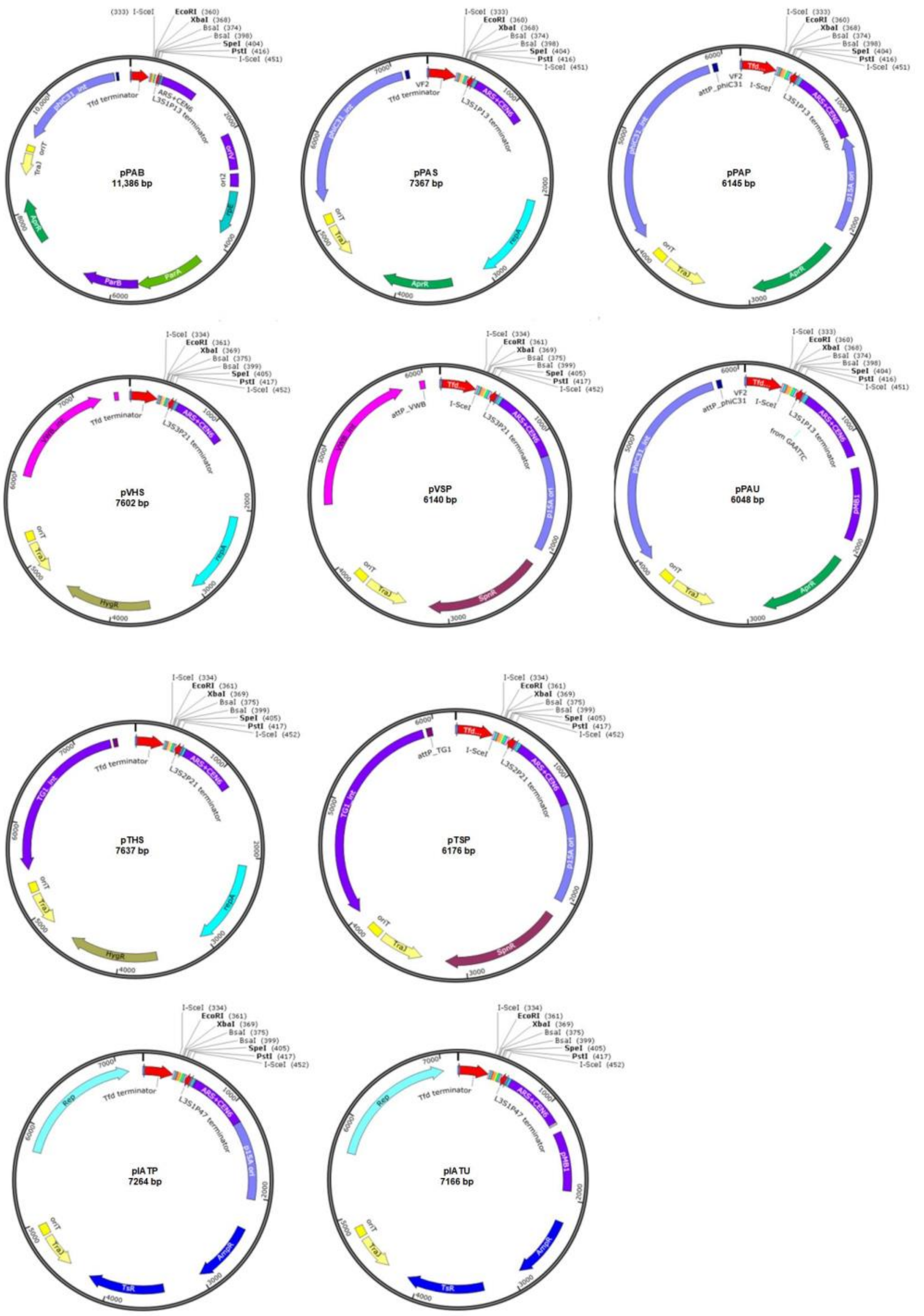


**Figure S2** Schematic maps of the 10 basic plasmids. Indicated sites for restriction enzymes are available for cloning DNA fragments using corresponding methods.

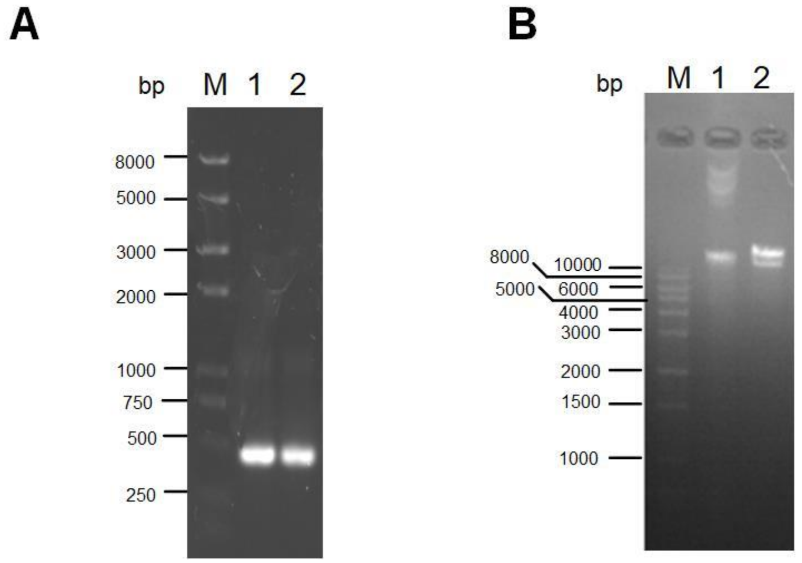


**Figure S3**. Confirmation of the plasmid pPAB-act. A. PCR verification of gene cluster were performed by PCR. M: Trans2K® Plus DNA Marker; 1: PCR product by PF1& PR1; 2: PCR product by PF2& PR2. B.Restriction enzyme digestion verification. M:1kb Plus DNA Ladder; 1: plamid pPAB-act; 2 pPAB-act was digested by I-*Sce*I.

**Table S1** Plasmid sequences

| Name | Sequence |
|---|---|
| pPAB | tgccacctgacgtctaagaagatcccgcaaaagcggcctttgactccctgcaagcctcagcgaccgaatatatcggttatgcgtgggcga<br>tggttgttgtcattgtcggcgcaactatcggtatcaagctgtttaagaaattcacctcgaaagcaagctgataaaccgatacaattaaaggct<br>ccttttggagccttttttttggagattttcaacgtgaaaaaattattattcgcaattcctttagttgttcctttctattctcactccgctgaaactgttg<br>aaagttgtttagcaaaacctcatacagaaaattcatattaccctgttatccctactccgagacagtcagagggtagaattccttctagagcctc<br>agagacccggccgggtctcaggagtactagttcctgcagcactcgttgctttatcggtatattaccctgttatccctaattttgttatcaataaa<br>aaaggcccccccgttagggaggccttattgttcgtcgctcactcaaaggcggtaatgacgctcagtggaacgaaaactcacgttaagggta<br>aaactgtattataagtaaatgcatgtatactaaactcacaaattagagcttcaatttaattatatcagttattacccataacttcgtatagcatacat<br>tatacgaagttatcccgggtaccgagctcgattcgtaacttacacgcgcctcgtatcttttaatgatggaataatttgggaatttactctgtgttt<br>atttatttttatgttttgtatttggattttagaaagtaaataaagaaggtagaagagttacggaatgaagaaaaaaaaataaacaaaggtttaaaa<br>aatttcaacaaaaagcgtactttacatatatatttattagacaagaaaagcagattaaatagatatacattcgattaacgataagtaaaatgtaa<br>aatcacaggattttcgtgtgtggtcttctacacagacaagatgaaacaattcggcattaatacctgagagcaggaagagcaagataaaagg<br>tagtatttgttggcgatccccctagagtcttttacatcttcggaaaacaaaaactattttttctttaatttctttttttactttctatttttaatttatatattt<br>atattaaaaaatttaaattataattatttttatagcacgtgagatcaacgtctcattttcgccaaaagttggcccagggcttcccggtatcaacag<br>ggacaccaggatttatttattctgcgaagtgatcttccgtcacaggtatttattcgcgataagctcatggagcggcgtaaccgtcgcacagga<br>aggacagagaaagcgcggatctgggaagtgacggacagaacggtcaggacctggattggggaggcggttgccgccgctgctgctga<br>cggtgtgacgttctctgttccggtcacaccacatacgttccgccattcctatgcgatgcacatgctgtatgccggtataccgctgaaagttctg<br>caaagcctgatgggacataagtccatcagttcaacggaagtctacacgaaggtttttgcgctggatgtggctgcccggcaccgggtgcag<br>tttgcgatgccggagtctgatgcggttgcgatgctgaaacaattatcctgagaataaatgccttggcctttatatggaaatgtggaactgagt<br>ggatatgctgtttttgtctgttaaacagagaagctggctgttatccactgagaagcgaacgaaacagtcgggaaaatctcccattatcgtaga<br>gatccgcattattaatctcaggagcctgtgtagcgtttataggaagtagtgttctgtcatgatgcctgcaagcggtaacgaaaacgatttgaat<br>atgccttcaggaacaatagaaatcttcgtgcggtgttacgttgaagtggagcggattatgtcagcaatggacagaacaacctaatgaacac<br>agaaccatgatgtggtctgtccttttacagccagtagtgctcgccgcagtcgagcgacagggcgaagccctcgagctggttgccctcgcc<br>gctgggctggcggccgtctatggccctgcaaacgcgccagaaacgccgtcgaagccgtgtgcgagacaccgcggccggccgccggc<br>gttgtggatacctcgcggaaaacttggccctcactgacagatgaggggcggacgttgacacttgaggggccgactcacccggcgcggc<br>gttgacagatgaggggcaggctcgatttcggccggcgacgtggagctggccagcctcgcaaatcggcgaaaacgcctgattttacgcg<br>agtttcccacagatgatgtggacaagcctggggataagtgccctgcggtattgacacttgaggggcgcgactactgacagatgaggggc<br>gcgatccttgacacttgaggggcagagtgctgacagatgaggggcgcacctattgacatttgaggggctgtccacaggcagaaaatcca<br>gcatttgcaagggtttccgcccgtttttcggccaccgctaacctgtcttttaacctgcttttaaaccaatatttataaaccttgtttttaaccaggg<br>ctgcgccctgtgcgcgtgaccgcgcacgccgaaggggggtgcccccccttctcgaaccctcccggtcgagtgagcgaggaagcacca<br>gggaacagcacttatatattctgcttacacacgatgcctgaaaaaacttcccttggggttatccacttatccacggggatatttttataattatttt<br>ttttatagtttttagatcttctttttttagagcgccttgtaggcctttatccatgctggttcatgagaaggtgttgtgacaaattgccctttcagtgtga<br>caaatcaccctcaaatgacagtcctgtctgtgacaaattgcccttaaccctgtgacaaattgccctcagaagaagctgtttttttcacaaagttat<br>ccctgcttattgactctttttttatttagtgtgacaatctaaaaacttgtcacacttcacatggatctgtcatggcggaaacagcggttatcaatcac<br>aagaaacgtaaaaatagcccgcgaatcgtccagtcaaacgacctcactgaggcggcatatagtctctcccgggatcaaaaacgtatgctg<br>tatctgttcgttgaccagatcagaaaatctgatggcaccctacaggaacatgacggtatctgcgagatccatgttgctaaatatgctgaaata<br>ttcggattgacctctgcggaagccagtaaggatatacggcaggcattgaagagtttcgcggggaaggaagtggtttttttatcgccctgaag<br>aggatgccggcgatgaaaaaggctatgaatcttttccttggtttatcaaacgtgcgcacagtccatccagagggctttacagtgtacatatca<br>acccatatctcattcccttctttatcgggttacagaaccggtttacgcagtttcggcttagtgaaacaaaagaaatcaccaatccgtatgccat<br>gcgtttatacgaatccctgtgtcagtatcgtaagccggatggctcaggcatcgtctctctgaaaatcgactggatcatagagcgttaccagct<br>gcctcaaagttaccagcgtatgcctgacttccgccgccgcttcctccaggtctgtgttaatgagatcaacagcagaactccaatgcgcctct<br>catacattgagaaaaagaaaggccgccagacgactcatatcgtattttccttccgcgatatcacttccatgacgacaggatagtctgagggt<br>tatctgtcacagatttgagggtggttcgtcacatttgttctgacctactgagggtaatttgtcacagttttgctgtttccttcagcctgcatggattt<br>tctcatactttttgaactgtaatttttaaggaagccaaatttgagggcagtttgtcacagttgatttccttctctttcccttcgtcatgtgacctgata<br>tcgggggttagttcgtcatcattgatgagggttgattatcacagtttattactctgaattggctatccgcgtgtgtacctctacctggagtttttcc<br>cacggtggatatttcttcttgcgctgagcgtaagagctatctgacagaacagttcttctttgcttcctcgccagttcgctcgctatgctcggtta<br>cacggctgcggcgagcgctagtgataataagtgactgaggtatgtgctcttcttatctccttttgtagtgttgctcttattttaaacaactttgcg<br>gttttttgatgactttgcgattttgttgttgctttgcagtaaattgcaagatttaataaaaaaacgcaaagcaatgattaaaggatgttcagaatga<br>aactcatggaaacacttaaccagtgcataaacgctggtcatgaaatgacgaaggctatcgccattgcacagtttaatgatgacagcccgga<br>agcgaggaaaataacccggcgctggagaataggtgaagcagcggatttagttggggtttcttctcaggctatcagagatgccgagaaag<br>cagggcgactaccgcacccggatatggaaattcgaggacgggttgagcaacgtgttggttatacaattgaacaaattaatcatatgcgtga<br>tgtgtttggtacgcgattgcgacgtgctgaagacgtatttccaccggtgatcggggttgctgcccataaaggtggcgtttacaaaacctcag<br>tttctgttcatcttgctcaggatctggctctgaaggggctacgtgttttgctcgtggaaggtaacgacccccagggaacagcctcaatgtatc<br>acggatgggtaccagatcttcatattcatgcagaagacactctcctgcctttctatcttggggaaaaggacgatgtcacttatgcaataaagc<br>ccacttgctggccggggcttgacattattccttcctgtctggctctgcaccgtattgaaactgagttaatgggcaaatttgatgaaggtaaact<br>gcccaccgatccacacctgatgctccgactggccattgaaactgttgctcatgactatgatgtcatagttattgacagcgcgcctaacctgg<br>gtatcggcacgattaatgtcgtatgtgctgctgatgtgctgattgttcccacgcctgctgagttgtttgactacacctccgcactccagtttttcg<br>atatgcttcgtgatctgctcaagaacgttgatcttaaagggttcgagcctgatgtacgtattttgcttaccaaatacagcaatagtaatggctct<br>cagtccccgtggatggaggagcaaattcgggatgcctggggaagcatggttctaaaaaatgttgtacgtgaaacggatgaagttggtaaa<br>ggtcagatccggatgagaactgtttttgaacaggccattgatcaacgctcttcaactggtgcctggagaaatgctctttctatttgggaacctg<br>tctgcaatgaaattttcgatcgtctgattaaaccacgctgggagattagataatgaagcgtgcgcctgttattccaaaacatacgctcaatact<br>caaccggttgaagatacttcgttatcgacaccagctgccccgatggtggattcgttaattgcgcgcgtaggagtaatggctcgcggtaatg<br>ccattactttgcctgtatgtggtcgggatgtgaagtttactcttgaagtgctccggggtgatagtgttgagaagacctctcgggtatggtcagg |

| | |
|---|---|
| | taatgaacgtgaccaggagctgcttactgaggacgcactggatgatctcatcccttcttttctactgactggtcaacagacaccggcgttcg<br>gtcgaagagtatctggtgtcatagaaattgccgatgggagtcgccgtcgtaaagctgctgcacttaccgaaagtgattatcgtgttctggttg<br>gcgagctggatgatgagcagatggctgcattatccagattgggtaacgattatcgcccaacaagtgcttatgaacgtggtcagcgttatgc<br>aagccgattgcagaatgaatttgctggaaatatttctgcgctggctgatgcggaaaatatttcacgtaagattattacccgctgtatcaacacc<br>gccaaattgcctaaatcagttgttgctctttttctcaccccggtgaactatctgcccggtcaggtgatgcacttcaaaaagcctttacagataa<br>agaggaattacttaagcagcaggcatctaaccttcatgagcagaaaaaagctggggtgatatttgaagctgaagaagttatcactcttttaac<br>ttctgtgcttaaaacgtcatctgcatcaagaacaagtttaagctcacgacatcagtttgctcctggagcgacagtattgtataagggcgataa<br>aatggtgcttaacctggacaggtcccgtgttccaactgagtgtatagagaaaattgaggccattcttaaggaacttgaaaagccagcaccct<br>gatgcgaccacgttttagtctacgtttatctgtctttacttaatgtcctttgttacaggccagaaagcataactggcctgaatattctctctgggcc<br>cactgttccacttgtatcgtcggtctgataatcagactgggaccacggtcccactcgtatcgtcggtctgattattagtctgggaccacggtc<br>ccactcgtatcgtcggtctgattattagtctgggaccacggtcccactcgtatcgtcggtctgataatcagactgggaccacggtcccactc<br>gtatcgtcggtctgattattagtctgggaccatggtcccactcgtatcgtcggtctgattattagtctgggaccacggtcccactcgtatcgtc<br>ggtctgattattagtctggaaccacggtcccactcgtatcgtcggtctgattattagtctgggaccacggtcccactcgtatcgtcggtctgat<br>tattagtctgggaccacgatcccactcgtgttgtcggtctgattatcggtctgggaccacggtcccacttgtattgtcgatcagactatcagcg<br>tgagactacgattccatcaatgcctgtcaagggcaagtattgacatgtcgtcgtaacctgtagaacggagtaacctcggtgtgcggttgtat<br>gcctgctgtggattgctgctgtgtcctgcttatccacaacattttgcgcacggttatgtggacaaaatacctgattttggtcatgagattatcaa<br>aaaggatcttcacctagatccttttggttcatgtgcagctccatcagcaaaaggggatgataagtttatcaccaccgactatttgcaacagtgc<br>cgttgatcgtgctatgatcgactgatgtcatcagcggtggagtgcaatgtcgtgcaatacgaatggcgaaaagccgagctcatcggtcagc<br>ttctcaaccttggggttaccccggcggtgtgctgctggtccacagctccttccgtagcgtccggcccctcgaagatgggccacttggact<br>gatcgaggccctgcgtgctgcgctgggtccgggagggacgctcgtcatgccctcgtggtcaggtctggacgacgagccgttcgatcctg<br>ccacgtcgcccgttacaccggaccttggagttgtctctgacacattctggcgcctgccaaatgtaaagcgcagcgcccatccatttgccttt<br>gcggcagcggggccacaggcagagcagatcatctctgatccattgcccctgccacctcactcgcctgcaagcccggtcgcccgtgtcc<br>atgaactcgatgggcaggtacttctcctcggcgtgggacacgatgccaacacgacgctgcatcttgccgagttgatggcaaaggttccct<br>atggggtgccgagacactgcaccattcttcaggatggcaagttggtacgcgtcgattatctcgagaatgaccactgctgtgagcgctttgc<br>cttggcggacaggtggctcaaggagaagagccttcagaaggaaggtccagtcggtcatgcctttgctcggttgatccgctcccgcgacat<br>tgtggcgacagccctgggtcaactgggccgagatccgttgatcttcctgcatccgccagaggcgggatgcgaagaatgcgatgccgctc<br>gccagtcgattggctgagctcatgagcggagaacgagatgacgttggaggggcaaggtcgcgctgattgctggggcaacacgtggag<br>cggatcggggattgtctttcttcagctcgctgatgatatgctgacgctcaatgccgtttggcctccgactaacgaaaatcccgcatttggacg<br>gctgatccgattggcacggcggacggcgaatggcggagcagacgctcgtccgggggcaatgagatatgaaaaagcctgaactcaccg<br>cgacgtatcgggccctggccagctagctagagtcgacctgggtccccggggatcggtcttgccttgctcgtcggtgatgtacttcaccag<br>ctccgcgaagtcgctcttcttgatggagcgcatggggacgtgcttggcaatcacgcgcacccccggccgttttagcggctaaaaaagtc<br>atggctctgccctcgggcggaccacgcccatcatgaccttgccaagctcgtcctgcttctcttcgatcttcgccagcagggcgaggatcgt<br>ggcatcaccgaaccgcgccgtgcgcgggtcgtcggtgagccagagtttcagcaggccgcccaggcggcccaggtcgccattgatgcg<br>ggccagctcgcggacgtgctcatagtccacgacgcccgtgattttgtagccctggccgacggccagcaggtaggccgacaggctcatg<br>ccggccgccgccgccttttcctcaatcgctcttcgttcgtctggaaggcagtacaccttgataggtgggctgcccttcctggttggcttggttt<br>catcagccatccgcttgccctcatctgttacgccggcggtagccggccagcctcgcagagcaggattcccgttgagcaccgccaggtgc<br>gaataagggacagtgaagaaggaacacccgctcgcgggtgggcctacttcacctatcctgcccggctgacgccgttggatacaccaag<br>gaaagtctacacgaaccctttggcaaaatcctgtatatcgtgcgaaaaaggatggatataccgaaaaaatcgctataatgaccccgaagca<br>gggttatgcagcggaaaagatccgtcgacctgctacgccgctacgtcttccgtgccgtcctgggcgtcgtcttcgtcgtcgtcggtcggcg<br>gcttcgcccacgtgatcgaagcgcgcttctcgatgggcgttccctgccccctgcccgtagtcgacttcgtgacaacgatcttgtctacgaa<br>gagcccgacgaacacgcgcttgtcgtctactgacgcgcgcccccaccacgacttagggccggtcgggtcagcgtcggcgtcttcgggg<br>aaccattggtcaaggggaagcttcggggcttcggcggcttcaagttcggcaagccgctcttccgccccttgctgccggagcgtcagcgct<br>gcctgttgcttccggaagtgcttcctgccaacgggtccgtcgtacgcgcctgccgcgcggtcttcgtacagctcttcaagggcgttcaggg<br>cgtcggcgcgctccgcaacaaggttcgcccgttcgccgctcttctcaggcgcctcagtgagcttgccgaagcgtcgggcggcttcccac<br>agaagcgccaacgtctcttcgtcgccttcggcgtgcctgatcttgttgaagatgcgttccgcaacgaacttgtcgagtgccgccatgctgac<br>gttgcacgtgccttcgtgctgcccaggtgcggacgggtcgaccaccttccggcgacggcagcggtaagagtccttgatcgattcttcccc<br>gcgcttcgaagtcatgacggcgccacactcgcagtacagcttgtccatggcggacagaatggcttgcccccgggaaagccccttgccgc<br>gccccctgccgtccaaccacgcctgaagctcataccactcagcgggctcgatgatcggtccgcaatcaagctcgaccggccggagcgt<br>gatcgggtcgcgctgaatgcggtaaccctcaatcttcgtggtcggcgtgccgtccggcttcttcttgtagatcacctcagcggcgaagccc<br>gcaatacgcgggtcccgaaggattcgcataacggttgccgggtcccaggcgcttgaagcggtcttcttcccaatcgtctcgccccgggtc<br>ggcacggcgtcagcgtccatgcgcttacaaagccccgtgatgctgcccgggtgaatggcggcttgactgcccggcttgaagggaaggt<br>gtttgtgcgtcttgatctcacgccaccaccaccggattacgtcgggctcgaactcgaagggtccggtaaggggagtggtcgagtgcgcaa<br>gcttgttgatgacgacattgaccattcggccgttgcgcgtgatctccttcgtctccgaaacaagctcgaagccgtaaggcgccttcccgcc<br>gacgtacccgcccaattcgcgctgaaggttcttcgtgtcgagaatcttcgccgacttcagcgaagattctttgtgcgacgcgtcgagccgc<br>ataatcaggtgaatcaggtccatgacgtttccctgccggaagacgccttcctgagtggaaacaatcgtcacgcccagggcgagcaattcc<br>gagacaatcggaatcgcgtccatgaccttcaggcgcgagaagcgcgacacgtcatagacaatgatcatgttgagccgcccggcgcggc<br>attcgttcaggatgcgttcgaactccgggcgctccgccgtcccgaacgccgacgtgcccggcgcttcgctgaaatgcccgacgaacctg<br>aaccggcccccgtcgcgctcgacttcgcgctgaaggtcggccgccttgtcttcgttggcgctacgctgtgtcgctgggcttgctgcgctcg<br>aactctcgcgctcgcgcgactgacggtcgtaagcacccgcgtacgtgtccaccccggtcacaaccccttgtgtcatgtcggcgaccctac<br>gcccccaactgagagaactcaaaggttaccccagttggggcactactcccgaaaaccgcttctgacctgggaaaacgtgaagccccgg<br>ggcatccgctgagggttgccgccggggcttcggtgtgtccgtcagtacgggccatagaggggcgtcgtgtaacccacccagggcgatt<br>gctccggcacggggaagccgcgccacgccttcgggacgtctggaatcgctagagcttgcatgc |

| | |
|---|---|
| pPAS | tgccacctgacgtctaagaagatcccgcaaaagcggcctttgactccctgcaagcctcagcgaccgaatatatcggttatgcgtgggcga<br>tggttgttgtcattgtcggcgcaactatcggtatcaagctgtttaagaaattcacctcgaaagcaagctgataaaccgatacaattaaaggct<br>cctttggagcctttttttttggagattttcaacgtgaaaaaattattattcgcaattcctttagttgttcctttctattctcactccgctgaaactgttg<br>aaagttgtttagcaaaacctcatacagaaaattcatattaccctgttatccctactccgagacagtcagagggtagaattccttctagagcctc<br>agagacccggccgggtctcaggagtactagttcctgcagcactcgttgctttatcggtatattaccctgttatccctaattttgttatcaataaa<br>aaaggcccccccgttagggaggccttattgttcgtcgctcactcaaaggcggtaatgacgctcagtggaacgaaaactcacgttaagggta<br>aaactgtattataagtaaatgcatgtatactaaactcacaaattagagcttcaatttaattatatcagttattacccataacttcgtatagcatacat<br>tatacgaagttatcccgggtaccgagctcgattcgtaacttacacgcgcctcgtatcttttaatgatggaataatttgggaatttactctgtgttt<br>atttatttttatgttttgtatttggattttagaaagtaaataaagaaggtagaagagttacggaatgaagaaaaaaaaaataaacaaaggtttaaaa<br>aatttcaacaaaaagcgtactttacatatatatttattagacaagaaaagcagattaaatagatatacattcgattaacgataagtaaaatgtaa<br>aatcacaggattttcgtgtgtggtcttctacacagacaagatgaaacaattcggcattaatacctgagagcaggaagagcaagataaaagg<br>tagtatttgttggcgatccccctagagtcttttacatcttcggaaaacaaaaactattttttctttaatttctttttttactttctatttttaatttatatattt<br>atattaaaaaatttaaattataattatttttatagcacgtgagctagagctgtcagaccaagtttacgagctcgcttggactcctgttgatagatc<br>cagtaatgacctcagaactccatctggatttgttcagaacgctcggttgccgccgggcgttttttattggtgagaatccaagcactagggaca<br>gtaagacgggtaagcctgttgatgataccgctgccttactgggtgcattagccagtctgaatgacctgtcacgggataatccgaagtggtc<br>agactggaaaatcagagggcaggaactgctgaacagcaaaaagtcagatagcaccacatagcagacccgccataaaacgccctgaga<br>agcccgtgacgggcttttcttgtattatgggtagtttccttgcatgaatccataaaaggcgcctgtagtgccatttacccccattcactgccag<br>agccgtgagcgcagcgaactgaatgtcacgaaaaagacagcgactcaggtgcctgatggtcggagacaaaaggaatattcagcgattt<br>gcccgagcttgcgagggtgctacttaagcctttagggttttaaggtctgttttgtagaggagcaaacagcgtttgcgacatccttttgtaatact<br>gcggaactgactaaagtagtgagttatacacagggctgggatctattctttttatctttttttattctttctttattctataaattataaccacttgaat<br>ataaacaaaaaaaacacacaaaggtctagcggaatttacagagggtctagcagaatttacaagttttccagcaaaggtctagcagaatttac<br>agatacccacaactcaaaggaaaaggacatgtaattatcattgactagcccatctcaattggtatagtgattaaaatcacctagaccaattga<br>gatgtatgtctgaattagttgttttcaaagcaaatgaactagcgattagtcgctatgacttaacggagcatgaaaccaagctaattttatgctgt<br>gtggcactactcaaccccacgattgaaaaccctacaaggaaagaacggacggtatcgttcacttataaccaatacgctcagatgatgaac<br>atcagtagggaaaatgcttatggtgtattagctaaagcaaccagagagctgatgacgagaactgtggaaatcaggaatcctttggttaaag<br>gctttgagattttccagtggacaaactatgccaagttctcaagcgaaaaattagaattagtttttagtgaagagatattgccttatcttttccagtt<br>aaaaaaattcataaaatataatctggaacatgttaagtcttttgaaaacaaatactctatgaggatttatgagtggttattaaaagaactaacac<br>aaaagaaaactcacaaggcaaatatagagattagccttgatgaatttaagttcatgttaatgcttgaaaataactaccatgagtttaaaaggct<br>taaccaatgggttttgaaaccaataagtaaagatttaaacacttacagcaatatgaaattggtggttgataagcgaggccgcccgactgata<br>cgttgattttccaagttgaactagatagacaaatggatctcgtaaccgaacttgagaacaaccagataaaaatgaatggtgacaaaatacca<br>acaaccattacatcagattcctacctacgtaacggactaagaaaaacactacacgatgctttaactgcaaaaattcagctcaccagttttgag<br>gcaaaatttttgagtgacatgcaaagtaagcatgatctcaatggttcgttctcatggctcacgcaaaaacaacgaaccacactagagaacat<br>actggctaaatacggaaggatctgaggttcttatggctcttgtatctatcagtgaagcatcaagactaacaaacaaaagtagaacaactgttc<br>accgttagatatcaaagggaaaactgtccatatgcacagatgaaaacggtgtaaaaaagatagatacatcagagcttttacgagtttttggtg<br>catttaaagctgttcaccatgaacagatcgacaatgtaactactagagattttggtcatgagattatcaaaaaggatcttcacctagatcctttt<br>ggttcatgtgcagctccatcagcaaaaggggatgataagtttatcaccaccgactatttgcaacagtgccgttgatcgtgctatgatcgactg<br>atgtcatcagcggtggagtgcaatgtcgtgcaatacgaatggcgaaaagccgagctcatcggtcagcttctcaaccttggggttacccccc<br>ggcggtgtgctgctggtccacagctccttccgtagcgtccggcccctcgaagatgggccacttggactgatcgaggccctgcgtgctgc<br>gctgggtccgggagggacgctcgtcatgccctcgtggtcaggtctggacgacgagccgttcgatcctgccacgtcgcccgttacaccgg<br>accttggagttgtctctgacacattctggcgcctgccaaatgtaaagcgcagcgcccatccatttgcctttgcggcagcggggccacaggc<br>agagcagatcatctctgatccattgcccctgccacctcactcgcctgcaagcccggtcgcccgtgtccatgaactcgatgggcaggtactt<br>ctcctcggcgtgggacacgatgccaacacgacgctgcatcttgccgagttgatggcaaaggttccctatggggtgccgagacactgcac<br>cattcttcaggatggcaagttggtacgcgtcgattatctcgagaatgaccactgctgtgagcgctttgccttggcggacaggtggctcaagg<br>agaagagccttcagaaggaaggtccagtcggtcatgcctttgctcggttgatccgctcccgcgacattgtggcgacagccctgggtcaac<br>tgggccgagatccgttgatcttcctgcatccgccagaggcgggatgcgaagaatgcgatgccgctcgccagtcgattggctgagctcatg<br>agcggagaacgagatgacgttggaggggcaaggtcgcgctgattgctggggcaacacgtggagcggatcggggattgtctttcttcag<br>ctcgctgatgatatgctgacgctcaatgccgtttggcctccgactaacgaaaatcccgcatttggacggctgatccgattggcacggcgga<br>cggcgaatggcggagcagacgctcgtccgggggcaatgagatatgaaaaagcctgaactcaccgcgacgtatcgggccctggccagc<br>tagctagagtcgacctgggtccccggggatcggtcttgccttgctcgtcggtgatgtacttcaccagctccgcgaagtcgctcttcttgatg<br>gagcgcatggggacgtgcttggcaatcacgcgcacccccggccgttttagcggctaaaaaagtcatggctctgccctcgggcggacc<br>acgcccatcatgaccttgccaagctcgtcctgcttctcttcgatcttcgccagcagggcgaggatcgtggcatcaccgaaccgcgccgtg<br>cgcgggtcgtcggtgagccagagtttcagcaggccgcccaggcggcccaggtcgccattgatgcgggccagctcgcggacgtgctca<br>tagtccacgacgcccgtgattttgtagccctggccgacggccagcaggtaggccgacaggctcatgccggccgccgccgccttttcctc<br>aatcgctcttcgttcgtctggaaggcagtacaccttgataggtgggctgcccttcctggttggcttggtttcatcagccatccgcttgccctca<br>tctgttacgccggcggtagccggccagcctcgcagagcaggattcccgttgagcaccgccaggtgcgaataagggacagtgaagaag<br>gaacacccgctcgcgggtgggcctacttcacctatcctgcccggctgacgccgttggatacaccaaggaaagtctacacgaaccctttgg<br>caaaatcctgtatatcgtgcgaaaaaggatggatataccgaaaaaatcgctataatgaccccgaagcagggttatgcagcggaaaagatc<br>cgtcgacctgctacgccgctacgtcttccgtgccgtcctgggcgtcgtcttcgtcgtcgtcggtcggcggcttcgcccacgtgatcgaagc<br>gcgcttctcgatgggcgttccctgccccctgcccgtagtcgacttcgtgacaacgatcttgtctacgaagagcccgacgaacacgcgcttg<br>tcgtctactgacgcgcgcccccaccacgacttagggccggtcgggtcagcgtcggcgtcttcggggaaccattggtcaaggggaagctt<br>cggggcttcggcggcttcaagttcggcaagccgctcttccgccccttgctgccggagcgtcagcgctgcctgttgcttccggaagtgcttc<br>ctgccaacgggtccgtcgtacgcgcctgccgcgcggtcttcgtacagctcttcaagggcgttcagggcgtcggcgcgctccgcaacaag<br>gttcgcccgttcgccgctcttctcaggcgcctcagtgagcttgccgaagcgtcgggcggcttcccacagaagcgccaacgtctcttcgtc<br>gccttcggcgtgcctgatcttgttgaagatgcgttccgcaacgaacttgtcgagtgccgccatgctgacgttgcacgtgccttcgtgctgcc<br>caggtgcggacgggtcgaccaccttccggcgacggcagcggtaagagtccttgatcgattcttccccgcgcttcgaagtcatgacggcg<br>ccacactcgcagtacagcttgtccatggcggacagaatggcttgcccccgggaaagccccttgccgcgccccctgccgtccaaccacg<br>cctgaagctcataccactcagcgggctcgatgatcggtccgcaatcaagctcgaccggccggagcgtgatcgggtcgcgctgaatgcg |

| | |
|---|---|
| | gtaaccctcaatcttcgtggtcggcgtgccgtccggcttcttcttgtagatcacctcagcggcgaagcccgcaatacgcgggtcccgaag<br>gattcgcataacggttgccgggtcccaggcgcttgaagcggtcttcttcccaatcgtctcgccccgggtcggcacggcgtcagcgtccat<br>gcgcttacaaagccccgtgatgctgcccgggtgaatggcggcttgactgcccggcttgaagggaaggtgtttgtgcgtcttgatctcacgc<br>caccaccaccggattacgtcgggctcgaactcgaagggtccggtaaggggagtggtcgagtgcgcaagcttgttgatgacgacattgac<br>cattcggccgttgcgcgtgatctccttcgtctccgaaacaagctcgaagccgtaaggcgccttcccgccgacgtacccgcccaattcgcg<br>ctgaaggttcttcgtgtcgagaatcttcgccgacttcagcgaagattctttgtgcgacgcgtcgagccgcataatcaggtgaatcaggtcca<br>tgacgtttccctgccggaagacgccttcctgagtggaaacaatcgtcacgcccagggcgagcaattccgagacaatcggaatcgcgtcc<br>atgaccttcaggcgcgagaagcgcgacacgtcatagacaatgatcatgttgagccgcccggcgcggcattcgttcaggatgcgttcgaa<br>ctccgggcgctccgccgtcccgaacgccgacgtgcccggcgcttcgctgaaatgcccgacgaacctgaaccggcccccgtcgcgctc<br>gacttcgcgctgaaggtcggccgccttgtcttcgttggcgctacgctgtgtcgctgggcttgctgcgctcgaactctcgcgctcgcgcgac<br>tgacggtcgtaagcacccgcgtacgtgtccaccccggtcacaaccccttgtgtcatgtcggcgaccctacgcccccaactgagagaactc<br>aaaggttaccccagttggggcactactcccgaaaaccgcttctgacctgggaaaacgtgaagccccggggcatccgctgagggttgcc<br>gccggggcttcggtgtgtccgtcagtacgggccatagaggggcgtcgtgtaacccacccagggcgattgctccggcacggggaagcc<br>gcgccacgccttcgggacgtctggaatcgctagagcttgcatgc |
| pPAP | tgccacctgacgtctaagaagatcccgcaaaagcggcctttgactccctgcaagcctcagcgaccgaatatatcggttatgcgtgggcga<br>tggttgttgtcattgtcggcgcaactatcggtatcaagctgtttaagaaattcacctcgaaagcaagctgataaaccgatacaattaaaggct<br>ccttttggagcctttttttttggagattttcaacgtgaaaaaattattattcgcaattcctttagttgttcctttctattctcactccgctgaaactgttg<br>aaagttgtttagcaaaacctcatacagaaaattcatattaccctgttatccctactccgagacagtcagagggtagaattccttctagagcctc<br>agagacccggccgggtctcaggagtactagttcctgcagcactcgttgctttatcggtatattaccctgttatccctaattttgttatcaataaa<br>aaaggcccccccgttagggaggccttattgttcgtcgctcactcaaaggcggtaatgacgctcagtggaacgaaaactcacgttaagggta<br>aaactgtattataagtaaatgcatgtatactaaactcacaaattagagcttcaatttaattatatcagttattacccataacttcgtatagcatacat<br>tatacgaagttatcccgggtaccgagctcgattcgtaacttacacgcgcctcgtatcttttaatgatggaataatttgggaatttactctgtgttt<br>atttatttttatgttttgtatttggattttagaaagtaaataaagaaggtagaagagttacggaatgaagaaaaaaaaaataaacaaaggtttaaaa<br>aatttcaacaaaaagcgtactttacatatatatttattagacaagaaaagcagattaaatagatatacattcgattaacgataagtaaaatgtaa<br>aatcacaggattttcgtgtgtggtcttctacacagacaagatgaaacaattcggcattaatacctgagagcaggaagagcaagataaaagg<br>tagtatttgttggcgatccccctagagtcttttacatcttcggaaaacaaaaactatttttttctttaatttctttttttactttctatttttaatttatatattt<br>atattaaaaaatttaaattataattatttttatagcacgtgagcgctagcggagtgtatactggcttactatgttggcactgatgagggtgtcagt<br>gaagtgcttcatgtggcaggagaaaaaaggctgcaccggtgcgtcagcagaatatgtgatacaggatatattccgcttcctcgctcactga<br>ctcgctacgctcggtcgttcgactgcggcgagcggaaatggcttacgaacggggcggagatttcctggaagatgccaggaagatactta<br>acagggaagtgagagggccgcggcaaagccgtttttccataggctccgcccccctgacaagcatcacgaaatctgacgctcaaatcagt<br>ggtggcgaaacccgacaggactataaagataccaggcgtttccccctggcggctccctcgtgcgctctcctgttcctgcctttcggtttaccg<br>gtgtcattccgctgttatggccgcgtttgtctcattccacgcctgacactcagttccgggtaggcagttcgctccaagctggactgtatgcac<br>gaaccccccgttcagtccgaccgctgcgccttatccggtaactatcgtcttgagtccaacccggaaagacatgcaaaagcaccactggca<br>gcagccactggtaattgatttagaggagttagtcttgaagtcatgcgccggttaaggctaaactgaaaggacaagttttggtgactgcgctc<br>ctccaagccagttacctcggttcaaagagttggtagctcagagaaccttcgaaaaaccgccctgcaaggcggttttttcgttttcagagcaa<br>gagattacgcgcagaccaaaacgatctcaagaagatcatcttattaagattttggtcatgagattatcaaaaaggatcttcacctagatcctttt<br>ggttcatgtgcagctccatcagcaaaaggggatgataagtttatcaccaccgactatttgcaacagtgccgttgatcgtgctatgatcgactg<br>atgtcatcagcggtggagtgcaatgtcgtgcaatacgaatggcgaaaagccgagctcatcggtcagcttctcaaccttggggttaccccc<br>ggcggtgtgctgctggtccacagctccttccgtagcgtccggcccctcgaagatgggccacttggactgatcgaggccctgcgtgctgc<br>gctgggtccgggagggacgctcgtcatgccctcgtggtcaggtctggacgacgagccgttcgatcctgccacgtcgcccgttacaccgg<br>accttggagttgtctctgacacattctggcgcctgccaaatgtaaagcgcagcgcccatccatttgcctttgcggcagcggggccacaggc<br>agagcagatcatctctgatccattgcccctgccacctcactcgcctgcaagcccggtcgcccgtgtccatgaactcgatgggcaggtactt<br>ctcctcggcgtgggacacgatgccaacacgacgctgcatcttgccgagttgatggcaaaggttccctatggggtgccgagacactgcac<br>cattcttcaggatggcaagttggtacgcgtcgattatctcgagaatgaccactgctgtgagcgctttgccttggcggacaggtggctcaagg<br>agaagagccttcagaaggaaggtccagtcggtcatgcctttgctcggttgatccgctcccgcgacattgtggcgacagccctgggtcaac<br>tgggccgagatccgttgatcttcctgcatccgccagaggcgggatgcgaagaatgcgatgccgctcgccagtcgattggctgagctcatg<br>agcggagaacgagatgacgttggaggggcaaggtcgcgctgattgctggggcaacacgtggagcggatcggggattgtctttcttcag<br>ctcgctgatgatatgctgacgctcaatgccgtttggcctccgactaacgaaaatcccgcatttggacggctgatccgattggcacggcgga<br>cggcgaatggcggagcagacgctcgtccgggggcaatgagatatgaaaaagcctgaactcaccgcgacgtatcgggccctggccagc<br>tagctagagtcgacctgggtccccggggatcggtcttgccttgctcgtcggtgatgtacttcaccagctccgcgaagtcgctcttcttgatg<br>gagcgcatggggacgtgcttggcaatcacgcgcaccccccggccgttttagcggctaaaaaagtcatggctctgccctcgggcggacc<br>acgcccatcatgaccttgccaagctcgtcctgcttctcttcgatcttcgccagcagggcgaggatcgtggcatcaccgaaccgcgccgtg<br>cgcgggtcgtcggtgagccagagtttcagcaggccgcccaggcggcccaggtcgccattgatgcgggccagctcgcggacgtgctca<br>tagtccacgacgcccgtgattttgtagccctggccgacggccagcaggtaggccgacaggctcatgccggccgccgccgccttttcctc<br>aatcgctcttcgttcgtctggaaggcagtacaccttgataggtgggctgcccttcctggttggcttggtttcatcagccatccgcttgccctca<br>tctgttacgccggcggtagccggccagcctcgcagagcaggattcccgttgagcaccgccaggtgcgaataagggacagtgaagaag<br>gaacacccgctcgcgggtgggcctacttcacctatcctgcccggctgacgccgttggatacaccaaggaaagtctacacgaaccctttgg<br>caaaatcctgtatatcgtgcgaaaaaggatggatataccgaaaaaatcgctataatgaccccgaagcagggttatgcagcggaaaagatc<br>cgtcgacctgctacgccgctacgtcttccgtgccgtcctgggcgtcgtcttcgtcgtcgtcggtcggcggcttcgcccacgtgatcgaagc<br>gcgcttctcgatgggcgttccctgccccctgcccgtagtcgacttcgtgacaacgatcttgtctacgaagagcccgacgaacacgcgcttg<br>tcgtctactgacgcgcgcccccaccacgacttagggccggtcgggtcagcgtcggcgtcttcggggaaccattggtcaaggggaagctt<br>cggggcttcggcggcttcaagttcggcaagccgctcttccgccccttgctgccggagcgtcagcgctgcctgttgcttccggaagtgcttc<br>ctgccaacgggtccgtcgtacgcgcctgccgcgcggtcttcgtacagctcttcaagggcgttcagggcgtcggcgcgctccgcaacaag<br>gttcgcccgttcgccgctcttctcaggcgcctcagtgagcttgccgaagcgtcgggcggcttcccacagaagcgccaacgtctcttcgtc<br>gccttcggcgtgcctgatcttgttgaagatgcgttccgcaacgaacttgtcgagtgccgccatgctgacgttgcacgtgccttcgtgctgcc<br>caggtgcggacgggtcgaccaccttccggcgacggcagcggtaagagtccttgatcgattcttccccgcgcttcgaagtcatgacggcg<br>ccacactcgcagtacagcttgtccatggcggacagaatggcttgcccccgggaaagccccttgccgcgccccctgccgtccaaccacg |

| | |
|---|---|
| | cctgaagctcataccactcagcgggctcgatgatcggtccgcaatcaagctcgaccggccggagcgtgatcgggtcgcgctgaatgcg<br>gtaaccctcaatcttcgtggtcggcgtgccgtccggcttcttcttgtagatcacctcagcggcgaagcccgcaatacgcgggtcccgaag<br>gattcgcataacggttgccgggtcccaggcgcttgaagcggtcttcttcccaatcgtctcgccccgggtcggcacggcgtcagcgtccat<br>gcgcttacaaagccccgtgatgctgcccgggtgaatggcggcttgactgcccggcttgaagggaaggtgtttgtgcgtcttgatctcacgc<br>caccaccaccggattacgtcgggctcgaactcgaagggtccggtaaggggagtggtcgagtgcgcaagcttgttgatgacgacattgac<br>cattcggccgttgcgcgtgatctccttcgtctccgaaacaagctcgaagccgtaaggcgccttcccgccgacgtacccgcccaattcgcg<br>ctgaaggttcttcgtgtcgagaatcttcgccgacttcagcgaagattctttgtgcgacgcgtcgagccgcataatcaggtgaatcaggtcca<br>tgacgtttccctgccggaagacgccttcctgagtggaaacaatcgtcacgcccagggcgagcaattccgagacaatcggaatcgcgtcc<br>atgaccttcaggcgcgagaagcgcgacacgtcatagacaatgatcatgttgagccgcccggcgcggcattcgttcaggatgcgttcgaa<br>ctccgggcgctccgccgtcccgaacgccgacgtgcccggcgcttcgctgaaatgcccgacgaacctgaaccggcccccgtcgcgctc<br>gacttcgcgctgaaggtcggccgccttgtcttcgttggcgctacgctgtgtcgctgggcttgctgcgctcgaactctcgcgctcgcgcgac<br>tgacggtcgtaagcacccgcgtacgtgtccaccccggtcacaaccccttgtgtcatgtcggcgaccctacgcccccaactgagagaactc<br>aaaggttaccccagttggggcactactcccgaaaaccgcttctgacctgggaaaacgtgaagccccggggcatccgctgagggttgcc<br>gccggggcttcggtgtgtccgtcagtacgggccatagaggggcgtcgtgtaacccacccagggcgattgctccggcacggggaagcc<br>gcgccacgccttcgggacgtctggaatcgctagagcttgcatgc |
| pPAU | tgccacctgacgtctaagaagatcccgcaaaagcggcctttgactccctgcaagcctcagcgaccgaatatatcggttatgcgtgggcga<br>tggttgttgtcattgtcggcgcaactatcggtatcaagctgtttaagaaattcacctcgaaagcaagctgataaaccgatacaattaaaggct<br>ccttttggagcctttttttttggagattttcaacgtgaaaaaattattattcgcaattcctttagttgttcctttctattctcactccgctgaaactgttg<br>aaagttgtttagcaaaacctcatacagaaaattcatattaccctgttatccctactccgagacagtcagagggtagaattccttctagagcctc<br>agagacccggccgggtctcaggagtactagttcctgcagcactcgttgctttatcggtatattaccctgttatccctaattttgttatcaataa<br>aaaggcccccccgttagggaggccttattgttcgtcgctcactcaaaggcggtaatgacgctcagtggaacgaaaactcacgttaagggta<br>aaactgtattataagtaaatgcatgtatactaaactcacaaattagagcttcaatttaattatatcagttattacccataacttcgtatagcatacat<br>tatacgaagttatcccgggtaccgagctcgattcgtaacttacacgcgcctcgtatcttttaatgatggaataatttgggaatttactctgtgttt<br>atttatttttatgttttgtatttggattttagaaagtaaataaagaaggtagaagagttacggaatgaagaaaaaaaaataaacaaaggtttaaaa<br>aatttcaacaaaaagcgtactttacatatatatttattagacaagaaaagcagattaaatagatatacattcgattaacgataagtaaaatgtaa<br>aatcacaggattttcgtgtgtggtcttctacacagacaagatgaaacaattcggcattaatacctgagagcaggaagagcaagataaaagg<br>tagtatttgttggcgatccccctagagtcttttacatcttcggaaaacaaaaactattttttctttaatttctttttttactttctatttttaatttatatattt<br>atattaaaaaatttaaattataattatttttatagcacgtgagcgctagcggagtgtatactcggttatccacagaatcaggggataacgcagg<br>aaagaacatgtgagcaaaaggccagcaaaaggccaggaaccgtaaaaaggccgcgttgctggcgtttttccacaggctccgcccccct<br>gacgagcatcacaaaaatcgacgctcaagtcagaggtggcgaaacccgacaggactataaagataccaggcgtttccccctggaagct<br>ccctcgtgcgctctcctgttccgaccctgccgcttaccggatacctgtccgcctttctcccttcgggaagcgtggcgctttctcatagctcac<br>gctgtaggtatctcagttcggtgtaggtcgttcgctccaagctgggctgtgtgcacgaaccccccgttcagcccgaccgctgcgccttatcc<br>ggtaactatcgtcttgagtccaacccggtaagacacgacttatcgccactggcagcagccactggtaacaggattagcagagcgaggtat<br>gtaggcggtgctacagagttcttgaagtggtggcctaactacggctacactagaagaacagtatttggtatctgcgctctgctgaagccagt<br>taccttcggaaaaagagttggtagctcttgatccggcaaacaaaccaccgctggtagcggtggtttttttgtttgcaagcagcagattacgc<br>gcagaaaaaaaggatctcaagaagatcctttgatcttttctacgggattttggtcatgagattatcaaaaaggatcttcacctagatccttttgg<br>ttcatgtgcagctccatcagcaaaaggggatgataagtttatcaccaccgactatttgcaacagtgccgttgatcgtgctatgatcgactgat<br>gtcatcagcggtggagtgcaatgtcgtgcaatacgaatggcgaaaagccgagctcatcggtcagcttctcaaccttggggttacccccgg<br>cggtgtgctgctggtccacagctccttccgtagcgtccggcccctcgaagatgggccacttggactgatcgaggccctgcgtgctgcgct<br>gggtccgggagggacgctcgtcatgccctcgtggtcaggtctggacgacgagccgttcgatcctgccacgtcgcccgttacaccggac<br>cttggagttgtctctgacacattctggcgcctgccaaatgtaaagcgcagcgcccatccatttgcctttgcggcagcggggccacaggca<br>gagcagatcatctctgatccattgcccctgccacctcactcgcctgcaagcccggtcgcccgtgtccatgaactcgatgggcaggtacttc<br>tcctcggcgtgggacacgatgccaacacgacgctgcatcttgccgagttgatggcaaaggttccctatggggtgccgagacactgcacc<br>attcttcaggatggcaagttggtacgcgtcgattatctcgagaatgaccactgctgtgagcgctttgccttggcggacaggtggctcaagga<br>gaagagccttcagaaggaaggtccagtcggtcatgcctttgctcggttgatccgctcccgcgacattgtggcgacagccctgggtcaact<br>gggccgagatccgttgatcttcctgcatccgccagaggcgggatgcgaagaatgcgatgccgctcgccagtcgattggctgagctcatg<br>agcggagaacgagatgacgttggaggggcaaggtcgcgctgattgctggggcaacacgtggagcggatcggggattgtctttcttcag<br>ctcgctgatgatatgctgacgctcaatgccgtttggcctccgactaacgaaaatcccgcatttggacggctgatccgattggcacggcgga<br>cggcgaatggcggagcagacgctcgtccgggggcaatgagatatgaaaaagcctgaactcaccgcgacgtatcgggccctggccagc<br>tagctagagtcgacctgggtccccggggatcggtcttgccttgctcgtcggtgatgtacttcaccagctccgcgaagtcgctcttcttgatg<br>gagcgcatggggacgtgcttggcaatcacgcgcacccccggccgttttagcggctaaaaaagtcatggctctgccctcgggcggacc<br>acgcccatcatgaccttgccaagctcgtcctgcttctcttcgatcttcgccagcagggcgaggatcgtggcatcaccgaaccgcgccgtg<br>cgcgggtcgtcggtgagccagagtttcagcaggccgcccaggcggcccaggtcgccattgatgcgggccagctcgcggacgtgctca<br>tagtccacgacgcccgtgattttgtagccctggccgacggccagcaggtaggccgacaggctcatgccggccgccgccgccttttcctc<br>aatcgctcttcgttcgtctggaaggcagtacacctgataggtgggctgcccttcctggttggcttggtttcatcagccatccgcttgccctca<br>tctgttacgccggcggtagccggccagcctcgcagagcaggattcccgttgagcaccgccaggtgcgaataagggacagtgaagaag<br>gaacacccgctcgcgggtgggcctacttcacctatcctgcccggctgacgccgttggatacaccaaggaaagtctacacgaaccctttgg<br>caaaatcctgtatatcgtgcgaaaaaggatggatataccgaaaaaatcgctataatgaccccgaagcagggttatgcagcggaaaagatc<br>cgtcgacctgctacgccgctacgtcttccgtgccgtcctgggcgtcgtcttcgtcgtcggtcggcggcttcgcccacgtgatcgaagc<br>gcgcttctcgatgggcgttccctgcccccctgcccgtagtcgacttcgtgacaacgatcttgtctacgaagagcccgacgaacacgcgcttg<br>tcgtctactgacgcgcgcccccaccacgacttagggccggtcgggtcagcgtcggcgtcttcggggaaccattggtcaaggggaagctt<br>cggggcttcggcggcttcaagttcggcaagccgctcttccgccccttgctgccggagcgtcagcgctgcctgttgcttccggaagtgcttc<br>ctgccaacgggtccgtcgtacgcgcctgccgcgcggtcttcgtacagctcttcaagggcgttcagggcgtcggcgcgctccgcaacaag<br>gttcgcccgttcgccgctcttctcaggcgcctcagtgagcttgccgaagcgtcgggcggcttcccacagaagcgccaacgtctcttcgtc<br>gccttcggcgtgcctgatcttgttgaagatgcgttccgcaacgaacttgtcgagtgccgccatgctgacgttgcacgtgccttcgtgctgcc<br>caggtgcggacgggtcgaccaccttccggcgacggcagcggtaagagtccttgatcgattcttccccgcgcttcgaagtcatgacggcg<br>ccacactcgcagtacagcttgtccatggcggacagaatggcttgcccccgggaaagccccttgccgcgcccccctgccgtccaaccacg |

| | |
|---|---|
| | cctgaagctcataccactcagcgggctcgatgatcggtccgcaatcaagctcgaccggccggagcgtgatcgggtcgcgctgaatgcg<br>gtaaccctcaatcttcgtggtcggcgtgccgtccggcttcttcttgtagatcacctcagcggcgaagcccgcaatacgcgggtcccgaag<br>gattcgcataacggttgccgggtcccaggcgcttgaagcggtcttcttcccaatcgtctcgccccgggtcggcacggcgtcagcgtccat<br>gcgcttacaaagccccgtgatgctgcccgggtgaatggcggcttgactgcccggcttgaagggaaggtgtttgtgcgtcttgatctcacgc<br>caccaccaccggattacgtcgggctcgaactcgaagggtccggtaaggggagtggtcgagtgcgcaagcttgttgatgacgacattgac<br>cattcggccgttgcgcgtgatctccttcgtctccgaaacaagctcgaagccgtaaggcgccttcccgccgacgtacccgcccaattcgcg<br>ctgaaggttcttcgtgtcgagaatcttcgccgacttcagcgaagattctttgtgcgacgcgtcgagccgcataatcaggtgaatcaggtcca<br>tgacgtttccctgccggaagacgccttcctgagtggaaacaatcgtcacgcccagggcgagcaattccgagacaatcggaatcgcgtcc<br>atgaccttcaggcgcgagaagcgcgacacgtcatagacaatgatcatgttgagccgcccggcgcggcattcgttcaggatgcgttcgaa<br>ctccgggcgctccgccgtcccgaacgccgacgtgcccggcgcttcgctgaaatgcccgacgaacctgaaccggcccccgtcgcgctc<br>gacttcgcgctgaaggtcggccgccttgtcttcgttggcgctacgctgtgtcgctgggcttgctgcgctcgaactctcgcgctcgcgcgac<br>tgacggtcgtaagcacccgcgtacgtgtccaccccggtcacaaccccttgtgtcatgtcggcgaccctacgcccccaactgagagaactc<br>aaaggttaccccagttggggcactactcccgaaaaccgcttctgacctgggaaaacgtgaagccccggggcatccgctgagggttgcc<br>gccggggcttcggtgtgtccgtcagtacgggccatagaggggcgtcgtgtaacccacccagggcgattgctccggcacggggaagcc<br>gcgccacgccttcgggacgtctggaatcgctagagcttgcatgc |
| pVHS | tgactgctctgagaaagttgagatcccgcaaaagcggcctttgactccctgcaagcctcagcgaccgaatatatcggttatgcgtgggcga<br>tggttgttgtcattgtcggcgcaactatcggtatcaagctgtttaagaaattcacctcgaaagcaagctgataaaccgatacaattaaaggct<br>ccttttggagcctttttttttggagattttcaacgtgaaaaaattattattcgcaattcctttagttgttcctttctattctcactccgctgaaactgttg<br>aaagttgtttagcaaaacctcatacagaaaattcatattaccctgttatccctactccgagacagtcagagggtagaattccttctagagcctc<br>agagacccggccgggtctcaggagtactagttcctgcagcactcgttgctttatcggtatattaccctgttatccctatggcagaccagaaa<br>caaaaaaaggcccccccgttagggaggccttcaataattggacagacagtattgcttacgaggacgctcagtggaacgaaaactcacgtta<br>agggtaaaactgtattataagtaaatgcatgtatactaaactcacaaattagagcttcaatttaattatatcagttattacccataacttcgtatag<br>catacattatacgaagttatcccgggtaccgagctcgattcgtaacttacacgcgcctcgtatcttttaatgatggaataatttgggaatttactc<br>tgtgtttatttatttttatgttttgtatttggattttagaaagtaaataaagaaggtagaagagttacggaatgaagaaaaaaaaataaacaaagg<br>tttaaaaaatttcaacaaaaagcgtactttacatatatatttattagacaagaaaagcagattaaatagatatacattcgattaacgataagtaaa<br>atgtaaaatcacaggattttcgtgtgtggtcttctacacagacaagatgaaacaattcggcattaatacctgagagcaggaagagcaagata<br>aaaggtagtatttgttggcgatccccctagagtcttttacatcttcggaaaacaaaaactatttttttctttaatttcttttttttactttctatttttaatttat<br>atatttatattaaaaaatttaaattataattatttttatagcacgtgaggctagagctgtcagaccaagtttacgagctcgcttggactcctgttgat<br>agatccagtaatgacctcagaactccatctggatttgttcagaacgctcggttgccgccgggcgttttttattggtgagaatccaagcactag<br>ggacagtaagacgggtaagcctgttgatgataccgctgccttactgggtgcattagccagtctgaatgacctgtcacgggataatccgaag<br>tggtcagactggaaaatcagagggcaggaactgctgaacagcaaaaagtcagatagcaccacatagcagacccgccataaaacgccct<br>gagaagcccgtgacgggcttttcttgtattatgggtagtttccttgcatgaatccataaaaggcgcctgtagtgccatttacccccattcactg<br>ccagagccgtgagcgcagcgaactgaatgtcacgaaaaagacagcgactcaggtgcctgatggtcggagacaaaaggaatattcagc<br>gatttgcccgagcttgcgagggtgctacttaagcctttagggttttaaggtctgttttgtagaggagcaaacagcgtttgcgacatccttttgta<br>atactgcggaactgactaaagtagtgagttatacacagggctgggatctattctttttatctttttttattctttctttattctataaattataaccactt<br>gaatataaacaaaaaaaacacacaaaggtctagcggaatttacagagggtctagcagaatttacaagttttccagcaaaggtctagcagaa<br>tttacagatacccacaactcaaaggaaaaggacatgtaattatcattgactagcccatctcaattggtatagtgattaaaatcacctagaccaa<br>ttgagatgtatgtctgaattagttgttttcaaagcaaatgaactagcgattagtcgctatgacttaacggagcatgaaaccaagctaattttatg<br>ctgtgtggcactactcaaccccacgattgaaaaccctacaaggaaagaacggacggtatcgttcacttataaccaatacgctcagatgatg<br>aacatcagtagggaaaatgcttatggtgtattagctaaagcaaccagagagctgatgacgagaactgtggaaatcaggaatcctttggtta<br>aaggctttgagattttccagtggacaaactatgccaagttctcaagcgaaaaattagaattagtttttagtgaagagatattgccttatcttttcc<br>agttaaaaaaattcataaaatataatctggaacatgttaagtcttttgaaaacaaatactctatgaggatttatgagtggttattaaaagaactaa<br>cacaaaagaaaactcacaaggcaaatatagagattagccttgatgaatttaagttcatgttaatgcttgaaaataactaccatgagtttaaaag<br>gcttaaccaatgggttttgaaaccaataagtaaagatttaaacacttacagcaatatgaaattggtggttgataagcgaggccgcccgactg<br>atacgttgattttccaagttgaactagatagacaaatggatctcgtaaccgaacttgagaacaaccagataaaaatgaatggtgacaaaata<br>ccaacaaccattacatcagattcctacctacgtaacggactaagaaaaacactacacgatgctttaactgcaaaaattcagctcaccagtttt<br>gaggcaaaatttttgagtgacatgcaaagtaagcatgatctcaatggttcgttctcatggctcacgcaaaaacaacgaaccacactagaga<br>acatactggctaaatacggaaggatctgaggttcttatggctcttgtatctatcagtgaagcatcaagactaacaaacaaaagtagaacaact<br>gttcaccgttagatatcaaagggaaaactgtccatatgcacagatgaaaacggtgtaaaaaagatagatacatcagagcttttacgagttttt<br>ggtgcatttaaagctgttcaccatgaacagatcgacaatgtaactactagagggttcatgtgcagctccatcagcaaaaggggatgataagt<br>ttatcaccaccgactatttgcaacagtgccgttgatcgtgctatgatcgactggatcggcggggcctgggacggcgcctcgccgtcgcag<br>aaccaggcggtggcgtacaccgtcgcctcggtcggcccgtagagattggcgatcccgaccgcagcaccaccgagaacgtccccgacg<br>tggccgaccagcccgtcatcgtcaacgcctgatccgcggtgcggacaggccgtgtcgcgaccggccgtgcggaattaagccggcccg<br>taccctgtgaatagaggtccgctgtgacacaagaatccctgttacttctcgaccgtattgattcggatgattcctacgcgagcctgcggaac<br>gaccaggagttctgggagccgctggcccgccgagccctggaggagctcgggctgccggtgccgccggtgctgcgggtgcccggcga<br>gagcaccaaccccgtactggtcggcgagcccggcccggtgatcaagctgttcggcgagcactggtgcggtccggagagcctcgcgtc<br>ggagtcggaggcgtacgcggtcctggcggacgccccggtgccggtgccccgcctcctcggccgcggcgagctgcggcccggcacc<br>ggagcctggccgtggccctacctggtgatgagccggatgaccggcaccacctggcggtccgcgatggacggcacgaccgaccggaa<br>cgcgctgctcgccctggcccgcgaactcggccgggtgctcggccggctgcacagggtgccgctgaccgggaacaccgtgctcacccc<br>ccattccgaggtcttcccggaactgctgcgggaacgccgcgcggcgaccgtcgaggaccaccgcgggtggggctacctctcgccccg<br>gctgctggaccgcctggaggactggctgccggacgtggacacgctgctggccggccgcgaaccccggttcgtccacggcgacctgca<br>cgggaccaacatcttcgtggacctggccgcgaccgaggtcaccgggatcgtcgacttcaccgacgtctatgcgggagactcccgctaca<br>gcctggtgcaactgcatctcaacgccttccggggcgaccgcgagatcctggccgcgctgctcgacggggcgcagtggaagcggaccg<br>aggacttcgcccgcgaactgctcgccttcaccttcctgcacgacttcgaggtgttcgaggagacgccgctggatctctccggcttcaccga<br>tccggaggaactggcgcagttcctctgggggccgccggacaccgcccccggcgcctgacgccccgggccgcccggcgccgccccc<br>ggcccccggcggccgccgggcggagccccgcccggcggcccgggagccccgggcccgcgccgaagcccgctgctgcgagctag<br>ctagagtcgacctgggtccccggggatcggtcttgccttgctcgtcggtgatgtacttcaccagctccgcgaagtcgctcttcttgatggag |

| | |
|---|---|
| | cgcatggggacgtgcttggcaatcacgcgcaccccccggccgttttagcggctaaaaaagtcatggctctgccctcgggcggaccacg<br>cccatcatgaccttgccaagctcgtcctgcttctcttcgatcttcgccagcagggcgaggatcgtggcatcaccgaaccgcgccgtgcgc<br>gggtcgtcggtgagccagagtttcagcaggccgcccaggcggcccaggtcgccattgatgcgggccagctcgcggacgtgctcatagt<br>ccacgacgcccgtgattttgtagccctggccgacggccagcaggtaggccgacaggctcatgccggccgccgccgccttttcctcaatc<br>gctcttcgttcgtctggaaggcagtacaccttgataggtgggctgcccttcctggttggcttggtttcatcagccatccgcttgccctcatctgt<br>tacgccggcggtagccggccagcctcgcagagcaggattcccgttgagcaccgccaggtgcgaataagggacagtgaagaaggaac<br>acccgctcgcgggtgggcctacttcacctatcctgcccggctgacgccgttggatacaccaaggaaagtctacacgaaccctttggcaaa<br>atcctgtatatcgtgcgaaaaaggatggatataccgaaaaaatcgctataatgaccccgaagcagggttatgcagcggaaaagatccgtc<br>gacctgagcagcagcgttccaagctgatcgagatgagcgcgctgaacgagtccaacaagcagtagccgacgcccgtacaaccgtcgg<br>cctgcctcgcgggtctaacgaaaatggactccaccactccccgtgattcctcgggcacttggtgcacgcagagtggtcgcatattcaccta<br>caagggggtacggtcggtcgagcggcctgccctcccccatggccgacggtcaaacgtcgcctgcctgcccgggggatacccatgtgc<br>attcgtgtccgcttcgcgcctctcgacccgctcaacttccggccgtacgacgccgctggaaacacggtcaccctgcctgccaccctcccc<br>cgggatgcttccctcgtagcccttcgagccgtccttgaagaactggctgtagagcagcccccggacggtgcagtctgctggtgtggggc<br>agccgtacacatcctgccccgcgttcccgaacagcggaggagcggacaggtgacccatggcgcctagagcgacgaacaacccacgg<br>cagttgagggcgaagagctgcggctgccagctgtgcatggagaagtacccgcccgagaagtacggagagcggaaccgccgacgcg<br>actgcaccggctcgtggcaggcgcgttaccgcgacccggccggcaaccagaagcagaaatgctttgcgatcaaggacggcggtaaga<br>aggcagccgaggcgcacctcgacaagatccgcacgcaggtccgcgaacggacgtacgccgacccgaagcgtggcgagatcaccct<br>gtcccagtggtggaaactgtggtgggaggcgcagccggaccgagcagtcacgaccgccaaccggaagcggtcgaactgggccgcg<br>cacatcgagccgaagtgggggcagtggcgtctctgcgacttggagtacatcgagctccaggcgtggatcacgaaggaggtgaagggct<br>accacacccggaagaaggttcatgaggtgctgaactcgatgctccgggccgccgtcaaggacggccggcgtatcccgttcaacccggc<br>ggccgacctggacattggcgaggcgccggcgaagcatccggacgaactgatgccgcccgaccgcgcgcagtgcgcgctgatcgtca<br>gtcacctcccgatgtactaccggccgctcgtcgtcttccttgaacacaccggcctccggtggggcgaggcgacggcgctgcgctggga<br>gaacgtcgacctggacgcccactacctcaaggtgaaggaagtgctcagtgacgacgaaggcaagctgttccggaagcctgcgccgaa<br>gagcaacgccgggttccgcacggtcccgctcacgccgcaggccgaggacgcgatccgcaccatggtcacccggtggcggccgactc<br>ccacgatcaccccgattggcgaggacccgtacgacctcgcgccggatgagctcgtgttccgcggcccacagggcggcgtcctgaccc<br>ggcacaacttccggcgcacatggatccctgcaatcaaggcggcaggcctcgcccgcgaggtgaagaaccgggacaccggccgcatg<br>gagtggtggccgcgggtgcacgaccttcgccacgtgttcgccacgtggctcaaggatgtgggcattgacgagaaggacacgcagacc<br>gtgatgggtcacgagcgagggtcgaaggtgacgtggttgtaccagcattcgccggccgacgtggcggcgaaggtgcgggcggcgat<br>ggctcccgagacggagggtgttcgaacgctgcgggcggtgtgacgccggatgccacgcagatgccacagggatgccacaacacccc<br>ctcactgagactcaccgagactcactgaaactcatttatggaggtgaagcccctatggcgacaggctcactgagactcactcagactcact<br>gaggctcatgatcgctttacgttctctcctaaagcgggtgtcgcaggttcgaatcctgccgggggcacaacctgcatcgcaggtcaggga<br>gtcaacggcccccccgttccattcgaacgggggccgttgtcgtacccggatgccacatagatgccacatccccacggaatcctgcggat<br>cacgtcgctcgaaagagt |
| pVSP | tgactgctctgagaaagttgagatcccgcaaaagcggcctttgactccctgcaagcctcagcgaccgaatatatcggttatgcgtgggcga<br>tggttgttgtcattgtcggcgcaactatcggtatcaagctgtttaagaaattcacctcgaaagcaagctgataaaccgatacaattaaaggct<br>ccttttggagcctttttttttggagattttcaacgtgaaaaaattattattcgcaattcctttagttgttcctttctattctcactccgctgaaactgttg<br>aaagttgtttagcaaaacctcatacagaaaattcatattaccctgttatccctactccgagacagtcagagggtagaattccttctagagcctc<br>agagacccggccgggtctcaggagtactagttcctgcagcactcgttgctttatcggtatattaccctgttatccctatggcagaccagaaa<br>caaaaaaaggcccccccgttagggaggccttcaataattggacagacagtattgcttacgaggacgctcagtggaacgaaaactcacgtta<br>agggtaaaactgtattataagtaaatgcatgtatactaaactcacaaattagagcttcaatttaattatatcagttattacccataacttcgtatag<br>catacattatacgaagttatcccgggtaccgagctcgattcgtaacttacacgcgcctcgtatcttttaatgatggaataatttgggaatttactc<br>tgtgtttatttatttttatgttttgtatttggattttagaaagtaaataaagaaggtagaagagttacggaatgaagaaaaaaaaaataaacaaagg<br>tttaaaaaatttcaacaaaaagcgtactttacatatatatttattagacaagaaaagcagattaaatagatatacattcgattaacgataagtaaa<br>atgtaaaatcacaggattttcgtgtgtggtcttctacacagacaagatgaaacaattcggcattaatacctgagagcaggaagagcaagata<br>aaaggtagtatttgttggcgatccccctagagtcttttacatcttcggaaaacaaaaactatttttttctttaatttctttttttactttctatttttaatttat<br>atatttatattaaaaaatttaaattataattatttttatagcacgtgagcgctagcggagtgtatactggcttactatgttggcactgatgagggtg<br>tcagtgaagtgcttcatgtggcaggagaaaaaaggctgcaccggtgcgtcagcagaatatgtgatacaggatatattccgcttcctcgctc<br>actgactcgctacgctcggtcgttcgactgcggcgagcggaaatggcttacgaacggggcggagatttcctggaagatgccaggaagat<br>acttaacagggaagtgagagggccgcggcaaagccgtttttccataggctccgcccccctgacaagcatcacgaaatctgacgctcaaa<br>tcagtggtggcgaaacccgacaggactataaagataccaggcgtttccccctggcggctccctcgtgcgctctcctgttcctgcctttcggttt<br>accggtgtcattccgctgttatggccgcgtttgtctcattccacgcctgacactcagttccgggtaggcagttcgctccaagctggactgtat<br>gcacgaaccccccgttcagtccgaccgctgcgccttatccggtaactatcgtcttgagtccaacccggaaagacatgcaaaagcaccact<br>ggcagcagccactggtaattgatttagaggagttagtcttgaagtcatgcgccggttaaggctaaactgaaaggacaagttttggtgactgc<br>gctcctccaagccagttacctcggttcaaagagttggtagctcagagaaccttcgaaaaaccgccctgcaaggcggttttttcgttttcaga<br>gcaagagattacgcgcagaccaaaacgatctcaagaagatcatcttattaagccagccaggacagaaatgcctcgacttcgctgctaccc<br>aaggttgccgggtgacgcacaccgtggaaacggatgaaggcacgaacccagtggacataagcctgttcggttcgtaagctgtaatgcaa<br>gtagcgtatgcgctcacgcaactggtccagaaccttgaccgaacgcagcggtggtaacggcgcagtggcggttttcatggcttgttatga<br>ctgtttttttggggtacagtctatgcctcgggcatccaagcagcaagcgcgttacgccgtgggtcgatgtttgatgttatggagcagcaacg<br>atgttacgcagcagggcagtcgccctaaaacaaagttaaacattatgagggaagcggtgatcgccgaagtatcgactcaactatcagagg<br>tagttggcgtcatcgagcgccatctcgaaccgacgttgctggccgtacatttgtacggctccgcagtggatggcggcctgaagccacaca<br>gtgatattgatttgctggttacggtgaccgtaaggcttgatgaaacaacgcggcgagctttgatcaacgaccttttggaaacttcggcttccc<br>ctggagagagcgagattctccgcgctgtagaagtcaccattgttgtgcacgacgacatcattccgtggcgttatccagctaagcgcgaact<br>gcaatttggagaatggcagcgcaatgacattcttgcaggtatcttcgagccagccacgatcgacattgatctggctatcttgctgacaaaag<br>caagagaacatagcgttgccttggtaggtccagcggcggaggaactctttgatccggttcctgaacaggatctatttgaggcgctaaatga<br>aaccttaacgctatggaactcgccgcccgactgggctggcgatgagcgaaatgtagtgcttacgttgtcccgcatttggtacagcgcagta<br>accggcaaaatcgcgccgaaggatgtcgctgccgactgggcaatggagcgcctgccggcccagtatcagcccgtcatacttgaagcta<br>gacaggcttatcttggacaagaagaagatcgcttggcctcgcgcgcagatcagttggaagaatttgtccactacgtgaaaggcgagatca |

| | |
|---|---|
| | ccaaggtagtcggcaaataaccctcgagccacccatgaccaaaatcccttaacgtgagttacgcgtcgttccactgagcgtcagacggtc<br>cccggggatcggtcttgccttgctcgtcggtgatgtacttcaccagctccgcgaagtcgctcttcttgatggagcgcatggggacgtgcttg<br>gcaatcacgcgcaccccccggccgttttagcggctaaaaaagtcatggctctgccctcgggcggaccacgcccatcatgaccttgccaa<br>gctcgtcctgcttctcttcgatcttcgccagcagggcgaggatcgtggcatcaccgaaccgcgccgtgcgcgggtcgtcggtgagccag<br>agtttcagcaggccgcccaggcggcccaggtcgccattgatgcgggccagctcgcggacgtgctcatagtccacgacgcccgtgatttt<br>gtagccctggccgacggccagcaggtaggccgacaggctcatgccggccgccgccgccttttcctcaatcgctcttcgttcgtctggaag<br>gcagtacaccttgataggtgggctgcccttcctggttggcttggtttcatcagccatccgcttgccctcatctgttacgccggcggtagccgg<br>ccagcctcgcagagcaggattcccgttgagcaccgccaggtgcgaataagggacagtgaagaaggaacacccgctcgcgggtgggc<br>ctacttcacctatcctgcccggctgacgccgttggatacaccaaggaaagtatacacgaaccctttggcaaaatcctgtatatcgtgcgaaa<br>aaggatggatataccgaaaaaatcgctataatgaccccgaagcagggttatgcagcggaaaagatccgtcgacctgagcagcagcgttc<br>caagctgatcgagatgagcgcgctgaacgagtccaacaagcagtagccgacgcccgtacaaccgtcggcctgcctcgcgggtctaac<br>gaaaatggactccaccactccccgtgattcctcgggcacttggtgcacgcagagtggtcgcatattcacctacaagggggtacggtcggt<br>cgagcggcctgccctcccccatggccgacggtcaaacgtcgcctgcctgcccgggggatacccatgtgcattcgtgtccgcttcgcgcc<br>tctcgacccgctcaacttccggccgtacgacgccgctggaaacacggtcaccctgcctgccaccctcccccgggatgcttccctcgtagc<br>ccttcgagccgtccttgaagaactggctgtagagcagcccccggacggtgcagtctgctggtgtggggcagccgtacacatcctgcccc<br>gcgttcccgaacagcggaggagcggacaggtgacccatggcgcctagagcgacgaacaacccacggcagttgagggcgaagagct<br>gcggctgccagctgtgcatggagaagtacccgcccgagaagtacggagagcggaaccgccgacgcgactgcaccggctcgtggcag<br>gcgcgttaccgcgacccggccggcaaccagaagcagaaatgctttgcgatcaaggacggcggtaagaaggcagccgaggcgcacct<br>cgacaagatccgcacgcaggtccgcgaacggacgtacgccgacccgaagcgtggcgagatcaccctgtcccagtggtggaaactgtg<br>gtgggaggcgcagccggaccgagcagtcacgaccgccaaccggaagcggtcgaactgggccgcgcacatcgagccgaagtgggg<br>gcagtggcgtctctgcgacttggagtacatcgagctccaggcgtggatcacgaaggaggtgaagggctaccacacccggaagaaggtt<br>catgaggtgctgaactcgatgctccgggccgccgtcaaggacggccggcgtatcccgttcaacccggcggccgacctggacattggcg<br>aggcgccggcgaagcatccggacgaactgatgccgcccgaccgcgcgcagtgcgcgctgatcgtcagtcacctcccgatgtactacc<br>ggccgctcgtcgtcttccttgaacacaccggcctccggtggggcgaggcgacggcgctgcgctgggagaacgtcgacctggacgccc<br>actacctcaaggtgaaggaagtgctcagtgacgacgaaggcaagctgttccggaagcctgcgccgaagagcaacgccgggttccgca<br>cggtcccgctcacgccgcaggccgaggacgcgatccgcaccatggtcacccggtggcggccgactcccacgatcaccccgattggcg<br>aggacccgtacgacctcgcgccggatgagctcgtgttccgcggcccacagggcggcgtcctgacccggcacaacttccggcgcacat<br>ggatccctgcaatcaaggcggcaggcctcgcccgcgaggtgaagaaccgggacaccggccgcatggagtggtggccgcgggtgca<br>cgaccttcgccacgtgttcgccacgtggctcaaggatgtgggcattgacgagaaggacacgcagaccgtgatgggtcacgagcgagg<br>gtcgaaggtgacgtggttgtaccagcattcgccggccgacgtggcggcgaaggtgcgggcggcgatggctcccgagacggagggtgt<br>tcgaacgctgcgggcggtgtgacgccggatgccacgcagatgccacagggatgccacaacaccccctcactgagactcaccgagact<br>cactgaaactcatttatggaggtgaagcccctatggcgacaggctcactgagactcactcagactcactgaggctcatgatcgctttacgtt<br>ctctcctaaagcgggtgtcgcaggttcgaatcctgccgggggcacaacctgcatcgcaggtcagggagtcaacggcccccgttccatt<br>cgaacgggggccgttgtcgtacccggatgccacatagatgccacatccccacggaatcctgcggatcacgtcgctcgaaagagt |
| pTHS | agtcagtatccagtcgtgtaggatcccgcaaaagcggcctttgactccctgcaagcctcagcgaccgaatatatcggttatgcgtgggcga<br>tggttgttgtcattgtcggcgcaactatcggtatcaagctgtttaagaaattcacctcgaaagcaagctgataaaccgatacaattaaaggct<br>ccttttggagccttttttttggagattttcaacgtgaaaaaattattattcgcaattcctttagttgttcctttctattctcactccgctgaaactgttg<br>aaagttgtttagcaaaacctcatacagaaaattcatattaccctgttatccctactccgagacagtcagagggtagaattccttctagagcctc<br>agagacccggccgggtctcaggagtactagttcctgcagcactcgttgctttatcggtatattaccctgttatccctaaggaccaaaacgaa<br>aaaaggccccccctttcgggaggcctcttttctggaatttggtaccgagccataaactgccaggcatcagacgctcagtggaacgaaaactc<br>acgttaagggtaaaactgtattataagtaaatgcatgtatactaaactcacaaattagagcttcaatttaattatatcagttattacccataacttc<br>gtatagcatacattatacgaagttatcccgggtaccgagctcgattcgtaacttacacgcgcctcgtatcttttaatgatggaataatttgggaa<br>tttactctgtgtttatttatttttatgttttgtatttggattttagaaagtaaataaagaaggtagaagagttacggaatgaagaaaaaaaaaataaac<br>aaaggtttaaaaaatttcaacaaaaagcgtactttacatatatatttattagacaagaaaagcagattaaatagatatacattcgattaacgata<br>agtaaaatgtaaaatcacaggattttcgtgtgtggtcttctacacagacaagatgaaacaattcggcattaatacctgagagcaggaagagc<br>aagataaaaggtagtatttgttggcgatccccctagagtcttttacatcttcggaaaacaaaaactatttttttctttaatttctttttttactttctatttt<br>taatttatatatttatattaaaaaatttaaattataattatttttatagcacgtgagctagagctgtcagaccaagtttacgagctcgcttggactcct<br>gttgatagatccagtaatgacctcagaactccatctggatttgttcagaacgctcggttgccgccgggcgttttttattggtgagaatccaagc<br>actagggacagtaagacgggtaagcctgttgatgataccgctgccttactgggtgcattagccagtctgaatgacctgtcacgggataatc<br>cgaagtggtcagactggaaaatcagagggcaggaactgctgaacagcaaaaagtcagatagcaccacatagcagacccgccataaaa<br>cgccctgagaagcccgtgacgggcttttcttgtattatgggtagtttccttgcatgaatccataaaaggcgcctgtagtgccatttacccccat<br>tcactgccagagccgtgagcgcagcgaactgaatgtcacgaaaaagacagcgactcaggtgcctgatggtcggagacaaaaggaatat<br>tcagcgatttgcccgagcttgcgagggtgctacttaagcctttagggttttaaggtctgttttgtagaggagcaaacagcgtttgcgacatcct<br>tttgtaatactgcggaactgactaaagtagtgagttatacacagggctgggatctattctttttatctttttttattctttctttattctataaattataac<br>cacttgaatataaacaaaaaaaacacacaaaggtctagcggaatttacagagggtctagcagaatttacaagttttccagcaaaggtctagc<br>agaatttacagatacccacaactcaaaggaaaaggacatgtaattatcattgactagcccatctcaattggtatagtgattaaaatcacctag<br>accaattgagatgtatgtctgaattagttgttttcaaagcaaatgaactagcgattagtcgctatgacttaacggagcatgaaaccaagctaat<br>tttatgctgtgtggcactactcaaccccacgattgaaaaccctacaaggaaagaacggacggtatcgttcacttataaccaatacgctcaga<br>tgatgaacatcagtagggaaaatgcttatggtgtattagctaaagcaaccagagagctgatgacgagaactgtggaaatcaggaatccttt<br>ggttaaaggctttgagattttccagtggacaaactatgccaagttctcaagcgaaaaattagaattagtttttagtgaagagatattgccttatct<br>tttccagttaaaaaaattcataaaatataatctggaacatgttaagtcttttgaaaacaaatactctatgaggatttatgagtggttattaaaagaa<br>ctaacacaaaagaaaactcacaaggcaaatatagagattagccttgatgaatttaagttcatgttaatgcttgaaaataactaccatgagttta<br>aaaggcttaaccaatgggttttgaaaccaataagtaaagatttaaacacttacagcaatatgaaattggtggttgataagcgaggccgcccg<br>actgatacgttgattttccaagttgaactagatagacaaatggatctcgtaaccgaacttgagaacaaccagataaaaatgaatggtgacaa<br>aataccaacaaccattacatcagattcctacctacgtaacggactaagaaaaacactacacgatgctttaactgcaaaaattcagctcacca<br>gttttgaggcaaaatttttgagtgacatgcaaagtaagcatgatctcaatggttcgttctcatggctcacgcaaaaacaacgaaccacactag<br>agaacatactggctaaatacggaaggatctgaggttcttatggctcttgtatctatcagtgaagcatcaagactaacaaacaaaagtagaac |

| | |
|---|---|
| | aactgttcaccgttagatatcaaagggaaaactgtccatatgcacagatgaaaacggtgtaaaaaagatagatacatcagagcttttacgag<br>tttttggtgcatttaaagctgttcaccatgaacagatcgacaatgtaactactagagggttcatgtgcagctccatcagcaaaaggggatgat<br>aagtttatcaccaccgactatttgcaacagtgccgttgatcgtgctatgatcgactggatcggcggggcctgggacggcgcctcgccgtcg<br>cagaaccaggcggtggcgtacaccgtcgcctcggtcggcccgtagagattggcgatcccgaccgcagcaccaccgagaacgtccccg<br>acgtggccgaccagcccgtcatcgtcaacgcctgatccgcggtgcggacaggccgtgtcgcgaccggccgtgcggaattaagccggc<br>ccgtaccctgtgaatagaggtccgctgtgacacaagaatccctgttacttctcgaccgtattgattcggatgattcctacgcgagcctgcgg<br>aacgaccaggagttctgggagccgctggcccgccgagccctggaggagctcgggctgccggtgccgccggtgctgcgggtgcccgg<br>cgagagcaccaaccccgtactggtcggcgagcccggcccggtgatcaagctgttcggcgagcactggtgcggtccggagagcctcgc<br>gtcggagtcggaggcgtacgcggtcctggcggacgccccggtgccggtgccccgcctcctcggccgcggcgagctgcggcccggc<br>accggagcctggccgtggccctacctggtgatgagccggatgaccggcaccacctggcggtccgcgatggacggcacgaccgaccg<br>gaacgcgctgctcgccctggcccgcgaactcggccgggtgctcggccggctgcacagggtgccgctgaccgggaacaccgtgctca<br>ccccccattccgaggtcttcccggaactgctgcgggaacgccgcgcggcgaccgtcgaggaccaccgcgggtggggctacctctcgc<br>cccggctgctggaccgcctggaggactggctgccggacgtggacacgctgctggccggccgcgaaccccggttcgtccacggcgac<br>ctgcacgggaccaacatcttcgtggacctggccgcgaccgaggtcaccgggatcgtcgacttcaccgacgtctatgcgggagactccc<br>gctacagcctggtgcaactgcatctcaacgccttccggggcgaccgcgagatcctggccgcgctgctcgacggggcgcagtggaagc<br>ggaccgaggacttcgcccgcgaactgctcgccttcaccttcctgcacgacttcgaggtgttcgaggagacgccgctggatctctccggct<br>tcaccgatccggaggaactggcgcagttcctctgggggccgccggacaccgcccccggcgcctgacgccccgggccgcccggcgc<br>cgcccccggcccccggcggccgccgggcggagccccgcccggcggcccgggagccccgggcccgcgccgaagcccgctgctgc<br>gagctagctagagtcgacctgggtccccggggatcggtcttgccttgctcgtcggtgatgtacttcaccagctccgcgaagtcgctcttctt<br>gatggagcgcatggggacgtgcttggcaatcacgcgcacccccggccgttttagcggctaaaaaagtcatggctctgccctcgggcg<br>gaccacgcccatcatgaccttgccaagctcgtcctgcttctcttcgatcttcgccagcagggcgaggatcgtggcatcaccgaaccgcgc<br>cgtgcgcgggtcgtcggtgagccagagtttcagcaggccgcccaggcggcccaggtcgccattgatgcgggccagctcgcggacgtg<br>ctcatagtccacgacgcccgtgattttgtagccctggccgacggccagcaggtaggccgacaggctcatgccggccgccgccgcctttt<br>cctcaatcgctcttcgttcgtctggaaggcagtacaccttgataggtgggctgcccttcctggttggcttggtttcatcagccatccgcttgcc<br>ctcatctgttacgccggcggtagccggccagcctcgcagagcaggattcccgttgagcaccgccaggtgcgaataagggacagtgaag<br>aaggaacacccgctcgcgggtgggcctacttcacctatcctgcccggctgacgccgttggatacaccaaggaaagtctacacgaaccct<br>ttggcaaaatcctgtatatcgtgcgaaaaaggatggatataccgaaaaaatcgctataatgaccccgaagcagggttatgcagcggaaaa<br>gatccgtcgacctgtcacgccgccgctgtgaacccgttcagcgtctccggctccgtctcttcctcgaccttcggcttcgcccagtggatacg<br>cacacggtcggcgatcgggggaacccgtccgggccgctgaacgcccttcggaagcttgatcacctcgatccggtccacgaagagccg<br>gacgaacgcccgacggtcttcaaggggcgctagcgcccaccacgaacccggtcccgtcgggtcgctgccccggtcgcccagccattc<br>gccgatcggcagcacgggggagtcggcggcgtcgagctgacgaagccgttcttcggccccttccatgcggagcgtcagcgcggcttc<br>ctcttcgagaaacgcgcgccgtcccatggcggaccggtagccgccgttccgcttgtcttcgtaaagctcttcgagcgccttcacggcgtcc<br>gcgcgctgtgccatcagttcggagcgctgacccttcaactccgggcgttcgtgcgtcgctccccagcgccgtgcggcttcgtacatgagc<br>gctgccgtgtcgccttcgaggtcgtcagggtcggcagggtcgaaggccgtgatgcgcgcgaagatcgcgccgacgatgtacgggtca<br>aggttgtgcatcgtgatcgagcatgacgacccgtcatgaacccggcgggggcacttgcacgcgtacgacgacttgtgagcttgatcacct<br>tcgcggacgttgcccgccatggtcgacccgttgctgtaacccgtctccgggtcgagctggccggagccgtagcagtaaaggacgtccat<br>tgccgacaggagcgattgcccccggtactgacccttcccgcgtcctcgaccctgaagccactcctgaagttcccaccattccgccgggg<br>gaatgtacggctcgaagccgggcagggtcagcggctccatggtgaccgggtcgcgcctgatcttgtaatggctgaagccgccccgcga<br>accgtcggcgcgcaccttgtatgcgatgtcagcttgatacccggcaatgcgcgggtcgctgagtacgcgcttcaaaacgccggggtccc<br>aatcggaaccggcgcgcttcttaccgaccagcgtgccgcgcgtaggcaccttgtcgcggtacagccgctcacaaagcccgttcagcga<br>cccagggtgaaacgacccggctccgccacccttgaagggcgtatcgcggtgcgtcttgatctcctgccacgcccagcggattaccgccc<br>cttcgctgccgtgtgcgccttcccaggtgtgcgcgctgggcaccagtcggcgaatggcaaccagctttccgccgtcttccgggttcggaa<br>ccatttcctcgaccgtgtcgaatccgtacggcgtcgaccccgtgtgtccgcccagccgcttcgccaattccttagcgttcgacacggcgac<br>gctcttgttcttcgactcatcatgcgaagcctgaaggcgcatgatcaggtgaataaggtccatcatttcgccggggcggaatgtgccttcgtt<br>cacgctgacaatggtcacgcccagccggagcaattccgtgacgaccggaataatgtccagcggctcttcgcggctgaggcgcgaaatg<br>taatgaacaatgatcatgttcatttcccggttccggcacatgtccaaaatccggttgaactccggccggtcgacgcccgtgaatgccgacgt<br>gccgggcgcttcgctgaagtgacccagccacttcacctcgacgccgtcgcgcgcgtactccttcgccagcgcctcagccttcccccggtt<br>cgcggcgcgctgggtggccggtgaagcggtcgaactgttctcccgttccgccgactgtcggtcgtagccgcctgccagaatgaccatgc<br>cttcaaccttcacgctcatgccgggcaggctatcccgcccaactgggtaagagcaaagactaacactgttgggctggaactactcccgaa<br>aaccgcttctgacctgggaaaacgtgaagccccggggcatccgctgagggttgccgccggggcttcggtgtgtccgtcagtacgggcc<br>atagaggggcgtcgtgtaacccacccagggcgattgctccggcacggggaagccgcgccacgccttcgggacgtctggaatcgctag<br>agcttgcatgc |
| pTSP | agtcagtatccagtcgtgtaggatcccgcaaaagcggcctttgactccctgcaagcctcagcgaccgaatatatcggttatgcgtgggcga<br>tggttgttgtcattgtcggcgcaactatcggtatcaagctgtttaagaaattcacctcgaaagcaagctgataaaccgatacaattaaaggct<br>ccttttggagcctttttttttggagattttcaacgtgaaaaaattattattcgcaattcctttagttgttcctttctattctcactccgctgaaactgttg<br>aaagttgtttagcaaaacctcatacagaaaattcatattaccctgttatccctactccgagacagtcagagggtagaattccttctagagcctc<br>agagacccggccgggtctcaggagtactagttcctgcagcactcgttgctttatcggtatattaccctgttatccctaaggaccaaaacgaa<br>aaaaggccccccctttcgggaggcctcttttctggaatttggtaccgagccataaactgccaggcatcagacgctcagtggaacgaaaactc<br>acgttaagggtaaaactgtattataagtaaatgcatgtatactaaactcacaaattagagcttcaatttaattatatcagttattacccataacttc<br>gtatagcatacattatacgaagttatcccgggtaccgagctcgattcgtaacttacacgcgcctcgtatcttttaatgatggaataatttgggaa<br>tttactctgtgtttatttatttttatgttttgtatttggattttagaaagtaaataaagaaggtagaagagttacggaatgaagaaaaaaaaaataaac<br>aaaggtttaaaaaatttcaacaaaaagcgtactttacatatatatttattagacaagaaaagcagattaaatagatatacattcgattaacgata<br>agtaaaatgtaaaatcacaggattttcgtgtgtggtcttctacacagacaagatgaaacaattcggcattaatacctgagagcaggaagagc<br>aagataaaaggtagtatttgttggcgatccccctagagtcttttacatcttcggaaaacaaaaactattttttctttaatttctttttttactttctatttt<br>taatttatatatttatattaaaaaatttaaattataattatttttatagcacgtgagcgctagcggagtgtatactggcttactatgttggcactgatg<br>agggtgtcagtgaagtgcttcatgtggcaggagaaaaaaggctgcaccggtgcgtcagcagaatatgtgatacaggatatattccgcttcc<br>tcgctcactgactcgctacgctcggtcgttcgactgcggcgagcggaaatggcttacgaacggggcggagatttcctggaagatgccag |

| | |
|---|---|
| | gaagatacttaacagggaagtgagagggccgcggcaaagccgtttttccataggctccgcccccctgacaagcatcacgaaatctgacg<br>ctcaaatcagtggtggcgaaacccgacaggactataaagataccaggcgtttccccctggcggctccctcgtgcgctctcctgttcctgcctt<br>tcggtttaccggtgtcattccgctgttatggccgcgtttgtctcattccacgcctgacactcagttccgggtaggcagttcgctccaagctgga<br>ctgtatgcacgaaccccccgttcagtccgaccgctgcgccttatccggtaactatcgtcttgagtccaacccggaaagacatgcaaaagca<br>ccactggcagcagccactggtaattgatttagaggagttagtcttgaagtcatgcgccggttaaggctaaactgaaaggacaagttttggtg<br>actgcgctcctccaagccagttacctcggttcaaagagttggtagctcagagaaccttcgaaaaaccgccctgcaaggcggttttttcgtttt<br>cagagcaagagattacgcgcagaccaaaacgatctcaagaagatcatcttattaagccagccaggacagaaatgcctcgacttcgctgct<br>acccaaggttgccgggtgacgcacaccgtggaaacggatgaaggcacgaacccagtggacataagcctgttcggttcgtaagctgtaat<br>gcaagtagcgtatgcgctcacgcaactggtccagaaccttgaccgaacgcagcggtggtaacggcgcagtggcggttttcatggcttgtt<br>atgactgtttttttggggtacagtctatgcctcgggcatccaagcagcaagcgcgttacgccgtgggtcgatgtttgatgttatggagcagca<br>acgatgttacgcagcagggcagtcgccctaaaacaaagttaaacattatgagggaagcggtgatcgccgaagtatcgactcaactatcag<br>aggtagttggcgtcatcgagcgccatctcgaaccgacgttgctggccgtacatttgtacggctccgcagtggatggcggcctgaagccac<br>acagtgatattgatttgctggttacggtgaccgtaaggcttgatgaaacaacgcggcgagctttgatcaacgaccttttggaaacttcggctt<br>cccctggagagagcgagattctccgcgctgtagaagtcaccattgttgtgcacgacgacatcattccgtggcgttatccagctaagcgcga<br>actgcaatttggagaatggcagcgcaatgacattcttgcaggtatcttcgagccagccacgatcgacattgatctggctatcttgctgacaa<br>aagcaagagaacatagcgttgccttggtaggtccagcggcggaggaactctttgatccggttcctgaacaggatctatttgaggcgctaaa<br>tgaaaccttaacgctatggaactcgccgcccgactgggctggcgatgagcgaaatgtagtgcttacgttgtcccgcatttggtacagcgca<br>gtaaccggcaaaatcgcgccgaaggatgtcgctgccgactgggcaatggagcgcctgccggcccagtatcagcccgtcatacttgaag<br>ctagacaggcttatcttggacaagaagaagatcgcttggcctcgcgcgcagatcagttggaagaatttgtccactacgtgaaaggcgaga<br>tcaccaaggtagtcggcaaataaccctcgagccacccatgaccaaaatcccttaacgtgagttacgcgtcgttccactgagcgtcagacg<br>gtccccggggatcggtcttgccttgctcgtcggtgatgtacttcaccagctccgcgaagtcgctcttcttgatggagcgcatggggacgtg<br>cttggcaatcacgcgcacccccggccgttttagcggctaaaaaagtcatggctctgccctcgggcggaccacgcccatcatgaccttgc<br>caagctcgtcctgcttctcttcgatcttcgccagcagggcgaggatcgtggcatcaccgaaccgcgccgtgcgcgggtcgtcggtgagc<br>cagagtttcagcaggccgcccaggcggcccaggtcgccattgatgcgggccagctcgcggacgtgctcatagtccacgacgcccgtg<br>attttgtagccctggccgacggccagcaggtaggccgacaggctcatgccggccgccgccgccttttcctcaatcgctcttcgttcgtctg<br>gaaggcagtacaccttgataggtgggctgcccttcctggttggcttggtttcatcagccatccgcttgccctcatctgttacgccggcggtag<br>ccggccagcctcgcagagcaggattcccgttgagcaccgccaggtgcgaataagggacagtgaagaaggaacacccgctcgcgggt<br>gggcctacttcacctatcctgcccggctgacgccgttggatacaccaaggaaagtctacacgaaccctttggcaaaatcctgtatatcgtgc<br>gaaaaaggatggatataccgaaaaaatcgctataatgaccccgaagcagggttatgcagcggaaaagatccgtcgacctgtcacgccgc<br>cgctgtgaacccgttcagcgtctccggctccgtctcttcctcgaccttcggcttcgcccagtggatacgcacacggtcggcgatcggggg<br>aacccgtccgggccgctgaacgcccttcggaagcttgatcacctcgatccggtccacgaagagccggacgaacgcccgacggtcttca<br>aggggcgctagcgcccaccacgaacccggtcccgtcgggtcgctgccccggtcgcccagccattcgccgatcggcagcacgggga<br>gtcggcggcgtcgagctgacgaagccgttcttcggccccttccatgcggagcgtcagcgcggcttcctcttcgagaaacgcgcgccgtc<br>ccatggcggaccggtagccgccgttccgcttgtcttcgtaaagctcttcgagcgccttcacggcgtccgcgcgctgtgccatcagttcgga<br>gcgctgacccttcaactccgggcgttcgtgcgtcgctccccagcgccgtgcggcttcgtacatgagcgctgccgtgtcgccttcgaggtc<br>gtcagggtcggcagggtcgaaggccgtgatgcgcgcgaagatcgcgccgacgatgtacgggtcaaggttgtgcatcgtgatcgagcat<br>gacgacccgtcatgaacccggcgggggcacttgcacgcgtacgacgacttgtgagcttgatcaccttcgcggacgttgcccgccatggt<br>cgacccgttgctgtaacccgtctccgggtcgagctggccggagccgtagcagtaaaggacgtccattgccgacaggagcgattgcccc<br>cggtactgacccttcccgcgtcctcgaccctgaagccactcctgaagttcccaccattccgccgggggaatgtacggctcgaagccggg<br>cagggtcagcggctccatggtgaccgggtcgcgcctgatcttgtaatggctgaagccgccccgcgaaccgtcggcgcgcaccttgtatg<br>cgatgtcagcttgatacccggcaatgcgcgggtcgctgagtacgcgcttcaaaacgccggggtcccaatcggaaccggcgcgcttctta<br>ccgaccagcgtgccgcgcgtaggcaccttgtcgcggtacagccgctcacaaagcccgttcagcgacccagggtgaaacgacccggct<br>ccgccacccttgaagggcgtatcgcggtgcgtcttgatctcctgccacgcccagcggattaccgccccttcgctgccgtgtgcgccttccc<br>aggtgtgcgcgctgggcaccagtcggcgaatggcaaccagctttccgccgtcttccgggttcggaaccatttcctcgaccgtgtcgaatc<br>cgtacggcgtcgaccccgtgtgtccgcccagccgcttcgccaattccttagcgttcgacacggcgacgctcttgttcttcgactcatcatgc<br>gaagcctgaaggcgcatgatcaggtgaataaggtccatcatttcgccgggggcggaatgtgccttcgttcacgctgacaatggtcacgccc<br>agccggagcaattccgtgacgaccggaataatgtccagcggctcttcgcggctgaggcgcgaaatgtaatgaacaatgatcatgttcattt<br>cccggttccggcacatgtccaaaatccggttgaactccggccggtcgacgcccgtgaatgccgacgtgccgggcgcttcgctgaagtga<br>cccagccacttcacctcgacgccgtcgcgcgcgtactccttcgccagcgcctcagccttcccccggttcgcggcgcgctgggtggccgg<br>tgaagcggtcgaactgttctcccgttccgccgactgtcggtcgtagccgcctgccagaatgaccatgccttcaaccttcacgctcatgccg<br>ggcaggctatcccgcccaactgggtaagagcaaagactaacactgttgggctggaactactcccgaaaaccgcttctgacctgggaaaa<br>cgtgaagccccggggcatccgctgagggttgccgccggggcttcggtgtgtccgtcagtacgggccatagaggggcgtcgtgtaaccc<br>acccagggcgattgctccggcacggggaagccgcgccacgccttcgggacgtctggaatcgctagagcttgcatgc |
| pIATP | acgaaacctacgataagagtggatcccgcaaaagcggcctttgactccctgcaagcctcagcgaccgaatatatcggttatgcgtgggcg<br>atggttgttgtcattgtcggcgcaactatcggtatcaagctgtttaagaaattcacctcgaaagcaagctgataaaccgatacaattaaaggc<br>tccttttggagcctttttttttggagattttcaacgtgaaaaaattattattcgcaattcctttagttgttcctttctattctcactccgctgaaactgttg<br>aaagttgtttagcaaaacctcatacagaaaattcatattaccctgttatccctactccgagacagtcagagggtagaattccttctagagcctc<br>agagacccggccgggtctcaggagtactagttcctgcagcactcgttgctttatcggtatattaccctgttatccctatttttgttgctataaaaa<br>aaggcccccccgatttgggaggcctttttttcgaaaatgccgactcactattacagacgacgctcagtggaacgaaaactcacgttaagggta<br>aaactgtattataagtaaatgcatgtatactaaactcacaaattagagcttcaatttaattatatcagttattacccataacttcgtatagcatacat<br>tatacgaagttatcccgggtaccgagctcgattcgtaacttacacgcgcctcgtatcttttaatgatggaataatttgggaatttactctgtgttt<br>atttatttttatgttttgtatttggattttagaaagtaaataaagaaggtagaagagttacggaatgaagaaaaaaaaaataaacaaaggtttaaaa<br>aatttcaacaaaaagcgtactttacatatatatttattagacaagaaaagcagattaaatagatatacattcgattaacgataagtaaaatgtaa<br>aatcacaggattttcgtgtgtggtcttctacacagacaagatgaaacaattcggcattaatacctgagagcaggaagagcaagataaaagg<br>tagtatttgttggcgatccccctagagtcttttacatcttcggaaaacaaaaactattttttctttaatttctttttttactttctatttttaatttatatattt<br>atattaaaaaatttaaattataattatttttatagcacgtgagcgctagcggagtgtatactggcttactatgttggcactgatgagggtgtcagt<br>gaagtgcttcatgtggcaggagaaaaaaggctgcaccggtgcgtcagcagaatatgtgatacaggatatattccgcttcctcgctcactga |

ctcgctacgctcggtcgttcgactgcggcgagcggaaatggcttacgaacggggcggagatttcctggaagatgccaggaagatactta
acagggaagtgagagggccgcggcaaagccgtttttccataggctccgcccccctgacaagcatcacgaaatctgacgctcaaatcagt
ggtggcgaaacccgacaggactataaagataccaggcgtttccccctggcggctccctcgtgcgctctcctgttcctgcctttcggtttaccg
gtgtcattccgctgttatggccgcgtttgtctcattccacgcctgacactcagttccgggtaggcagttcgctccaagctggactgtatgcac
gaaccccccgttcagtccgaccgctgcgccttatccggtaactatcgtcttgagtccaacccggaaagacatgcaaaagcaccactggca
gcagccactggtaattgatttagaggagttagtcttgaagtcatgcgccggttaaggctaaactgaaaggacaagttttggtgactgcgctc
ctccaagccagttacctcggttcaaagagttggtagctcagagaaccttcgaaaaaccgccctgcaaggcggttttttcgttttcagagcaa
gagattacgcgcagaccaaaacgatctcaagaagatcatcttattaaggatggcctttttgcgtttctacaaactcttttgtttatttttctaaata
cattcaaatatgtatccgctcatgagacaataaccctgataaatgcttcaataatattgaaaaaggaagactatgagtattcaacatttccgtgt
cgcccttattcccttttttgcggcattttgccttcctgtttttgctcacccagaaacgctggtgaaagtaaaagatgctgaagatcagttgggtg
cacgagtgggttacatcgaactggatctcaacagcggtaagatccttgagagttttcgccccgaagaacgttttccaatgatgagcactttta
aagttctgctatgtggcgcggtattatcccgtgttgacgccgggcaagagcaactcggtcgccgcatacactattctcagaatgacttggtt
gagtactcaccagtcacagaaaagcatcttacggatggcatgacagtaagagaattatgcagtgctgccataaccatgagtgataacactg
cggccaacttacttctgacaacgatcggaggaccgaaggagctaaccgcttttttgcacaacatgggggatcatgtaactcgccttgatcgt
tgggaaccggagctgaatgaagccataccaaacgacgagcgtgacaccacgatgccagcagcaatggcaacaacgttgcgcaaacta
ttaactggcgaactacttactctagcttcccggcaacaattaatagactggatggaggcggataaagttgcaggaccacttctgcgctcggc
ccttccggctggctggtttattgctgataaatctggagccggtgagcgtggatctcgcggtatcattgcagcactggggccagatggtaag
ccctcccgtatcgtagttatctacacgacggggagtcaggcaactatggatgaacgaaatagacagatcgctgagataggtgcctcactg
attaagcattggtaactgtcagaccaagtttactcatatatactttagattgatttacgcgccctgtagcggcgcattaagcgcggcgggtgtg
gtggttacgcgcagcgtgaccgctacacttgccagcgccctagcgcccgctcctttcgctttcttcccttcctttctcgccacgttcgccggc
tttccccgtcaacgcctgatgcggtattttctccttacgcatctgtgcggtatttcacaccgcatatgcggggatcgaccgcgcgggtcccg
gacggggaagagcggggagctttgccagagagcgacgacttccccttgcgttggtgattgccggtcagggcagccatccgccatcgtc
gcgtagggtgtcacaccccaggaatcgcgtcactgaacacagcagccggtaggacgaccatgactgagttggacaccatcgcaaatcc
gtccgatcccgcggtgcagcggatcatcgatgtcaccaagccgtcgcgatccaacataaagacaacgttgatcgaggacgtcgagccc
ctcatgcacagcatcgcggccggggtggagttcatcgaggtctacggcagcgacagcagtccttttccatctgagttgctggatctgtgcg
ggcggcagaacataccggtccgcctcatcgactcctcgatcgtcaaccagttgttcaaggggggagcggaaggccaagacattcggcatc
gcccgcgtccctcgcccggccaggttcggcgatatcgcgagccggcgtggggacgtcgtcgttctcgacggggtgaagatcgtcggg
aacatcggcgcgatagtacgcacgtcgctcgcgctcggagcgtcggggatcatcctggtcgacagtgacatcaccagcatcgcggacc
ggcgtctccaaagggccagccgaggttacgtcttctcccttcccgtcgttctctccggtcgcgaggaggccatcgccttcattcgggacag
cggtatgcagctgatgacgctcaaggcggatggcgacatttccgtgaaggaactcggggacaatccggatcggctggccttgctgttcg
gcagcgaaaagggtgggccttccgacctgttcgaggaggcgtcttccgcctcggtttccatccccatgatgagccagaccgagtctctca
acgtttccgtttccctcggaatcgcgctgcacgagaggatcgacaggaatctcgcggccaaccgataagcgcctctgttcctcggacgct
cggttcctcgacctcgattcgtcagtgatgatcggccgttgtgggctcaccgggctagctagagtcgacctgcggtccccggggatcggt
cttgccttgctcgtcggtgatgtacttcaccagctccgcgaagtcgctcttcttgatggagcgcatggggacgtgcttggcaatcacgcgca
ccccccggccgttttagcggctaaaaaagtcatggctctgccctcgggcggaccacgcccatcatgaccttgccaagctcgtcctgcttct
cttcgatcttcgccagcagggcgaggatcgtggcatcaccgaaccgcgccgtgcgcgggtcgtcggtgagccagagtttcagcaggcc
gcccaggcggcccaggtcgccattgatgcgggccagctcgcggacgtgctcatagtccacgacgcccgtgattttgtagccctggccga
cggccagcaggtaggccgacaggctcatgccggccgccgccgccttttcctcaatcgctcttcgttcgtctggaaggcagtacaccttgat
aggtgggctgcccttcctggttggcttggtttcatcagccatccgcttgccctcatctgttacgccggcggtagccggccagcctcgcaga
gcaggattcccgttgagcaccgccaggtgcgaataagggacagtgaagaaggaacacccgctcgcgggtgggcctacttcacctatcc
tgcccggctgacgccgttggatacaccaaggaaagtctacacgaaccctttggcaaaatcctgtatatcgtgcgaaaaaggatggatatac
cgaaaaaatcgctataatgaccccgaagcagggttatgcagcggaaaagatccgtcgacctgccgcacgtcaaagccccgccggatca
ccggcggggctctcttcggccctccaagtcacaccagccccaaggggcgtcgggagtggcggagggaacctctggcccgattggtgc
caggattcccaccagaccaaagagcaacgggccggacttcgcacctccgacccgtccgctcccagactcgcgccccttagccgggcg
agacaggaacgttgctcgtgcccagagtacggagcgatgccgaggcattgccagatcggcccgccgggccccgctgccactgcggg
accgcaattgcccacacaccgggcaaacggccgcgtatctactgctcagaccgctgccggatggcagcgaagcgggcgatcgcgcgt
gtgacgcgagatgccgcccgaggcaaaagcgaacaccttgggaaagaaacaacagagtttcccgcacccctccgacctgcggtttctc
cggacggggtggatggggagagcccgagaggcgacagcctctcggaagtaggaagcacgtcgcggagcgacgctgcccgactgcg
gaaagccgcccggtacagccgccgccggacgctgtggcggatcagcggggacgccgcgtgcaagggctgcggccgcgccctgatg
gaccctgcctccggcgtgatcgtcgcccagacggcggccggaacgtccgtggtcctgggcctgatgcggtgcgggcggatctggctct
gcccggtctgcgccgccacgatccggcacaagcgggccgaggagatcaccgccgccgtggtcgagtggatcaagcgcgggggggac
cgcctacctggtcaccttcacggcccgccatgggcacacggaccggctcgcggacctcatggacgccctccagggcacccggaagac
gccggacagcccccggcggccgggcgcctaccagcgactgatcacgggcggcacgtgggccggacgccgggccaaggacgggc
accgggccgccgaccgcgagggcatccgggaccggatcgggtacgtcggcatgatccgcgcgaccgaagtcaccgtggggcagat
caacggctggcacccgcacatccacgcgatcgtcctggtcggcggccggaccgaggggggagcggtccgcgaagcagatcgtcgcca
ccttcgagccgaccggcgccgcgctcgacgagtggcaggggcactggcggtccgtgtggaccgccgccctgcgcaaggtcaacccc
gccttcacgcccgacgaccggcacggcgtcgacttcaagcggctggagacggagcgcgacgccaacgacctcgccgagtacatcgc
caagacccaggacgggaaggcgcccgccctcgaactcgcccgcgccgacctcaagacggcgaccggcgggaacgtcgccccgttc
gaactcctcggacggatcggggacctgaccggcggcatgaccgaggacgacgccgccggggtcggctcgctggagtggaacctctc
gcgctggcacgagtacgagcgggcaacccggggacgccgggccatcgaatggacccgctacctgcggcagatgctcgggctcgac
ggcggcgacaccgaggccgacgacctcgatctgctcctggcggccgacgccgacggcggggagctgcgggccggggtcgccgtga
ccgaggacggatggcacgcggtcacccgccgcgccctcgacctcgaggcgacccgggccgccgaaggcaaggacggcaacgagg
attcggcggccgtgggcgaacgggtgcgggaggtcctggcgctggccgacgcggccgacacagtggtggtgctcacggcgggggga
ggtggccgaggcgtacgccgacatgctcgccgccctcgcccagcgccgcgaggaagcaactgcacgccgacggcgagagcaggac
gacgaccaggacgacgacgccgacgaccgccaggagcgggccgcccggcacatcgcccggctcgcaagtgggcccacttcgcact
aactcgctccccccgccgtacgtcatcccggtgacgtacggcgggggtcggtgacgtacgcggcgacggcggccggggtcgaagc
cgcgggagtaatcctgggattactcgcccggggtcggccccgccggcacttcgtgcaggcggta

| | |
|---|---|
| pIATU | acgaaacctacgataagagtggatcccgcaaaagcggcctttgactccctgcaagcctcagcgaccgaatatatcggttatgcgtgggcg<br>atggttgttgtcattgtcggcgcaactatcggtatcaagctgtttaagaaattcacctcgaaagcaagctgataaaccgatacaattaaaggc<br>tcctttggagcctttttttttggagattttcaacgtgaaaaaattattattcgcaattcctttagttgttcctttctattctcactccgctgaaactgttg<br>aaagttgtttagcaaaacctcatacagaaaattcatattaccctgttatccctactccgagacagtcagagggtagaattccttctagagcctc<br>agagacccggccgggtctcaggagtactagttcctgcagcactcgttgctttatcggtatattaccctgttatccctatttttgttgctataaaaa<br>aaggcccccgatttgggaggcctttttcgaaaagctcactcaaaggcggtaatgacgctcagtggaacgaaaactcacgttaagggta<br>aaactgtattataagtaaatgcatgtatactaaactcacaaattagagcttcaatttaattatatcagttattacccataacttcgtatagcatacat<br>tatacgaagttatcccgggtaccgagctcgattcgtaacttacacgcgcctcgtatcttttaatgatggaataatttgggaatttactctgtgttt<br>atttatttttatgttttgtatttggattttagaaagtaaataaagaaggtagaagagttacggaatgaagaaaaaaaaataaacaaaggtttaaaa<br>aatttcaacaaaaagcgtactttacatatatatttattagacaagaaaagcagattaaatagatatacattcgattaacgataagtaaaatgtaa<br>aatcacaggattttcgtgtgtggtcttctacacagacaagatgaaacaattcggcattaatacctgagagcaggaagagcaagataaaagg<br>tagtatttgttggcgatccccctagagtcttttacatcttcggaaaacaaaaactatttttctttaatttctttttttactttctatttttaatttatatattt<br>atattaaaaaatttaaattataattatttttatagcacgtgagcgctagcggagtgtatactcggttatccacagaatcaggggataacgcagg<br>aaagaacatgtgagcaaaaggccagcaaaaggccaggaaccgtaaaaaggccgcgttgctggcgtttttccacaggctccgcccccct<br>gacgagcatcacaaaaatcgacgctcaagtcagaggtggcgaaacccgacaggactataaagataccaggcgtttccccctggaagct<br>ccctcgtgcgctctcctgttccgaccctgccgcttaccggatacctgtccgcctttctcccttcgggaagcgtggcgctttctcatagctcac<br>gctgtaggtatctcagttcggtgtaggtcgttcgctccaagctgggctgtgtgcacgaaccccccgttcagcccgaccgctgcgccttatcc<br>ggtaactatcgtcttgagtccaacccggtaagacacgacttatcgccactggcagcagccactggtaacaggattagcagagcgaggtat<br>gtaggcggtgctacagagttcttgaagtggtggcctaactacggctacactagaagaacagtatttggtatctgcgctctgctgaagccagt<br>taccttcggaaaaagagttggtagctcttgatccggcaaacaaaccaccgctggtagcggtggtttttttgtttgcaagcagcagattacgc<br>gcagaaaaaaaggatctcaagaagatcctttgatcttttctacggggatggcctttttgcgtttctacaaactcttttgtttatttttctaaatacatt<br>caaatatgtatccgctcatgagacaataaccctgataaatgcttcaataatattgaaaaaggaagactatgagtattcaacatttccgtgtcgc<br>ccttattcccttttttgcggcattttgccttcctgtttttgctcacccagaaacgctggtgaaagtaaaagatgctgaagatcagttgggtgcac<br>gagtgggttacatcgaactggatctcaacagcggtaagatccttgagagttttcgccccgaagaacgttttccaatgatgagcacttttaaag<br>ttctgctatgtggcgcggtattatcccgtgttgacgccgggcaagagcaactcggtcgccgcatacactattctcagaatgacttggttgagt<br>actcaccagtcacagaaaagcatcttacggatggcatgacagtaagagaattatgcagtgctgccataaccatgagtgataacactgcgg<br>ccaacttacttctgacaacgatcggaggaccgaaggagctaaccgcttttttgcacaacatgggggatcatgtaactcgccttgatcgttgg<br>gaaccggagctgaatgaagccataccaaacgacgagcgtgacaccacgatgccagcagcaatggcaacaacgttgcgcaaactattaa<br>ctggcgaactacttactctagcttcccggcaacaattaatagactggatggaggcggataaagttgcaggaccacttctgcgctcggccctt<br>ccggctggctggtttattgctgataaatctggagccggtgagcgtggatctcgcggtatcattgcagcactggggccagatggtaagccct<br>cccgtatcgtagttatctacacgacggggagtcaggcaactatggatgaacgaaatagacagatcgctgagataggtgcctcactgattaa<br>gcattggtaactgtcagaccaagtttactcatatatactttagattgatttacgcgccctgtagcggcgcattaagcgcggcgggtgtggtgg<br>ttacgcgcagcgtgaccgctacacttgccagcgccctagcgcccgctcctttcgctttcttcccttcctttctcgccacgttcgccggctttcc<br>ccgtcaacgcctgatgcggtattttctccttacgcatctgtgcggtatttcacaccgcatatgcggggatcgaccgcgcgggtcccggacg<br>gggaagagcggggagctttgccagagagcgacgacttccccttgcgttggtgattgccggtcagggcagccatccgccatcgtcgcgta<br>gggtgtcacaccccaggaatcgcgtcactgaacacagcagccggtaggacgaccatgactgagttggacaccatcgcaaatccgtccg<br>atcccgcggtgcagcggatcatcgatgtcaccaagccgtcgcgatccaacataaagacaacgttgatcgaggacgtcgagcccctcatg<br>cacagcatcgcggccggggtggagttcatcgaggtctacggcagcgacagcagtccttttccatctgagttgctggatctgtgcgggcgg<br>cagaacataccggtccgcctcatcgactcctcgatcgtcaaccagttgttcaagggggagcggaaggccaagacattcggcatcgcccg<br>cgtccctcgcccggccaggttcggcgatatcgcgagccggcgtggggacgtcgtcgttctcgacggggtgaagatcgtcgggaacatc<br>ggcgcgatagtacgcacgtcgctcgcgctcggagcgtcggggatcatcctggtcgacagtgacatcaccagcatcgcggaccggcgtc<br>tccaaagggccagccgaggttacgtcttctcccttcccgtcgttctctccggtcgcgaggaggccatcgccttcattcgggacagcggtat<br>gcagctgatgacgctcaaggcggatggcgacatttccgtgaaggaactcggggacaatccggatcggctggccttgctgttcggcagcg<br>aaaagggtgggccttccgacctgttcgaggaggcgtcttccgcctcggtttccatccccatgatgagccagaccgagtctctcaacgtttc<br>cgtttccctcggaatcgcgctgcacgagaggatcgacaggaatctcgcggccaaccgataagcgcctctgttcctcggacgctcggttcc<br>tcgacctcgattcgtcagtgatgatcggccgttgtgggctcaccgggctagctagagtcgacctgcggtccccggggatcggtcttgcctt<br>gctcgtcggtgatgtacttcaccagctccgcgaagtcgctcttcttgatggagcgcatggggacgtgcttggcaatcacgcgcaccccccc<br>ggccgttttagcggctaaaaaagtcatggctctgccctcgggcggaccacgcccatcatgaccttgccaagctcgtcctgcttctcttcgat<br>cttcgccagcagggcgaggatcgtggcatcaccgaaccgcgccgtgcgcgggtcgtcggtgagccagagtttcagcaggccgcccag<br>gcggcccaggtcgccattgatgcgggccagctcgcggacgtgctcatagtccacgacgcccgtgattttgtagccctggccgacggcca<br>gcaggtaggccgacaggctcatgccggccgccgccgccttttcctcaatcgctcttcgttcgtctggaaggcagtacaccttgataggtgg<br>gctgcccttcctggttggcttggtttcatcagccatccgcttgccctcatctgttacgccggcggtagccggccagcctcgcagagcaggat<br>tcccgttgagcaccgccaggtgcgaataagggacagtgaagaaggaacacccgctcgcgggtgggcctacttcacctatcctgcccgg<br>ctgacgccgttggatacaccaaggaaagtctacacgaaccctttggcaaaatcctgtatatcgtgcgaaaaaggatggatataccgaaaa<br>aatcgctataatgaccccgaagcagggttatgcagcggaaaagatccgtcgacctgccgcacgtcaaagccccgccggatcaccggcg<br>gggctctcttcggccctccaagtcacaccagccccaaggggcgtcgggagtggcggagggaacctctggcccgattggtgccaggatt<br>cccaccagaccaaagagcaacgggccggacttcgcacctccgacccgtccgctcccagactcgcgccccttagccgggcgagacag<br>gaacgttgctcgtgcccagagtacggagcgatgccgaggcattgccagatcggcccgccgggccccgctgccactgcgggaccgcaa<br>ttgcccacacaccgggcaaacggccgcgtatctactgctcagaccgctgccggatggcagcgaagcgggcgatcgcgcgtgtgacgc<br>gagatgccgcccgaggcaaaagcgaacaccttgggaaagaaacaacagagtttcccgcacccctccgacctgcggtttctccggacgg<br>ggtggatggggagagcccgagaggcgacagcctctcggaagtaggaagcacgtcgcggagcgacgctgcccgactgcggaaagcc<br>gcccggtacagccgccgccggacgctgtggcggatcagcggggacgccgcgtgcaagggctgcggccgcgccctgatggaccctg<br>cctccggcgtgatcgtcgcccagacggcggccggaacgtccgtggtcctgggcctgatgcggtgcgggcggatctggctctgcccggt<br>ctgcgccgccacgatccggcacaagcgggccgaggagatcaccgccgccgtggtcgagtggatcaagcgcgggggggaccgcctac<br>ctggtcaccttcacggcccgccatgggcacacggaccggctcgcggacctcatggacgccctccagggcacccggaagacgccgga<br>cagcccccggcggccgggcgcctaccagcgactgatcacgggcggcacgtgggccggacgccgggccaaggacgggcaccggg<br>ccgccgaccgcgagggcatccgggaccggatcgggtacgtcggcatgatccgcgcgaccgaagtcaccgtggggcagatcaacggc |

| | |
|---|---|
| | tggcacccgcacatccacgcgatcgtcctggtcggcggccggaccgagggggagcggtccgcgaagcagatcgtcgccaccttcga<br>gccgaccggcgccgcgctcgacgagtggcaggggcactggcggtccgtgtggaccgccgccctgcgcaaggtcaaccccgccttca<br>cgcccgacgaccggcacggcgtcgacttcaagcggctggagacggagcgcgacgccaacgacctcgccgagtacatcgccaagac<br>ccaggacgggaaggcgcccgccctcgaactcgcccgcgccgacctcaagacggcgaccggcgggaacgtcgccccgttcgaactc<br>ctcggacggatcggggacctgaccggcggcatgaccgaggacgacgccgccggggtcggctcgctggagtggaacctctcgcgctg<br>gcacgagtacgagcgggcaacccggggacgccgggccatcgaatggacccgctacctgcggcagatgctcgggctcgacggcggc<br>gacaccgaggccgacgacctcgatctgctcctggcggccgacgccgacggcggggagctgcgggccggggtcgccgtgaccgagg<br>acggatggcacgcggtcacccgccgcgccctcgacctcgaggcgacccgggccgccgaaggcaaggacggcaacgaggattcggc<br>ggccgtgggcgaacgggtgcgggaggtcctggcgctggccgacgcggccgacacagtggtggtgctcacggcgggggaggtggcc<br>gaggcgtacgccgacatgctcgccgccctcgcccagcgccgcgaggaagcaactgcacgccgacggcgagagcaggacgacgac<br>caggacgacgacgccgacgaccgccaggagcgggccgcccggcacatcgcccggctcgcaagtgggcccacttcgcactaactcg<br>ctcccccccgccgtacgtcatcccggtgacgtacggcgggggtcggtgacgtacgcggcgacggcggccggggtcgaagccgcgg<br>gagtaatcctgggattactcgcccggggtcggccccgccggcacttcgtgcaggcggta |
| pPAS-PT | tgccacctgacgtctaagaagatcccgcaaaagcggcctttgactccctgcaagcctcagcgaccgaatatatcggttatgcgtgggcga<br>tggttgttgtcattgtcggcgcaactatcggtatcaagctgtttaagaaattcacctcgaaagcaagctgataaaccgatacaattaaaggct<br>ccttttggagcctttttttttggagattttcaacgtgaaaaaattattattcgcaattcctttagttgttcctttctattctcactccgctgaaactgttg<br>aaagttgtttagcaaaacctcatacagaaaattcatattaccctgttatccctactccgagacagtcagagggtagaattcgcggccgcttct<br>agagctcggtaccaaattccagaaaagaggccgcgaaagcggcctttttttcgttttggtcctactagatgcctccacaccgctcgtcacatc<br>ctgggagaccttaccattcgccattcaggctgcgcaactgttgggaagggcgatcggtgcgggcctcttcgctattacgccagctggcga<br>aagggggatgtgctgcaaggcgattaagttgggtaacgccagggttttcccagtcacgacgttgtaaaacgacggccagtgaatccgtaa<br>tcatggtcatagctgtttcctgtgtgaaattgttatccgctcacaattccacacaacatacgagccggaagcataaagtgtaaagcctggggt<br>gcctaatgagtgagctaactcacattaattgcgttgcgcggtctcaagctgtcaccggatgtgctttccggtctgatgagtccgtgaggacg<br>aaacagcctctacaaataattttgtttaatactagagaaagaggagaaatactagatgcgtaaaggcgaagagctgttcactggtgtcgtcc<br>ctattctggtggaactggatggtgatgtcaacggtcataagttttccgtgcgtggcgagggtgaaggtgacgcaactaatggtaaactgac<br>gctgaagttcatctgtactactggtaaactgccggtaccttggccgactctggtaacgacgctgacttatggtgttcagtgctttgctcgttatc<br>cggaccatatgaagcagcatgacttcttcaagtccgccatgccggaaggctatgtgcaggaacgcacgatttcctttaaggatgacggca<br>cgtacaaaacgcgtgcggaagtgaaatttgaaggcgataccctggtaaaccgcattgagctgaaaggcattgactttaaagaagacggc<br>aatatcctgggccataagctggaatacaattttaacagccacaatgtttacatcaccgccgataaacaaaaaaatggcattaaagcgaatttt<br>aaaattcgccacaacgtggaggatggcagcgtgcagctggctgatcactaccagcaaaacactccaatcggtgatggtcctgttctgctg<br>ccagacaatcactatctgagcacgcaaagcgttctgtctaaagatccgaacgagaaacgcgatcatatggttctgctggagttcgtaaccg<br>cagcgggcatcacgcatggtatggatgaactgtacaaatgatgataagccaggcatcaaataaaacgaaaggctcagtcgaaagactgg<br>gcctttcgttttatctgttgtttgtcggtgaacgctctctactagagtcacactggctcaccttcgggtgggcctttctgcgtttatatactagagc<br>tgctaacaaagcccgaaaggaagctgagttggctgctgccaccgctgagcaataactagcataaccccttggggcctctaaacgggtctt<br>gaggggttttttgctgaaaggaggaactatatccggattactagaggtcatgcttgccatctgttttcttgcaagattactagttcctgcagcac<br>tcgttgctttatcggtatattaccctgttatccctaattttgttatcaataaaaaaggccccccgttagggaggccttattgttcgtcgctcactca<br>aaggcggtaatgacgctcagtggaacgaaaactcacgttaagggtaaaactgtattataagtaaatgcatgtatactaaactcacaaattag<br>agcttcaatttaattatatcagttattacccataacttcgtatagcatacattatacgaagttatcccgggtaccgagctcgattcgtaacttacac<br>gcgcctcgtatcttttaatgatggaataatttgggaatttactctgtgtttatttatttttatgttttgtatttggattttagaaagtaaataaagaaggt<br>agaagagttacggaatgaagaaaaaaaaaataaacaaaggtttaaaaaatttcaacaaaaagcgtactttacatatatatttattagacaagaa<br>aagcagattaaatagatatacattcgattaacgataagtaaaatgtaaaatcacaggattttcgtgtgtggtcttctacacagacaagatgaaa<br>caattcggcattaatacctgagagcaggaagagcaagataaaaggtagtatttgttggcgatccccctagagtcttttacatcttcggaaaac<br>aaaaactatttttctttaatttctttttttactttctatttttaatttatatatttatattaaaaaatttaaattataattatttttatagcacgtgagctagag<br>ctgtcagaccaagtttacgagctcgcttggactcctgttgatagatccagtaatgacctcagaactccatctggatttgttcagaacgctcggt<br>tgccgccgggcgttttttattggtgagaatccaagcactagggacagtaagacgggtaagcctgttgatgataccgctgccttactgggtgc<br>attagccagtctgaatgacctgtcacgggataatccgaagtggtcagactggaaaatcagagggcaggaactgctgaacagcaaaaagt<br>cagatagcaccacatagcagacccgccataaaacgccctgagaagcccgtgacgggcttttcttgtattatgggtagtttccttgcatgaat<br>ccataaaaggcgcctgtagtgccatttacccccattcactgccagagccgtgagcgcagcgaactgaatgtcacgaaaaagacagcgac<br>tcaggtgcctgatggtcggagacaaaaggaatattcagcgatttgcccgagcttgcgagggtgctacttaagcctttagggttttaaggtct<br>gttttgtagaggagcaaacagcgtttgcgacatccttttgtaatactgcggaactgactaaagtagtgagttatacacagggctgggatctatt<br>ctttttatctttttttattctttctttattctataaattataaccacttgaatataaacaaaaaaaacacacaaaggtctagcggaatttacagagggt<br>ctagcagaatttacaagttttccagcaaaggtctagcagaatttacagatacccacaactcaaaggaaaaggacatgtaattatcattgacta<br>gcccatctcaattggtatagtgattaaaatcacctagaccaattgagatgtatgtctgaattagttgttttcaaagcaaatgaactagcgattagt<br>cgctatgacttaacggagcatgaaaccaagctaattttatgctgtgtggcactactcaaccccacgattgaaaaccctacaaggaaagaac<br>ggacggtatcgttcacttataaccaatacgctcagatgatgaacatcagtagggaaaatgcttatggtgtattagctaaagcaaccagagag<br>ctgatgacgagaactgtggaaatcaggaatcctttggttaaaggctttgagattttccagtggacaaactatgccaagttctcaagcgaaaa<br>attagaattagtttttagtgaagagatattgccttatcttttccagttaaaaaaattcataaaatataatctggaacatgttaagtcttttgaaaaca<br>aatactctatgaggatttatgagtggttattaaaagaactaacacaaaagaaaactcacaaggcaaatatagagattagccttgatgaatttaa<br>gttcatgttaatgcttgaaaataactaccatgagtttaaaaggcttaaccaatgggttttgaaaccaataagtaaagatttaaacacttacagca<br>atatgaaattggtggttgataagcgaggccgcccgactgatacgttgattttccaagttgaactagatagacaaatggatctcgtaaccgaa<br>cttgagaacaaccagataaaaatgaatggtgacaaaataccaacaaccattacatcagattcctacctacgtaacggactaagaaaaacac<br>tacacgatgctttaactgcaaaaattcagctcaccagttttgaggcaaaatttttgagtgacatgcaaagtaagcatgatctcaatggttcgttc<br>tcatggctcacgcaaaaacaacgaaccacactagagaacatactggctaaatacggaaggatctgaggttcttatggctcttgtatctatca<br>gtgaagcatcaagactaacaaacaaaagtagaacaactgttcaccgttagatatcaaagggaaaactgtccatatgcacagatgaaaacg<br>gtgtaaaaaagatagatacatcagagcttttacgagtttttggtgcatttaaagctgttcaccatgaacagatcgacaatgtaactactagaga<br>ttttggtcatgagattatcaaaaaggatcttcacctagatccttttggttcatgtgcagctccatcagcaaaaggggatgataagtttatcacca<br>ccgactatttgcaacagtgccgttgatcgtgctatgatcgactgatgtcatcagcggtggagtgcaatgtcgtgcaatacgaatggcgaaaa<br>gccgagctcatcggtcagcttctcaaccttggggttaccccggcggtgtgctgctggtccacagctccttccgtagcgtccggcccctcg |

| | |
|---|---|
| | aagatgggccacttggactgatcgaggccctgcgtgctgcgctgggtccgggagggacgctcgtcatgccctcgtggtcaggtctggac<br>gacgagccgttcgatcctgccacgtcgcccgttacaccggaccttggagttgtctctgacacattctggcgcctgccaaatgtaaagcgca<br>gcgcccatccatttgcctttgcggcagcggggccacaggcagagcagatcatctctgatccattgcccctgccacctcactcgcctgcaa<br>gcccggtcgcccgtgtccatgaactcgatgggcaggtacttctcctcggcgtgggacacgatgccaacacgacgctgcatcttgccgagt<br>tgatggcaaaggttccctatggggtgccgagacactgcaccattcttcaggatggcaagttggtacgcgtcgattatctcgagaatgacca<br>ctgctgtgagcgctttgccttggcggacaggtggctcaaggagaagagccttcagaaggaaggtccagtcggtcatgcctttgctcggtt<br>gatccgctcccgcgacattgtggcgacagccctgggtcaactgggccgagatccgttgatcttcctgcatccgccagaggcgggatgcg<br>aagaatgcgatgccgctcgccagtcgattggctgagctcatgagcggagaacgagatgacgttggaggggcaaggtcgcgctgattgc<br>tggggcaacacgtggagcggatcggggattgtctttcttcagctcgctgatgatatgctgacgctcaatgccgtttggcctccgactaacga<br>aaatcccgcatttggacggctgatccgattggcacggcggacggcgaatggcggagcagacgctcgtccgggggcaatgagatatga<br>aaaagcctgaactcaccgcgacgtatcgggccctggccagctagctagagtcgacctgggtccccggggatcggtcttgccttgctcgtc<br>ggtgatgtacttcaccagctccgcgaagtcgctcttcttgatggagcgcatggggacgtgcttggcaatcacgcgcacccccggccgttt<br>tagcggctaaaaaagtcatggctctgccctcgggcggaccacgcccatcatgaccttgccaagctcgtcctgcttctcttcgatcttcgcca<br>gcagggcgaggatcgtggcatcaccgaaccgcgccgtgcgcgggtcgtcggtgagccagagtttcagcaggccgcccaggcggccc<br>aggtcgccattgatgcgggccagctcgcggacgtgctcatagtccacgacgcccgtgattttgtagccctggccgacggccagcaggta<br>ggccgacaggctcatgccggccgccgccgccttttcctcaatcgctcttcgttcgtctggaaggcagtacaccttgataggtgggctgccc<br>ttcctggttggcttggtttcatcagccatccgcttgccctcatctgttacgccggcggtagccggccagcctcgcagagcaggattcccgtt<br>gagcaccgccaggtgcgaataagggacagtgaagaaggaacacccgctcgcgggtgggcctacttcacctatcctgcccggctgacg<br>ccgttggatacaccaaggaaagtctacacgaaccctttggcaaaatcctgtatatcgtgcgaaaaaggatggatataccgaaaaaatcgct<br>ataatgaccccgaagcagggttatgcagcggaaaagatccgtcgacctgctacgccgctacgtcttccgtgccgtcctgggcgtcgtcttc<br>gtcgtcgtcggtcggcggcttcgcccacgtgatcgaagcgcgcttctcgatgggcgttccctgccccctgcccgtagtcgacttcgtgaca<br>acgatcttgtctacgaagagcccgacgaacacgcgcttgtcgtctactgacgcgcgcccccaccacgacttagggccggtcgggtcagc<br>gtcggcgtcttcggggaaccattggtcaaggggaagcttcggggcttcggcggcttcaagttcggcaagccgctcttccgccccttgctg<br>ccggagcgtcagcgctgcctgttgcttccggaagtgcttcctgccaacgggtccgtcgtacgcgcctgccgcgcggtcttcgtacagctct<br>tcaagggcgttcagggcgtcggcgcgctccgcaacaaggttcgcccgttcgccgctcttctcaggcgcctcagtgagcttgccgaagcg<br>tcgggcggcttcccacagaagcgccaacgtctcttcgtcgccttcggcgtgcctgatcttgttgaagatgcgttccgcaacgaacttgtcga<br>gtgccgccatgctgacgttgcacgtgccttcgtgctgcccaggtgcggacgggtcgaccaccttccggcgacggcagcggtaagagtc<br>cttgatcgattcttccccgcgcttcgaagtcatgacggcgccacactcgcagtacagcttgtccatggcggacagaatggcttgcccccgg<br>gaaagccccttgccgcgccccctgccgtccaaccacgcctgaagctcataccactcagcgggctcgatgatcggtccgcaatcaagctc<br>gaccggccggagcgtgatcgggtcgcgctgaatgcggtaaccctcaatcttcgtggtcggcgtgccgtccggcttcttcttgtagatcacc<br>tcagcggcgaagcccgcaatacgcgggtcccgaaggattcgcataacggttgccgggtcccaggcgcttgaagcggtcttcttcccaat<br>cgtctcgccccgggtcggcacggcgtcagcgtccatgcgcttacaaagccccgtgatgctgcccgggtgaatggcggcttgactgccc<br>ggcttgaagggaaggtgtttgtgcgtcttgatctcacgccaccaccaccggattacgtcgggctcgaactcgaagggtccggtaagggg<br>agtggtcgagtgcgcaagcttgttgatgacgacattgaccattcggccgttgcgcgtgatctccttcgtctccgaaacaagctcgaagccgt<br>aaggcgccttcccgccgacgtacccgcccaattcgcgctgaaggttcttcgtgtcgagaatcttcgccgacttcagcgaagattctttgtgc<br>gacgcgtcgagccgcataatcaggtgaatcaggtccatgacgtttccctgccggaagacgccttcctgagtggaaacaatcgtcacgccc<br>agggcgagcaattccgagacaatcggaatcgcgtccatgaccttcaggcgcgagaagcgcgacacgtcatagacaatgatcatgttgag<br>ccgcccggcgcggcattcgttcaggatgcgttcgaactccgggcgctccgccgtcccgaacgccgacgtgcccggcgcttcgctgaaa<br>tgcccgacgaacctgaaccggcccccgtcgcgctcgacttcgcgctgaaggtcggccgccttgtcttcgttggcgctacgctgtgtcgct<br>gggcttgctgcgctcgaactctcgcgctcgcgcgactgacggtcgtaagcacccgcgtacgtgtccaccccggtcacaaccccttgtgtc<br>atgtcggcgaccctacgcccccaactgagagaactcaaaggttaccccagttggggcactactcccgaaaaccgcttctgacctgggaa<br>aacgtgaagccccggggcatccgctgagggttgccgccggggcttcggtgtgtccgtcagtacgggccatagaggggcgtcgtgtaac<br>ccacccagggcgattgctccggcacggggaagccgcgccacgccttcgggacgtctggaatcgctagagcttgcatgc |
| pTHS-XGk | agtcagtatccagtcgtgtaggatcccgcaaaagcggcctttgactccctgcaagcctcagcgaccgaatatatcggttatgcgtgggcga<br>tggttgttgtcattgtcggcgcaactatcggtatcaagctgtttaagaaattcacctcgaaagcaagctgataaaccgatacaattaaaggct<br>ccttttggagcctttttttttggagattttcaacgtgaaaaaattattattcgcaattcctttagttgttcctttctattctcactccgctgaaactgttg<br>aaagttgtttagcaaaacctcatacagaaaattcatattaccctgttatccctactccgagacagtcagagggtagaattcggcttgttcacatt<br>cgaaccgtctctgctttgacaacatgctgtgcggtgttgtaaagtcgtggccaggagaatacgacagtctaagtaaggagtgtccatatga<br>gaccgcgcaacgcaattaatgtgagttagctcactcattaggcaccccaggctttacactttatgcttccggctcgtatgttgtgtggaattgt<br>gagcggataacaatttcacacaggaaacagctatgaccatgattacggattcactggccgtcgttttacaacgtcgtgactgggaaaaccc<br>tggcgttacccaacttaatcgccttgcagcacatccccctttcgccagctggcgtaatagcgaagaggcccgcaccgatcgcccttcccaa<br>cagttgcgcagcctgaatggcgaatggtaaggtctccggtgctgcagcactcgttgctttatcggtatattaccctgttatccctaaggacca<br>aaacgaaaaaaggccccccctttcgggaggcctcttttctggaatttggtaccgagccataaactgccaggcatcagacgctcagtggaac<br>gaaaactcacgttaagggtaaaactgtattataagtaaatgcatgtatactaaactcacaaattagagcttcaatttaattatatcagttattacc<br>cataacttcgtatagcatacattatacgaagttatcccgggtaccgagctcgattcgtaacttacacgcgcctcgtatcttttaatgatggaata<br>atttgggaatttactctgtgtttatttatttttatgttttgtatttggattttagaaagtaaataaagaaggtagaagagttacggaatgaagaaaaa<br>aaaataaacaaaggtttaaaaaatttcaacaaaaagcgtactttacatatatatttattagacaagaaaagcagattaaatagatatacattcga<br>ttaacgataagtaaaatgtaaaatcacaggattttcgtgtgtggtcttctacacagacaagatgaaacaattcggcattaatacctgagagca<br>ggaagagcaagataaaaggtagtatttgttggcgatccccctagagtcttttacatcttcggaaaacaaaaactatttttttctttaatttcttttttt<br>actttctatttttaatttatatatttatattaaaaaatttaaattataattatttttatagcacgtgagctagagctgtcagaccaagtttacgagctcg<br>cttggactcctgttgatagatccagtaatgacctcagaactccatctggatttgttcagaacgctcggttgccgccgggcgttttttattggtga<br>gaatccaagcactagggacagtaagacgggtaagcctgttgatgataccgctgccttactgggtgcattagccagtctgaatgacctgtca<br>cgggataatccgaagtggtcagactggaaaatcagagggcaggaactgctgaacagcaaaaagtcagatagcaccacatagcagacc<br>cgccataaaacgccctgagaagcccgtgacgggcttttcttgtattatgggtagtttccttgcatgaatccataaaaggcgcctgtagtgcca<br>tttacccccattcactgccagagccgtgagcgcagcgaactgaatgtcacgaaaaagacagcgactcaggtgcctgatggtcggagaca<br>aaaggaatattcagcgatttgcccgagcttgcgagggtgctacttaagcctttagggttttaaggtctgttttgtagaggagcaaacagcgttt<br>gcgacatccttttgtaatactgcggaactgactaaagtagtgagttatacacagggctgggatctattctttttatctttttttattctttctttattcta |

| | |
|---|---|
| | taaattataaccacttgaatataaacaaaaaaaacacacaaaggtctagcggaatttacagagggtctagcagaatttacaagttttccagca<br>aaggtctagcagaatttacagatacccacaactcaaaggaaaaggacatgtaattatcattgactagcccatctcaattggtatagtgattaa<br>aatcacctagaccaattgagatgtatgtctgaattagttgttttcaaagcaaatgaactagcgattagtcgctatgacttaacggagcatgaaa<br>ccaagctaattttatgctgtgtggcactactcaaccccacgattgaaaaccctacaaggaaagaacggacggtatcgttcacttataaccaa<br>tacgctcagatgatgaacatcagtagggaaaatgcttatggtgtattagctaaagcaaccagagagctgatgacgagaactgtggaaatca<br>ggaatcctttggttaaaggctttgagattttccagtggacaaactatgccaagttctcaagcgaaaaattagaattagtttttagtgaagagata<br>ttgccttatcttttccagttaaaaaaattcataaaatataatctggaacatgttaagtcttttgaaaacaaatactctatgaggatttatgagtggtt<br>attaaaagaactaacacaaaagaaaactcacaaggcaaatatagagattagccttgatgaatttaagttcatgttaatgcttgaaaataactac<br>catgagtttaaaaggcttaaccaatgggttttgaaaccaataagtaaagatttaaacacttacagcaatatgaaattggtggttgataagcga<br>ggccgcccgactgatacgttgattttccaagttgaactagatagacaaatggatctcgtaaccgaacttgagaacaaccagataaaaatga<br>atggtgacaaaataccaacaaccattacatcagattcctacctacgtaacggactaagaaaaacactacacgatgctttaactgcaaaaatt<br>cagctcaccagttttgaggcaaaatttttgagtgacatgcaaagtaagcatgatctcaatggttcgttctcatggctcacgcaaaaacaacga<br>accacactagagaacatactggctaaatacggaaggatctgaggttcttatggctcttgtatctatcagtgaagcatcaagactaacaaaca<br>aaagtagaacaactgttcaccgttagatatcaaagggaaaactgtccatatgcacagatgaaaacggtgtaaaaaagatagatacatcaga<br>gcttttacgagtttttggtgcatttaaagctgttcaccatgaacagatcgacaatgtaactactagagggttcatgtgcagctccatcagcaaa<br>aggggatgataagtttatcaccaccgactatttgcaacagtgccgttgatcgtgctatgatcgactggatcggcggggcctgggacggcg<br>cctcgccgtcgcagaaccaggcggtggcgtacaccgtcgcctcggtcggcccgtagagattggcgatcccgaccgcagcaccaccga<br>gaacgtccccgacgtggccgaccagcccgtcatcgtcaacgcctgatccgcggtgcggacaggccgtgtcgcgaccggccgtgcgga<br>attaagccggcccgtaccctgtgaatagaggtccgctgtgacacaagaatccctgttacttctcgaccgtattgattcggatgattcctacgc<br>gagcctgcggaacgaccaggagttctgggagccgctggcccgccgagccctggaggagctcgggctgccggtgccgccggtgctgc<br>gggtgcccggcgagagcaccaaccccgtactggtcggcgagcccggcccggtgatcaagctgttcggcgagcactggtgcggtccg<br>gagagcctcgcgtcggagtcggaggcgtacgcggtcctggcggacgccccggtgccggtgccccgcctcctcggccgcggcgagct<br>gcggcccggcaccggagcctggccgtggccctacctggtgatgagccggatgaccggcaccacctggcggtccgcgatggacggca<br>cgaccgaccggaacgcgctgctcgccctggcccgcgaactcggccgggtgctcggccggctgcacagggtgccgctgaccgggaac<br>accgtgctcacccccccattccgaggtcttcccggaactgctgcgggaacgccgcgcggcgaccgtcgaggaccaccgcgggtggggc<br>tacctctcgccccggctgctggaccgcctggaggactggctgccggacgtggacacgctgctggccggccgcgaaccccggttcgtcc<br>acggcgacctgcacgggaccaacatcttcgtggacctggccgcgaccgaggtcaccgggatcgtcgacttcaccgacgtctatgcggg<br>agactcccgctacagcctggtgcaactgcatctcaacgccttccggggcgaccgcgagatcctggccgcgctgctcgacggggcgcag<br>tggaagcggaccgaggacttcgcccgcgaactgctcgccttcaccttcctgcacgacttcgaggtgttcgaggagacgccgctggatctc<br>tccggcttcaccgatccggaggaactggcgcagttcctctgggggccgccggacaccgcccccggcgcctgacgccccgggccgcc<br>cggcgccgcccccggcccccggcggccgccgggcggagccccgcccggcggcccgggagccccgggcccgcgccgaagcccg<br>ctgctgcgagctagctagagtcgacctgggtccccggggatcggtcttgccttgctcgtcggtgatgtacttcaccagctccgcgaagtcg<br>ctcttcttgatggagcgcatggggacgtgcttggcaatcacgcgcacccccggccgttttagcggctaaaaaagtcatggctctgccctc<br>gggcggaccacgcccatcatgaccttgccaagctcgtcctgcttctcttcgatcttcgccagcagggcgaggatcgtggcatcaccgaac<br>cgcgccgtgcgcgggtcgtcggtgagccagagtttcagcaggccgcccaggcggcccaggtcgccattgatgcgggccagctcgcg<br>gacgtgctcatagtccacgacgcccgtgattttgtagccctggccgacggccagcaggtaggccgacaggctcatgccggccgccgcc<br>gccttttcctcaatcgctcttcgttcgtctggaaggcagtacaccttgataggtgggctgcccttcctggttggcttggtttcatcagccatccg<br>cttgccctcatctgttacgccggcggtagccggccagcctcgcagagcaggattcccgttgagcaccgccaggtgcgaataagggaca<br>gtgaagaaggaacacccgctcgcgggtgggcctacttcacctatcctgcccggctgacgccgttggatacaccaaggaaagtctacacg<br>aacccttttggcaaaatcctgtatatcgtgcgaaaaaggatggatataccgaaaaaatcgctataatgaccccgaagcagggttatgcagcg<br>gaaaagatccgtcgacctgtcacgccgccgctgtgaacccgttcagcgtctccggctccgtctcttcctcgaccttcggcttcgcccagtg<br>gatacgcacacggtcggcgatcgggggaacccgtccgggccgctgaacgcccttcggaagcttgatcacctcgatccggtccacgaag<br>agccggacgaacgcccgacggtcttcaaggggcgctagcgcccaccacgaacccggtcccgtcgggtcgctgccccggtcgcccag<br>ccattcgccgatcggcagcacgggggagtcggcggcgtcgagctgacgaagccgttcttcggccccttccatgcggagcgtcagcgc<br>ggcttcctcttcgagaaacgcgcgccgtcccatggcggaccggtagccgccgttccgcttgtcttcgtaaagctcttcgagcgccttcacg<br>gcgtccgcgcgctgtgccatcagttcggagcgctgacccttcaactccgggcgttcgtgcgtcgctccccagcgccgtgcggcttcgtac<br>atgagcgctgccgtgtcgccttcgaggtcgtcagggtcggcagggtcgaaggccgtgatgcgcgcgaagatcgcgccgacgatgtac<br>gggtcaaggttgtgcatcgtgatcgagcatgacgacccgtcatgaacccggcgggggcacttgcacgcgtacgacgacttgtgagcttg<br>atcaccttcgcggacgttgcccgccatggtcgacccgttgctgtaacccgtctccgggtcgagctggccggagccgtagcagtaaagga<br>cgtccattgccgacaggagcgattgcccccggtactgacccttcccgcgtcctcgaccctgaagccactcctgaagttcccaccattccgc<br>cgggggaatgtacggctcgaagccgggcagggtcagcggctccatggtgaccgggtcgcgcctgatcttgtaatggctgaagccgccc<br>cgcgaaccgtcggcgcgcaccttgtatgcgatgtcagcttgatacccggcaatgcgcgggtcgctgagtacgcgcttcaaaacgccggg<br>gtcccaatcggaaccggcgcgcttcttaccgaccagcgtgccgcgcgtaggcaccttgtcgcggtacagccgctcacaaagcccgttca<br>gcgacccagggtgaaacgacccggctccgccacccttgaagggcgtatcgcggtgcgtcttgatctcctgccacgcccagcggattacc<br>gccccttcgctgccgtgtgcgccttcccaggtgtgcgcgctgggcaccagtcggcgaatggcaaccagctttccgccgtcttccgggttc<br>ggaaccatttcctcgaccgtgtcgaatccgtacggcgtcgaccccgtgtgtccgcccagccgcttcgccaattccttagcgttcgacacgg<br>cgacgctcttgttcttcgactcatcatgcgaagcctgaaggcgcatgatcaggtgaataaggtccatcatttcgccggggcggaatgtgcct<br>tcgttcacgctgacaatggtcacgcccagccggagcaattccgtgacgaccggaataatgtccagcggctcttcgcggctgaggcgcga<br>aatgtaatgaacaatgatcatgttcatttcccggttccggcacatgtccaaaatccggttgaactccggccggtcgacgcccgtgaatgccg<br>acgtgccgggcgcttcgctgaagtgacccagccacttcacctcgacgccgtcgcgcgcgtactccttcgccagcgcctcagccttcccc<br>cggttcgcggcgcgctgggtggccggtgaagcggtcgaactgttctcccgttccgccgactgtcggtcgtagccgcctgccagaatgac<br>catgccttcaaccttcacgctcatgccgggcaggctatcccgcccaactgggtaagagcaaagactaacactgttgggctggaactactc<br>ccgaaaaccgcttctgacctgggaaaacgtgaagccccggggcatccgctgagggttgccgccggggcttcggtgtgtccgtcagtac<br>gggccatagaggggcgtcgtgtaacccacccagggcgattgctccggcacggggaagccgcgccacgccttcgggacgtctggaatc<br>gctagagcttgcatgc |

| | |
|---|---|
| pTHS-XG23 | agtcagtatccagtcgtgtaggatcccgcaaaagcggcctttgactccctgcaagcctcagcgaccgaatatatcggttatgcgtgggcga<br>tggttgttgtcattgtcggcgcaactatcggtatcaagctgtttaagaaattcacctcgaaagcaagctgataaaccgatacaattaaaggct<br>cctttggagcctttttttttggagattttcaacgtgaaaaaattattattcgcaattcctttagttgttcctttctattctcactccgctgaaactgttg<br>aaagttgtttagcaaaacctcatacagaaaattcatattaccctgttatccctactccgagacagtcagagggtagaattcggcttgttcacatt<br>cgaaccgtctctgctttgacaacatgctgtgcggtgttgtaaagtcctgctaaggagaatacgacagtctaagtaaggagtgtccatatgag<br>accgcgcaacgcaattaatgtgagttagctcactcattaggcaccccaggctttacactttatgcttccggctcgtatgttgtgtggaattgtg<br>agcggataacaatttcacacaggaaacagctatgaccatgattacggattcactggccgtcgttttacaacgtcgtgactgggaaaaccct<br>ggcgttacccaacttaatcgccttgcagcacatccccctttcgccagctggcgtaatagcgaagaggcccgcaccgatcgcccttcccaa<br>cagttgcgcagcctgaatggcgaatggtaaggtctccggtgctgcagcactcgttgctttatcggtatattaccctgttatccctaaggacca<br>aaacgaaaaaaggccccccctttcgggaggcctcttttctggaatttggtaccgagccataaactgccaggcatcagacgctcagtggaac<br>gaaaactcacgttaagggtaaaactgtattataagtaaatgcatgtatactaaactcacaaattagagcttcaatttaattatatcagttattacc<br>cataacttcgtatagcatacattatacgaagttatcccgggtaccgagctcgattcgtaacttacacgcgcctcgtatcttttaatgatggaata<br>atttgggaatttactctgtgtttatttatttttatgttttgtatttggattttagaaagtaaataaagaaggtagaagagttacggaatgaagaaaaa<br>aaaataaacaaaggtttaaaaaatttcaacaaaaagcgtactttacatatatatttattagacaagaaaagcagattaaatagatatacattcga<br>ttaacgataagtaaaatgtaaaatcacaggattttcgtgtgtggtcttctacacagacaagatgaaacaattcggcattaatacctgagagca<br>ggaagagcaagataaaaggtagtatttgttggcgatccccctagagtcttttacatcttcggaaaacaaaaactatttttttctttaatttcttttttt<br>actttctatttttaatttatatatttatattaaaaaatttaaattataattatttttatagcacgtgagctagagctgtcagaccaagtttacgagctcg<br>cttggactcctgttgatagatccagtaatgacctcagaactccatctggatttgttcagaacgctcggttgccgccgggcgtttttattggtga<br>gaatccaagcactagggacagtaagacgggtaagcctgttgatgataccgctgccttactgggtgcattagccagtctgaatgacctgtca<br>cgggataatccgaagtggtcagactggaaaatcagagggcaggaactgctgaacagcaaaaagtcagatagcaccacatagcagacc<br>cgccataaaacgccctgagaagcccgtgacgggcttttcttgtattatgggtagtttccttgcatgaatccataaaaggcgcctgtagtgcca<br>tttacccccattcactgccagagccgtgagcgcagcgaactgaatgtcacgaaaaagacagcgactcaggtgcctgatggtcggagaca<br>aaaggaatattcagcgatttgcccgagcttgcgagggtgctacttaagcctttagggttttaaggtctgttttgtagaggagcaaacagcgttt<br>gcgacatccttttgtaatactgcggaactgactaaagtagtgagttatacacagggctgggatctattctttttatctttttttattctttctttattcta<br>taaattataaccacttgaatataaacaaaaaaaacacacaaaggtctagcggaatttacagagggtctagcagaatttacaagttttccagca<br>aaggtctagcagaatttacagatacccacaactcaaaggaaaaggacatgtaattatcattgactagcccatctcaattggtatagtgattaa<br>aatcacctagaccaattgagatgtatgtctgaattagttgttttcaaagcaaatgaactagcgattagtcgctatgacttaacggagcatgaaa<br>ccaagctaattttatgctgtgtggcactactcaaccccacgattgaaaaccctacaaggaaagaacggacggtatcgttcacttataaccaa<br>tacgctcagatgatgaacatcagtagggaaaatgcttatggtgtattagctaaagcaaccagagagctgatgacgagaactgtggaaatca<br>ggaatcctttggttaaaggctttgagattttccagtggacaaactatgccaagttctcaagcgaaaaattagaattagtttttagtgaagagata<br>ttgccttatcttttccagttaaaaaaattcataaaatataatctggaacatgttaagtcttttgaaaacaaatactctatgaggatttatgagtggtt<br>attaaaagaactaacacaaaagaaaactcacaaggcaaatatagagattagccttgatgaatttaagttcatgttaatgcttgaaaataactac<br>catgagtttaaaaggcttaaccaatgggttttgaaaccaataagtaaagatttaaacacttacagcaatatgaaattggtggttgataagcga<br>ggccgcccgactgatacgttgattttccaagttgaactagatagacaaatggatctcgtaaccgaacttgagaacaaccagataaaaatga<br>atggtgacaaaataccaacaaccattacatcagattcctacctacgtaacggactaagaaaaacactacacgatgctttaactgcaaaaatt<br>cagctcaccagttttgaggcaaaatttttgagtgacatgcaaagtaagcatgatctcaatggttcgttctcatggctcacgcaaaaacaacga<br>accacactagagaacatactggctaaatacggaaggatctgaggttcttatggctcttgtatctatcagtgaagcatcaagactaacaaaca<br>aaagtagaacaactgttcaccgttagatatcaaagggaaaactgtccatatgcacagatgaaaacggtgtaaaaaagatagatacatcaga<br>gcttttacgagtttttggtgcatttaaagctgttcaccatgaacagatcgacaatgtaactactagagggttcatgtgcagctccatcagcaaa<br>aggggatgataagtttatcaccaccgactatttgcaacagtgccgttgatcgtgctatgatcgactggatcggcggggcctgggacggcg<br>cctcgccgtcgcagaaccaggcggtggcgtacaccgtcgcctcggtcggcccgtagagattggcgatcccgaccgcagcaccaccga<br>gaacgtccccgacgtggccgaccagcccgtcatcgtcaacgcctgatccgcggtgcggacaggccgtgtcgcgaccggccgtgcgga<br>attaagccggcccgtaccctgtgaatagaggtccgctgtgacacaagaatccctgttacttctcgaccgtattgattcggatgattcctacgc<br>gagcctgcggaacgaccaggagttctgggagccgctggcccgccgagccctggaggagctcgggctgccggtgccgccggtgctgc<br>gggtgcccggcgagagcaccaaccccgtactggtcggcgagcccggcccggtgatcaagctgttcggcgagcactggtgcggtccg<br>gagagcctcgcgtcggagtcggaggcgtacgcggtcctggcggacgccccggtgccggtgccccgcctcctcggccgcggcgagct<br>gcggcccggcaccggagcctggccgtggccctacctggtgatgagccggatgaccggcaccacctggcggtccgcgatggacggca<br>cgaccgaccggaacgcgctgctcgccctggcccgcgaactcggccgggtgctcggccggctgcacagggtgccgctgaccgggaac<br>accgtgctcaccccccattccgaggtcttcccggaactgctgcgggaacgccgcgcggcgaccgtcgaggaccaccgcgggtggggc<br>tacctctcgccccggctgctggaccgcctggaggactggctgccggacgtggacacgctgctggccggccgcgaaccccggttcgtcc<br>acggcgacctgcacgggaccaacatcttcgtggacctggccgcgaccgaggtcaccgggatcgtcgacttcaccgacgtctatgcggg<br>agactcccgctacagcctggtgcaactgcatctcaacgccttccggggcgaccgcgagatcctggccgcgctgctcgacggggcgcag<br>tggaagcggaccgaggacttcgcccgcgaactgctcgccttcaccttcctgcacgacttcgaggtgttcgaggagacgccgctggatctc<br>tccggcttcaccgatccggaggaactggcgcagttcctctgggggccgccggacaccgcccccggcgcctgacgccccgggccgcc<br>cggcgccgcccccggcccccggcggccgccgggcggagccccgcccggcggcccgggagccccgggcccgcgccgaagcccg<br>ctgctgcgagctagctagagtcgacctgggtccccggggatcggtcttgccttgctcgtcggtgatgtacttcaccagctccgcgaagtcg<br>ctcttcttgatggagcgcatggggacgtgcttggcaatcacgcgcacccccggccgttttagcggctaaaaaagtcatggctctgccctc<br>gggcggaccacgcccatcatgaccttgccaagctcgtcctgcttctcttcgatcttcgccagcagggcgaggatcgtggcatcaccgaac<br>cgcgccgtgcgcgggtcgtcggtgagccagagtttcagcaggccgcccaggcggcccaggtcgccattgatgcgggccagctcgcg<br>gacgtgctcatagtccacgacgcccgtgattttgtagccctggccgacggccagcaggtaggccgacaggctcatgccggccgccgcc<br>gccttttcctcaatcgctcttcgttcgtctggaaggcagtacaccttgataggtgggctgcccttcctggttggcttggtttcatcagccatccg<br>cttgccctcatctgttacgccggcggtagccggccagcctcgcagagcaggattcccgttgagcaccgccaggtgcgaataagggaca<br>gtgaagaaggaacacccgctcgcgggtgggcctacttcacctatcctgcccggctgacgccgttggatacaccaaggaaagtctacacg<br>aaccctttggcaaaatcctgtatatcgtgcgaaaaaggatggatataccgaaaaaatcgctataatgaccccgaagcagggttatgcagcg<br>gaaaagatccgtcgacctgtcacgccgccgctgtgaacccgttcagcgtctccggctccgtctcttcctcgaccttcggcttcgcccagtg<br>gatacgcacacggtcggcgatcgggggaacccgtccgggccgctgaacgcccttcggaagcttgatcacctcgatccggtccacgaag<br>agccggacgaacgcccgacggtcttcaaggggcgctagcgcccaccacgaacccggtcccgtcgggtcgctgccccggtcgcccag |

| | |
|---|---|
| | ccattcgccgatcggcagcacgggggagtcggcggcgtcgagctgacgaagccgttcttcggccccttccatgcggagcgtcagcgc<br>ggcttcctcttcgagaaacgcgcgccgtcccatggcggaccggtagccgccgttccgcttgtcttcgtaaagctcttcgagcgccttcacg<br>gcgtccgcgcgctgtgccatcagttcggagcgctgacccttcaactccgggcgttcgtgcgtcgctccccagcgccgtgcggcttcgtac<br>atgagcgctgccgtgtcgccttcgaggtcgtcagggtcggcagggtcgaaggccgtgatgcgcgcgaagatcgcgccgacgatgtac<br>gggtcaaggttgtgcatcgtgatcgagcatgacgacccgtcatgaacccggcgggggcacttgcacgcgtacgacgacttgtgagcttg<br>atcaccttcgcggacgttgcccgccatggtcgacccgttgctgtaacccgtctccgggtcgagctggccggagccgtagcagtaaagga<br>cgtccattgccgacaggagcgattgcccccggtactgacccttcccgcgtcctcgaccctgaagccactcctgaagttcccaccattccgc<br>cgggggaatgtacggctcgaagccgggcagggtcagcggctccatggtgaccgggtcgcgcctgatcttgtaatggctgaagccgccc<br>cgcgaaccgtcggcgcgcaccttgtatgcgatgtcagcttgatacccggcaatgcgcgggtcgctgagtacgcgcttcaaaacgccggg<br>gtcccaatcggaaccggcgcgcttcttaccgaccagcgtgccgcgcgtaggcaccttgtcgcggtacagccgctcacaaagcccgttca<br>gcgacccagggtgaaacgacccggctccgccacccttgaagggcgtatcgcggtgcgtcttgatctcctgccacgcccagcggattacc<br>gccccttcgctgccgtgtgcgccttcccaggtgtgcgcgctgggcaccagtcggcgaatggcaaccagctttccgccgtcttccgggttc<br>ggaaccatttcctcgaccgtgtcgaatccgtacggcgtcgaccccgtgtgtccgcccagccgcttcgccaattccttagcgttcgacacgg<br>cgacgctcttgttcttcgactcatcatgcgaagcctgaaggcgcatgatcaggtgaataaggtccatcatttcgccggggcggaatgtgcct<br>tcgttcacgctgacaatggtcacgcccagccggagcaattccgtgacgaccggaataatgtccagcggctcttcgcggctgaggcgcga<br>aatgtaatgaacaatgatcatgttcatttcccggttccggcacatgtccaaaatccggttgaactccggccggtcgacgcccgtgaatgccg<br>acgtgccgggcgcttcgctgaagtgacccagccacttcacctcgacgccgtcgcgcgcgtactccttcgccagcgcctcagccttcccc<br>cggttcgcggcgcgctgggtggccggtgaagcggtcgaactgttctcccgttccgccgactgtcggtcgtagccgcctgccagaatgac<br>catgccttcaaccttcacgctcatgccgggcaggctatcccgcccaactgggtaagagcaaagactaacactgttgggctggaactactc<br>ccgaaaaccgcttctgacctgggaaaacgtgaagccccggggcatccgctgagggttgccgccggggcttcggtgtgtccgtcagtac<br>gggccatagaggggcgtcgtgtaacccacccagggcgattgctccggcacggggaagccgcgccacgccttcgggacgtctggaatc<br>gctagagcttgcatgc |
| pTHS-XGe | agtcagtatccagtcgtgtaggatcccgcaaaagcggcctttgactccctgcaagcctcagcgaccgaatatatcggttatgcgtgggcga<br>tggttgttgtcattgtcggcgcaactatcggtatcaagctgtttaagaaattcacctcgaaagcaagctgataaaccgatacaattaaaggct<br>cctttggagcctttttttttggagattttcaacgtgaaaaaattattattcgcaattcctttagttgttcctttctattctcactccgctgaaactgttg<br>aaagttgtttagcaaaacctcatacagaaaattcatattaccctgttatccctactccgagacagtcagagggtagaattcgtaccagcccga<br>cccgagcacgcgccggcacgcctggtcgatgtcggaccggagttcgaggtacgcggcttgcaggtccaggaaggggacgtccatgc<br>gagtgtccgttcgagtggcggcttgcgcccgatgctagtcgcggttgatcggcgatcgcaggtgcacgcggtcgatcttgacggctggc<br>gagaggtgcggggaggatctgaccgacgcggtccacacgtggcaccgcgatgctgttgtgggcacaatcgtgccggttggtaggatcg<br>atccactagttctagaaataattttgtttaacattaaagaggagaaattacatatgagaccgcgcaacgcaattaatgtgagttagctcactcat<br>taggcaccccaggctttacactttatgcttccggctcgtatgttgtgtggaattgtgagcggataacaatttcacacaggaaacagctatgac<br>catgattacggattcactggccgtcgttttacaacgtcgtgactgggaaaaccctggcgttacccaacttaatcgccttgcagcacatcccc<br>ctttcgccagctggcgtaatagcgaagaggcccgcaccgatcgcccttcccaacagttgcgcagcctgaatggcgaatggtaaggtctc<br>cggtgctgcagcactcgttgctttatcggtatattaccctgttatccctaaggaccaaaacgaaaaaaggccccccctttcgggaggcctcttt<br>tctggaatttggtaccgagccataaactgccaggcatcagacgctcagtggaacgaaaactcacgttaagggtaaaactgtattataagta<br>aatgcatgtatactaaactcacaaattagagcttcaatttaattatatcagttattacccataacttcgtatagcatacattatacgaagttatccc<br>gggtaccgagctcgattcgtaacttacacgcgcctcgtatcttttaatgatggaataatttgggaatttactctgtgtttatttatttttatgttttgta<br>tttggattttagaaagtaaataaagaaggtagaagagttacggaatgaagaaaaaaaaaataaacaaaggtttaaaaaatttcaacaaaaagc<br>gtactttacatatatatttattagacaagaaaagcagattaaatagatatacattcgattaacgataagtaaaatgtaaaatcacaggattttcgt<br>gtgtggtcttctacacagacaagatgaaacaattcggcattaatacctgagagcaggaagagcaagataaaaggtagtatttgttggcgat<br>ccccctagagtcttttacatcttcggaaaacaaaaactatttttctttaatttctttttttactttctatttttaatttatatatttatattaaaaaatttaaat<br>tataattatttttatagcacgtgagctagagctgtcagaccaagtttacgagctcgcttggactcctgttgatagatccagtaatgacctcagaa<br>ctccatctggatttgttcagaacgctcggttgccgccgggcgttttttattggtgagaatccaagcactagggacagtaagacgggtaagcc<br>tgttgatgataccgctgccttactgggtgcattagccagtctgaatgacctgtcacgggataatccgaagtggtcagactggaaaatcagag<br>ggcaggaactgctgaacagcaaaaagtcagatagcaccacatagcagacccgccataaaacgccctgagaagcccgtgacgggctttt<br>cttgtattatgggtagtttccttgcatgaatccataaaaggcgcctgtagtgccatttacccccattcactgccagagccgtgagcgcagcga<br>actgaatgtcacgaaaaagacagcgactcaggtgcctgatggtcggagacaaaaggaatattcagcgatttgcccgagcttgcgagggt<br>gctacttaagcctttagggttttaaggtctgttttgtagaggagcaaacagcgtttgcgacatccttttgtaatactgcggaactgactaaagta<br>gtgagttatacacagggctgggatctattctttttatctttttttattctttctttattctataaattataaccacttgaatataaacaaaaaaaacaca<br>caaaggtctagcggaatttacagagggtctagcagaatttacaagttttccagcaaaggtctagcagaatttacagatacccacaactcaaa<br>ggaaaaggacatgtaattatcattgactagcccatctcaattggtatagtgattaaaatcacctagaccaattgagatgtatgtctgaattagtt<br>gttttcaaagcaaatgaactagcgattagtcgctatgacttaacggagcatgaaaccaagctaattttatgctgtgtggcactactcaacccc<br>acgattgaaaaccctacaaggaaagaacggacggtatcgttcacttataaccaatacgctcagatgatgaacatcagtagggaaaatgctt<br>atggtgtattagctaaagcaaccagagagctgatgacgagaactgtggaaatcaggaatcctttggttaaaggctttgagattttccagtgg<br>acaaactatgccaagttctcaagcgaaaaattagaattagtttttagtgaagagatattgccttatcttttccagttaaaaaaattcataaaatata<br>atctggaacatgttaagtcttttgaaaacaaatactctatgaggatttatgagtggttattaaaagaactaacacaaaagaaaactcacaaggc<br>aaatatagagattagccttgatgaatttaagttcatgttaatgcttgaaaataactaccatgagtttaaaaggcttaaccaatgggttttgaaacc<br>aataagtaaagatttaaacacttacagcaatatgaaattggtggttgataagcgaggccgcccgactgatacgttgattttccaagttgaact<br>agatagacaaatggatctcgtaaccgaacttgagaacaaccagataaaaatgaatggtgacaaaataccaacaaccattacatcagattcc<br>tacctacgtaacggactaagaaaaacactacacgatgctttaactgcaaaaattcagctcaccagttttgaggcaaaatttttgagtgacatg<br>caaagtaagcatgatctcaatggttcgttctcatggctcacgcaaaaacaacgaaccacactagagaacatactggctaaatacggaagg<br>atctgaggttcttatggctcttgtatctatcagtgaagcatcaagactaacaaacaaaagtagaacaactgttcaccgttagatatcaaaggg<br>aaaactgtccatatgcacagatgaaaacggtgtaaaaaagatagatacatcagagcttttacgagtttttggtgcatttaaagctgttcaccat<br>gaacagatcgacaatgtaactactagagggttcatgtgcagctccatcagcaaaaggggatgataagtttatcaccaccgactatttgcaac<br>agtgccgttgatcgtgctatgatcgactggatcggcggggcctgggacggcgcctcgccgtcgcagaaccaggcggtggcgtacaccg<br>tcgcctcggtcggcccgtagagattggcgatcccgaccgcagcaccaccgagaacgtccccgacgtggccgaccagcccgtcatcgtc<br>aacgcctgatccgcggtgcggacaggccgtgtcgcgaccggccgtgcggaattaagccggcccgtaccctgtgaatagaggtccgctg |

| | |
|---|---|
| | tgacacaagaatccctgttacttctcgaccgtattgattcggatgattcctacgcgagcctgcggaacgaccaggagttctgggagccgct<br>ggcccgccgagccctggaggagctcgggctgccggtgccgccggtgctgcgggtgcccggcgagagcaccaaccccgtactggtcg<br>gcgagcccggcccggtgatcaagctgttcggcgagcactggtgcggtccggagagcctcgcgtcggagtcggaggcgtacgcggtcc<br>tggcggacgccccggtgccggtgccccgcctcctcggccgcggcgagctgcggcccggcaccggagcctggccgtggccctacctg<br>gtgatgagccggatgaccggcaccacctggcggtccgcgatggacggcacgaccgaccggaacgcgctgctcgccctggcccgcga<br>actcggccgggtgctcggccggctgcacagggtgccgctgaccgggaacaccgtgctcacccccccattccgaggtcttcccggaactg<br>ctgcgggaacgccgcgcggcgaccgtcgaggaccaccgcgggtggggctacctctcgccccggctgctggaccgcctggaggactg<br>gctgccggacgtggacacgctgctggccggccgcgaaccccggttcgtccacggcgacctgcacgggaccaacatcttcgtggacct<br>ggccgcgaccgaggtcaccgggatcgtcgacttcaccgacgtctatgcgggagactcccgctacagcctggtgcaactgcatctcaacg<br>ccttccggggcgaccgcgagatcctggccgcgctgctcgacggggcgcagtggaagcggaccgaggacttcgcccgcgaactgctc<br>gccttcaccttcctgcacgacttcgaggtgttcgaggagacgccgctggatctctccggcttcaccgatccggaggaactggcgcagttc<br>ctctgggggccgccggacaccgcccccggcgcctgacgccccgggccgcccggcgccgcccccggcccccggcggccgccggg<br>cggagccccgcccggcggcccgggagccccgggcccgcgccgaagcccgctgctgcgagctagctagagtcgacctgggtccccg<br>gggatcggtcttgccttgctcgtcggtgatgtacttcaccagctccgcgaagtcgctcttcttgatggagcgcatggggacgtgcttggcaa<br>tcacgcgcacccccccggccgttttagcggctaaaaaagtcatggctctgccctcgggcggaccacgcccatcatgaccttgccaagctc<br>gtcctgcttctcttcgatcttcgccagcagggcgaggatcgtggcatcaccgaaccgcgccgtgcgcgggtcgtcggtgagccagagttt<br>cagcaggccgcccaggcggcccaggtcgccattgatgcgggccagctcgcggacgtgctcatagtccacgacgcccgtgattttgtag<br>ccctggccgacggccagcaggtaggccgacaggctcatgccggccgccgccgccttttcctcaatcgctcttcgttcgtctggaaggca<br>gtacaccttgataggtgggctgcccttcctggttggcttggtttcatcagccatccgcttgccctcatctgttacgccggcggtagccggcca<br>gcctcgcagagcaggattcccgttgagcaccgccaggtgcgaataagggacagtgaagaaggaacacccgctcgcgggtgggcctac<br>ttcacctatcctgcccggctgacgccgttggatacaccaaggaaagtctacacgaaccctttggcaaaatcctgtatatcgtgcgaaaaag<br>gatggatataccgaaaaaatcgctataatgaccccgaagcagggttatgcagcggaaaagatccgtcgacctgtcacgccgccgctgtg<br>aacccgttcagcgtctccggctccgtctcttcctcgaccttcggcttcgcccagtggatacgcacacggtcggcgatcgggggaacccgt<br>ccgggccgctgaacgcccttcggaagcttgatcacctcgatccggtccacgaagagccggacgaacgcccgacggtcttcaaggggc<br>gctagcgcccaccacgaacccggtcccgtcgggtcgctgccccggtcgcccagccattcgccgatcggcagcacgggggagtcggc<br>ggcgtcgagctgacgaagccgttcttcggccccttccatgcggagcgtcagcgcggcttcctcttcgagaaacgcgcgccgtcccatgg<br>cggaccggtagccgccgttccgcttgtcttcgtaaagctcttcgagcgccttcacggcgtccgcgcgctgtgccatcagttcggagcgctg<br>acccttcaactccgggcgttcgtgcgtcgctccccagcgccgtgcggcttcgtacatgagcgctgccgtgtcgccttcgaggtcgtcagg<br>gtcggcagggtcgaaggccgtgatgcgcgcgaagatcgcgccgacgatgtacgggtcaaggttgtgcatcgtgatcgagcatgacgac<br>ccgtcatgaacccggcgggggcacttgcacgcgtacgacgacttgtgagcttgatcaccttcgcggacgttgcccgccatggtcgaccc<br>gttgctgtaacccgtctccgggtcgagctggccggagccgtagcagtaaaggacgtccattgccgacaggagcgattgcccccggtact<br>gacccttcccgcgtcctcgaccctgaagccactcctgaagttcccaccattccgccgggggaatgtacggctcgaagccgggcagggtc<br>agcggctccatggtgaccgggtcgcgcctgatcttgtaatggctgaagccgccccgcgaaccgtcggcgcgcaccttgtatgcgatgtc<br>agcttgatacccggcaatgcgcgggtcgctgagtacgcgcttcaaaacgccggggtcccaatcggaaccggcgcgcttcttaccgacca<br>gcgtgccgcgcgtaggcaccttgtcgcggtacagccgctcacaaagcccgttcagcgacccagggtgaaacgacccggctccgccac<br>ccttgaagggcgtatcgcggtgcgtcttgatctcctgccacgcccagcggattaccgccccttcgctgccgtgtgcgccttcccaggtgtg<br>cgcgctgggcaccagtcggcgaatggcaaccagctttccgccgtcttccgggttcggaaccatttcctcgaccgtgtcgaatccgtacgg<br>cgtcgaccccgtgtgtccgcccagccgcttcgccaattccttagcgttcgacacggcgacgctcttgttcttcgactcatcatgcgaagcct<br>gaaggcgcatgatcaggtgaataaggtccatcatttcgccggggcggaatgtgccttcgttcacgctgacaatggtcacgcccagccgg<br>agcaattccgtgacgaccggaataatgtccagcggctcttcgcggctgaggcgcgaaatgtaatgaacaatgatcatgttcatttcccggtt<br>ccggcacatgtccaaaatccggttgaactccggccggtcgacgcccgtgaatgccgacgtgccgggcgcttcgctgaagtgacccagc<br>cacttcacctcgacgccgtcgcgcgcgtactccttcgccagcgcctcagccttcccccggttcgcggcgcgctgggtggccggtgaagc<br>ggtcgaactgttctcccgttccgccgactgtcggtcgtagccgcctgccagaatgaccatgccttcaaccttcacgctcatgccgggcagg<br>ctatcccgcccaactgggtaagagcaaagactaacactgttgggctggaactactcccgaaaaccgcttctgacctgggaaaacgtgaa<br>gccccggggcatccgctgagggttgccgccggggcttcggtgtgtccgtcagtacgggccatagaggggcgtcgtgtaacccacccag<br>ggcgattgctccggcacggggaagccgcgccacgccttcgggacgtctggaatcgctagagcttgcatgc |
| pTHS-<br>XGSN* | agtcagtatccagtcgtgtaggatcccgcaaaagcggcctttgactccctgcaagcctcagcgaccgaatatatcggttatgcgtgggcga<br>tggttgttgtcattgtcggcgcaactatcggtatcaagctgtttaagaaattcacctcgaaagcaagctgataaaccgatacaattaaaggct<br>ccttttggagccttttttttggagattttcaacgtgaaaaaattattattcgcaattcctttagttgttcctttctattctcactccgctgaaactgttg<br>aaagttgtttagcaaaacctcatacagaaaattcatattaccctgttatccctactccgagacagtcagagggtagaattctcatacgaattta<br>ccctcgcttccacgacaacaccgataatcttgcagtttccgttgataggagtcataggccatgaaggattcaggcctttcaggtacttctgac<br>cgccatctatgaccagtttcttgaatgttgcttcgttcgcgtcagtcagtttggctacaacaaggcttccattcactggctcccgtccagtatct<br>acgagcaccatatgaccttcagggatgctttgacctacaggtgaggtcatggaatcaccttcaaccttcagccagaatccattgccgagga<br>ggttaacgtcactgtcataccattcatcaatgtccttgatatcgtagggttcacaagcttcacaccacgaaccagctctaaccatgctaatcaa<br>tggatatttccctttgggctcaacgtgcccaacaaatctgctgccaccgccaccgctgccaccgccaccccgcttgaacttggcgtaccgc<br>tcccgggcgatctccagggtgctgttccgcacgcgctcgaaccgctccttgtccttctgccacagcgaccgcacggcgaggccgcgga<br>cggagttgaagatcagccagaggatgtcctcggcgtcgtcgcgggacaggccccggctcaccaggacgccgagccacatgtcctcca<br>cgacgaagcggttccgctccacggtccgctggatgccctcgcgcagggcggggtcccggtccgcggcgacgatcaggtcgagcgag<br>atggagaagtcgtcgtccaggaagaactccgcggcgtcgtcgagcatctgctggatgacgtcgtcctccggcttcagcttcgcgaggcg<br>ggcccgggagcgctcggtgatctgctcgtacagccactcgaaggtcgccaggagcagctcgagcttggtggggaagtggtggctctgg<br>gcgccgcgcgacacgcccgcggcgcccgggacgtccgcgatccggaagccggcgtagcccttctcgcgcaggacgccgagggcc<br>gccgcgatcagcttgccctgggtttccatcgcccgctccgcctgggtccgccgcttgggggacatgatgaccaccatctagtatttctcctc<br>tttctctagtattaaacaaaattatttgtagaggctgtttcgtcctcacggactcatcagaccggaaagcacatccggtgacagcttggccac<br>gactttacaggaggtaatccgcgggtgtcaaagcagagacggttcgaatgtgaacacaggatgtgacgagcggtgtggaggcatttacc<br>aatgcttaatcagtgaggcacctatctcagcgatctgtctatttcgttcatccatagttgcctgactccccgtcgtgtagataactacgatacgg<br>gagggcttaccatctggccccagtgctgcaatgataccgcgagatccacgctcaccggctccagatttatcagcaataaaccagccagcc<br>ggaagggccgagcgcagaagtggtcctgcaactttatccgcctccatccagtctattaattgttgccgggaagctagagtaagtagttcgc |

| | |
|---|---|
| | cagttaatagtttgcgcaacgttgttgccattgctacaggcatcgtggtgtcacgctcgtcgtttggtatggcttcattcagctccggttcccaa<br>cgatcaaggcgagttacatgatcccccatgttgtgcaaaaaagcggttagctccttcggtcctccgatcgttgtcagaagtaagttggccgc<br>agtgttatcactcatggttatggcagcactgcataattctcttactgtcatgccatccgtaagatgcttttctgtgactggtgagtactcaaccaa<br>gtcattctgagaatagtgtatgcggcgaccgagttgctcttgcccggcgtcaatacgggataataccgcgccacatagcagaactttaaaa<br>gtgctcatcattggaaaacgttcttcggggcgaaaactctcaaggatcttaccgctgttgagatccagttcgatgtaacccactcgtgcaccc<br>aactgatcttcagcatcttttactttcaccagcgtttctgggtgagcaaaaacaggaaggcaaaatgccgcaaaaaagggaataagggcga<br>cacggaaatgttgaatactcatactcttcctttttcaatattattgaagcatttatcagggttattgtctcatgagcggatacatatttgaatgtattt<br>agaaaaataaacaaataggggttccgcgcacatttccccgaaaagtgccacctgacgtctaagaaaccattattatcatgacattaacctat<br>aaaaataggcgtatcacgaggccctttcatgcctccacaccgctcgtcacatcctgaacaaacagacaatctggtctgtttacgtctctgcttt<br>gacacggacaagcgcatggtgtaaagtaacaaacagacaatctggtctgtttgtattcatatgagaccgcgcaacgcaattaatgtgagtta<br>gctcactcattaggcaccccaggctttacactttatgcttccggctcgtatgttgtgtggaattgtgagcggataacaatttcacacaggaaac<br>agctatgaccatgattacggattcactggccgtcgttttacaacgtcgtgactgggaaaaccctggcgttacccaacttaatcgccttgcagc<br>acatccccctttcgccagctggcgtaatagcgaagaggcccgcaccgatcgcccttcccaacagttgcgcagcctgaatggcgaatggt<br>aaggtctccggtgctgcagcactcgttgctttatcggtatattaccctgttatccctaaggaccaaaacgaaaaaaggccccccctttcggga<br>ggcctcttttctggaatttggtaccgagccataaactgccaggcatcagacgctcagtggaacgaaaactcacgttaagggtaaaactgtat<br>tataagtaaatgcatgtatactaaactcacaaattagagcttcaatttaattatatcagttattacccataacttcgtatagcatacattatacgaa<br>gttatcccgggtaccgagctcgattcgtaacttacacgcgcctcgtatcttttaatgatggaataatttgggaatttactctgtgtttatttattttta<br>tgttttgtatttggattttagaaagtaaataaagaaggtagaagagttacggaatgaagaaaaaaaaataaacaaaggtttaaaaaatttcaac<br>aaaaagcgtactttacatatatatttattagacaagaaaagcagattaaatagatatacattcgattaacgataagtaaaatgtaaaatcacag<br>gattttcgtgtgtggtcttctacacagacaagatgaaacaattcggcattaatacctgagagcaggaagagcaagataaaaggtagtatttgt<br>tggcgatccccctagagtcttttacatcttcggaaaacaaaaactatttttttctttaatttctttttttactttctatttttaatttatatatttatattaaaa<br>aatttaaattataattatttttatagcacgtgagctagagctgtcagaccaagtttacgagctcgcttggactcctgttgatagatccagtaatga<br>cctcagaactccatctggatttgttcagaacgctcggttgccgccgggcgttttttattggtgagaatccaagcactagggacagtaagacg<br>ggtaagcctgttgatgataccgctgccttactgggtgcattagccagtctgaatgacctgtcacgggataatccgaagtggtcagactgga<br>aaatcagagggcaggaactgctgaacagcaaaaagtcagatagcaccacatagcagacccgccataaaacgccctgagaagcccgtg<br>acgggcttttcttgtattatgggtagtttccttgcatgaatccataaaaggcgcctgtagtgccatttacccccattcactgccagagccgtga<br>gcgcagcgaactgaatgtcacgaaaaagacagcgactcaggtgcctgatggtcggagacaaaaggaatattcagcgatttgcccgagct<br>tgcgagggtgctacttaagcctttagggttttaaggtctgttttgtagaggagcaaacagcgtttgcgacatccttttgtaatactgcggaact<br>gactaaagtagtgagttatacacagggctgggatctattctttttatctttttttattctttctttattctataaattataaccacttgaatataaacaaa<br>aaaaacacacaaaggtctagcggaatttacagagggtctagcagaatttacaagttttccagcaaaggtctagcagaatttacagataccc<br>acaactcaaaggaaaaggacatgtaattatcattgactagcccatctcaattggtatagtgattaaaatcacctagaccaattgagatgtatgt<br>ctgaattagttgttttcaaagcaaatgaactagcgattagtcgctatgacttaacggagcatgaaaccaagctaattttatgctgtgtggcacta<br>ctcaaccccacgattgaaaaccctacaaggaaagaacggacggtatcgttcacttataaccaatacgctcagatgatgaacatcagtagg<br>gaaaatgcttatggtgtattagctaaagcaaccagagagctgatgacgagaactgtggaaatcaggaatcctttggttaaaggctttgagat<br>tttccagtggacaaactatgccaagttctcaagcgaaaaattagaattagtttttagtgaagagatattgccttatcttttccagttaaaaaaattc<br>ataaaatataatctggaacatgttaagtcttttgaaaacaaatactctatgaggatttatgagtggttattaaaagaactaacacaaaagaaaac<br>tcacaaggcaaatatagagattagccttgatgaatttaagttcatgttaatgcttgaaaataactaccatgagtttaaaaggcttaaccaatgg<br>gttttgaaaccaataagtaaagatttaaacacttacagcaatatgaaattggtggttgataagcgaggccgcccgactgatacgttgattttcc<br>aagttgaactagatagacaaatggatctcgtaaccgaacttgagaacaaccagataaaaatgaatggtgacaaaataccaacaaccattac<br>atcagattcctacctacgtaacggactaagaaaaacactacacgatgctttaactgcaaaaattcagctcaccagttttgaggcaaaattttttg<br>agtgacatgcaaagtaagcatgatctcaatggttcgttctcatggctcacgcaaaaacaacgaaccacactagagaacatactggctaaat<br>acggaaggatctgaggttcttatggctcttgtatctatcagtgaagcatcaagactaacaaacaaaagtagaacaactgttcaccgttagata<br>tcaaagggaaaactgtccatatgcacagatgaaaacggtgtaaaaaagatagatacatcagagcttttacgagtttttggtgcatttaaagct<br>gttcaccatgaacagatcgacaatgtaactactagagggttcatgtgcagctccatcagcaaaaggggatgataagtttatcaccaccgact<br>atttgcaacagtgccgttgatcgtgctatgatcgactggatcggcggggcctgggacggcgcctcgccgtcgcagaaccaggcggtgg<br>cgtacaccgtcgcctcggtcggcccgtagagattggcgatcccgaccgcagcaccaccgagaacgtccccgacgtggccgaccagcc<br>cgtcatcgtcaacgcctgatccgcggtgcggacaggccgtgtcgcgaccggccgtgcggaattaagccggcccgtaccctgtgaatag<br>aggtccgctgtgacacaagaatccctgttacttctcgaccgtattgattcggatgattcctacgcgagcctgcggaacgaccaggagttctg<br>ggagccgctggcccgccgagccctggaggagctcgggctgccggtgccgccggtgctgcgggtgcccggcgagagcaccaacccc<br>gtactggtcggcgagcccggcccggtgatcaagctgttcggcgagcactggtgcggtccggagagcctcgcgtcggagtcggaggcg<br>tacgcggtcctggcggacgccccggtgccggtgccccgcctcctcggccgcggcgagctgcggcccggcaccggagcctggccgtg<br>gccctacctggtgatgagccggatgaccggcaccacctggcggtccgcgatggacggcacgaccgaccggaacgcgctgctcgccct<br>ggcccgcgaactcggccgggtgctcggccggctgcacagggtgccgctgaccgggaacaccgtgctcaccccccattccgaggtctt<br>cccggaactgctgcgggaacgccgcgcggcgaccgtcgaggaccaccgcgggtggggctacctctcgccccggctgctggaccgcc<br>tggaggactggctgccggacgtggacacgctgctggccggccgcgaaccccggttcgtccacggcgacctgcacgggaccaacatct<br>tcgtggacctggccgcgaccgaggtcaccgggatcgtcgacttcaccgacgtctatgcgggagactcccgctacagcctggtgcaactg<br>catctcaacgccttccggggcgaccgcgagatcctggccgcgctgctcgacggggcgcagtggaagcggaccgaggacttcgcccg<br>cgaactgctcgccttcaccttcctgcacgacttcgaggtgttcgaggagacgccgctggatctctccggcttcaccgatccggaggaactg<br>gcgcagttcctctgggggccgccggacaccgcccccggcgcctgacgccccgggccgcccggcgccgcccccggcccccggcgg<br>ccgccgggcggagccccgcccggcggcccgggagccccgggcccgcgccgaagcccgctgctgcgagctagctagagtcgacct<br>gggtccccggggatcggtcttgccttgctcgtcggtgatgtacttcaccagctccgcgaagtcgctcttcttgatggagcgcatggggacg<br>tgcttggcaatcacgcgcacccccggccgttttagcggctaaaaaagtcatggctctgccctcgggcggaccacgcccatcatgacctt<br>gccaagctcgtcctgcttctcttcgatcttcgccagcagggcgaggatcgtggcatcaccgaaccgcgccgtgcgcgggtcgtcggtga<br>gccagagtttcagcaggccgcccaggcggcccaggtcgccattgatgcgggccagctcgcggacgtgctcatagtccacgacgcccg<br>tgattttgtagccctggccgacggccagcaggtaggccgacaggctcatgccggccgccgccttttcctcaatcgctcttcgttcgtct<br>ggaaggcagtacaccttgataggtgggctgcccttcctggttggcttggtttcatcagccatccgcttgccctcatctgttacgccggcggta<br>gccggccagcctcgcagagcaggattcccgttgagcaccgccaggtgcgaataagggacagtgaagaaggaacacccgctcgcggg |

| | |
|---|---|
| | tgggcctacttcacctatcctgcccggctgacgccgttggatacaccaaggaaagtctacacgaaccctttggcaaaatcctgtatatcgtg<br>cgaaaaaggatggatataccgaaaaaatcgctataatgaccccgaagcagggttatgcagcggaaaagatccgtcgacctgtcacgccg<br>ccgctgtgaacccgttcagcgtctccggctccgtctcttcctcgaccttcggcttcgcccagtggatacgcacacggtcggcgatcgggg<br>gaacccgtccgggccgctgaacgcccttcggaagcttgatcacctcgatccggtccacgaagagccggacgaacgcccgacggtcttc<br>aaggggcgctagcgcccaccacgaacccggtcccgtcgggtcgctgccccggtcgcccagccattcgccgatcggcagcacggggg<br>agtcggcggcgtcgagctgacgaagccgttcttcggccccttccatgcggagcgtcagcgcggcttcctcttcgagaaacgcgcgccgt<br>cccatggcggaccggtagccgccgttccgcttgtcttcgtaaagctcttcgagcgccttcacggcgtccgcgcgctgtgccatcagttcgg<br>agcgctgacccttcaactccgggcgttcgtgcgtcgctccccagcgccgtgcggcttcgtacatgagcgctgccgtgtcgccttcgaggt<br>cgtcagggtcggcagggtcgaaggccgtgatgcgcgcgaagatcgcgccgacgatgtacgggtcaaggttgtgcatcgtgatcgagc<br>atgacgacccgtcatgaacccggcgggggcacttgcacgcgtacgacgacttgtgagcttgatcaccttcgcggacgttgcccgccatg<br>gtcgacccgttgctgtaacccgtctccgggtcgagctggccggagccgtagcagtaaaggacgtccattgccgacaggagcgattgccc<br>ccggtactgacccttcccgcgtcctcgaccctgaagccactcctgaagttcccaccattccgccgggggaatgtacggctcgaagccgg<br>gcagggtcagcggctccatggtgaccgggtcgcgcctgatcttgtaatggctgaagccgccccgcgaaccgtcggcgcgcaccttgtat<br>gcgatgtcagcttgatacccggcaatgcgcgggtcgctgagtacgcgcttcaaaacgccggggtcccaatcggaaccggcgcgcttctt<br>accgaccagcgtgccgcgcgtaggcaccttgtcgcggtacagccgctcacaaagcccgttcagcgacccagggtgaaacgacccggc<br>tccgccacccttgaagggcgtatcgcggtgcgtcttgatctcctgccacgcccagcggattaccgccccttcgctgccgtgtgcgccttcc<br>caggtgtgcgcgctgggcaccagtcggcgaatggcaaccagctttccgccgtcttccgggttcggaaccatttcctcgaccgtgtcgaat<br>ccgtacggcgtcgaccccgtgtgtccgcccagccgcttcgccaattccttagcgttcgacacggcgacgctcttgttcttcgactcatcatg<br>cgaagcctgaaggcgcatgatcaggtgaataaggtccatcatttcgccggggcggaatgtgccttcgttcacgctgacaatggtcacgcc<br>cagccggagcaattccgtgacgaccggaataatgtccagcggctcttcgcggctgaggcgcgaaatgtaatgaacaatgatcatgttcatt<br>tcccggttccggcacatgtccaaaatccggttgaactccggccggtcgacgcccgtgaatgccgacgtgccgggcgcttcgctgaagtg<br>acccagccacttcacctcgacgccgtcgcgcgcgtactccttcgccagcgcctcagccttcccccggttcgcggcgcgctgggtggccg<br>gtgaagcggtcgaactgttctcccgttccgccgactgtcggtcgtagccgcctgccagaatgaccatgccttcaaccttcacgctcatgcc<br>gggcaggctatcccgcccaactgggtaagagcaaagactaacactgttgggctggaactactcccgaaaaccgcttctgacctgggaaa<br>acgtgaagccccggggcatccgctgagggttgccgccggggcttcggtgtgtccgtcagtacgggccatagaggggcgtcgtgtaacc<br>cacccagggcgattgctccggcacggggaagccgcgccacgccttcgggacgtctggaatcgctagagcttgcatgc |

**Table S2: The sequences of $P_{T7}$ promoter variants**

| Promoter mutants (No.) | Sequence (5′—3′) |
|---|---|
| *kas*Op* | TGTTCACATTCGAACCGTCTCTGCTTTGACAAC ATGCTGTGCGGTGTTGTAAAGTCGTGGCCA |
| T7 | TAATACGACTCACTATAGGGG |
| 1 | CTAAACGACTCACTATAGGGG |
| 2 | AGATACGACTCACTATAGGGG |
| 3 | GTAAACGACTCACTATAGGGG |
| 4 | TGATACGACTCACTATAGGGG |
| 5 | ATTTACGACTCACTATAGGGG |
| 6 | GAATACGACTCACTATAGGGG |
| 7 | GATTACGACTCACTATAGGGG |
| 8 | CAAAACGACTCACTATAGGGG |
| 9 | ACTTACGACTCACTATAGGGG |
| 10 | TCCTACGACTCACTATAGGGG |
| 11 | GCCTACGACTCACTATAGGGG |
| 12 | GTGAACGACTCACTATAGGGG |
| 13 | ACCTACGACTCACTATAGGGG |
| 14 | CACAACGACTCACTATAGGGG |
| 15 | ATAGACGACTCACTATAGGGG |
| 16 | CGCAACGACTCACTATAGGGG |
| 17 | CGCACCGACTCACTATAGGGG |
| 18 | TGCAGCGACTCACTATAGGGG |
| 19 | GGTCGCGACTCACTATAGGGG |
| 20 | CTACACGACTCACTATAGGGG |
| 21 | TTTGTCGACTCACTATAGGGG |
| 22 | TATCGCGACTCACTATAGGGG |
| 23 | CACTACGACTCACTATAGGGG |
| 24 | AAAGCCGACTCACTATAGGGG |
| 25 | GCATTCGACTCACTATAGGGG |
| 26 | AGCCACGACTCACTATAGGGG |
| 27 | CTCATCGACTCACTATAGGGG |
| 28 | CCGTTCGACTCACTATAGGGG |
| 29 | CGAAACGACTCACTATAGGGG |
| 30 | AGGAACGACTCACTATAGGGG |
| 31 | CATCACGACTCACTATAGGGG |
| 32 | GACAACGACTCACTATAGGGG |
| 33 | AATCACGACTCACTATAGGGG |
| 34 | GGCTACGACTCACTATAGGGG |
| 35 | AGTTACGACTCACTATAGGGG |
| 36 | TAAAACGACTCACTATAGGGG |
| 37 | ATAAACGACTCACTATAGGGG |
| 38 | ATCTACGACTCACTATAGGGG |
| 39 | ACAAACGACTCACTATAGGGG |
| 40 | ATACACGACTCACTATAGGGG |
| 41 | TAAACCGACTCACTATAGGGG |

**Table S3 The sequence of T7 RNAP***

| Name | Sequence (5′—3′) |
|---|---|
| T7 RNAP* | atgaacacgattaacatcgctaagaacgacttctctgacatcgaactggctgctatcccgttcaacactctggctgaccattacggtgagc<br>gtCTCgctcgcgaacagttggcccttgagcatgagtcttacgagatgggtgaagcacgcttccgcaagatgtttgagcgtcaacttaaa<br>gctggtgaggttgcggataacgctgccgccaagcctctcatcactaccctactccctaagatgattgcacgcatcaacgactggtttgag<br>gaagtgaaagctaagcgcggcaagcgcccgacagccttccagttcctgcaagaaatcaagccggaagccgtagcgtacatcaccatta<br>agaccactctggcttgcctaaccagtgctgacaatacaaccgttcaggctgtagcaagcgcaatcggtcgggccattgaggacgaggct<br>cgcttcggtcgtatccgtgaccttgaagctaagcacttcaagaaaaacgttgaggaacaactcaacaagcgcgtagggcacgtctacaa<br>gaaagcatttatgcaagttgtcgaggctgacatgctctctaagggtctactcggtggcgaggcgtggtcttcgtggcataaggaagactct<br>attcatgtaggagtacgctgcatcgagatgctcattgagtcaaccggaatggttagcCTCcaccgccaaaatgctggcgtagtaggtca<br>agactctgagactatcgaactcgcacctgaatacgctgaggctatcgcaacccgtgcaggtgcgctggctggcatctctccgatgttcca<br>accttgcgtagttcctcctaagccgtggactggcattactggtggtggctattgggctaacggtcgtcgtcctctggcgctggtgcgtactc<br>acagtaagaaagcactgatgcgctacgaagacgtttacatgcctgaggtgtacaaagcgattaacattgcgcaaaacaccgcatggaaa<br>atcaacaagaaagtcctagcggtcgccaacgtaatcaccaagtggaagcattgtccggtcgaggacatccctgcgattgagcgtgaag<br>aactcccgatgaaaccggaagacatcgacatgaatcctgaggctctcaccgcgtggaaacgtgctgccgctgctgtgtaccgcaagga<br>caaggctcgcaagtctcgccgtatcagccttgagttcatgcttgagcaagccaataagtttgctaaccataaggccatctggttcccttaca<br>acatggactggcgcggtcgtgtttacgctgtgtcaatgttcaacccgcaaggtaacgatatgaccaaaggactgcttacgctggcgaaag<br>gtaaaccaatcggtaaggaaggttactactggctgaaaatccacggtgcaaactgtgcgggtgtcgataaggttccgttccctgagcgca<br>tcaagttcattgaggaaaaccacgagaacatcatggcttgcgctaagtctccactggagaacacttggtgggctgagcaagattctccgtt<br>ctgcttccttgcgttctgctttgagtacgctggggtacagcaccacggcctgagctataactgctcccttccgctggcgtttgacgggtcttg<br>ctctggcatccagcacttctccgcgatgctccgagatgaggtaggtggtcgcgcggttaacttgcttcctagtgaaaccgttcaggacatc<br>tacgggattgttgctaagaaagtcaacgagattctacaagcagacgcaatcaatgggaccgataacgaagtagttaccgtgaccgatga<br>gaacactggtgaaatctctgagaaagtcaagctgggcactaaggcactggctggtcaatggctggcttacggtgttactcgcagtgtgac<br>taagcgttcagtcatgacgctggcttacgggtccaaagagttcggcttccgtcaacaagtgctggaagataccattcagccagctattgatt<br>ccggcaagggtctgatgttcactcagccgaatcaggctgctggatacatggctaagctgatttgggaatctgtgagcgtgacggtggtag<br>ctgcggttgaagcaatgaactggcttaagtctgctgctaagctgctggctgctgaggtcaaagataagaagactggagagattcttcgca<br>agcgttgcgctgtgcattgggtaactcctgatggtttccctgtgtggcaggaatacaagaagcctattcagacgcgcttgaacctgatgttc<br>ctcggtcagttccgcCTCcagcctaccattaacaccaacaaagatagcgagattgatgcacacaaacaggagtctggtatcgctccta<br>actttgtacacagccaagacggtagccaccttcgtaagactgtagtgtgggcacacgagaagtacggaatcgaatcttttgcactgattca<br>cgactccttcggtaccattccggctgacgctgcgaacctgttcaaagcagtgcgcgaaactatggttgacacatatgagtcttgtgatgtac<br>tggctgatttctacgaccagttcgctgaccagttgcacgagtctcaattggacaaaatgccagcacttccggctaaaggtaacttgaacctc<br>cgtgacatcCTCgagtcggacttcgcgttcgcgtaa |

**Table S4** Oligonucleotides used in this study

| Primers | Sequence (5′—3′) |
|---|---|
| **Cloning and reconstruction of *act* gene cluster** | |
| guide RNA-F | gttttagagctagaaatagcaagttaaaataaggctagtc |
| guide RNA-R | aaaagcaccgactcggtgccactttttcaagttgataacggactagccttattttaact |
| sgRNA-actE | taatacgactcactatagg**gcgtccacgcgacctggatc**gttttagagctagaaatagcaa |
| sgRNA-actR | taatacgactcactatagg**cgcaagttttcgtgccggtt**gttttagagctagaaatagcaa |
| sgRNA-1 | taatacgactcactatagg**ccctcgagccgggttcgagt**gttttagagctagaaatagcaa |
| sgRNA-2 | taatacgactcactatagg**gtggtcgcgtcaagaaacgg**gttttagagctagaaatagcaa |
| sgRNA-3 | taatacgactcactatagg**tcgagcctccttcgagccac**gttttagagctagaaatagcaa |
| sgRNA-4 | taatacgactcactatagg**ttcggtgtgtccgtcagtac**gttttagagctagaaatagcaa |
| PF-1 | gctgttcacctggggagtgct |
| PR-1 | cgtagacgagcggcacgacac |
| PF-2 | taccgttcacaggtcgcggcg |
| PR-2 | gctcccgcgaacatcacgtc |
| P1yz-F | atctggtggccgtagaacga |
| P1yz-R | gcgctgaacaggttgatcca |
| P2yz-F | cgggaatctcctcgggcatg |
| P2yz-R | gtcggcttcaacggaactcatg |
| P3yz-F | agatggggtgccgctggac |
| P3yz-R | cgtgatgacgactctgcgcttc |
| Construction of pTHS-T7 RNAP* plasmid | |
| T7 RNAP* -F | cta*ggtctc*acctgatgaacacgattaacatcgctaag (*Bsa*I) |
| T7 RNAP* -R | ctg*ggtctc*tagctttacgcgaacgcgaagtccgact (*Bsa*I) |
| OLMA linkers for construction pPAS-$P_{T7(X)}$ series plasmids | |
| T7F | cctgTAATACGACTCACTATAGGGG |
| T7R | agctCCCCTATAGTGAGTCGTATTA |
| 1F | cctgCTAAACGACTCACTATAGGGG |
| 1R | agctCCCCTATAGTGAGTCGTTTAG |
| 2F | cctgAGATACGACTCACTATAGGGG |
| 2R | agctCCCCTATAGTGAGTCGTATCT |
| 3F | cctgGTAAACGACTCACTATAGGGG |
| 3R | agctCCCCTATAGTGAGTCGTTTAC |
| 4F | cctgTGATACGACTCACTATAGGGG |
| 4R | agctCCCCTATAGTGAGTCGTATCA |
| 5F | cctgATTTACGACTCACTATAGGGG |
| 5R | agctCCCCTATAGTGAGTCGTAAAT |
| 6F | cctgGAATACGACTCACTATAGGGG |
| 6R | agctCCCCTATAGTGAGTCGTATTC |
| 7F | cctgGATTACGACTCACTATAGGGG |
| 7R | agctCCCCTATAGTGAGTCGTAATC |
| 8F | cctgCAAAACGACTCACTATAGGGG |
| 8R | agctCCCCTATAGTGAGTCGTTTTG |

| | |
|---|---|
| 9F | cctgACTTACGACTCACTATAGGGG |
| 9R | agctCCCCTATAGTGAGTCGTAAGT |
| 10F | cctgTCCTACGACTCACTATAGGGG |
| 10R | agctCCCCTATAGTGAGTCGTAGGA |
| 11F | cctgGCCTACGACTCACTATAGGGG |
| 11R | agctCCCCTATAGTGAGTCGTAGGC |
| 12F | cctgGTGAACGACTCACTATAGGGG |
| 12R | agctCCCCTATAGTGAGTCGTTCAC |
| 13F | cctgACCTACGACTCACTATAGGGG |
| 13R | agctCCCCTATAGTGAGTCGTAGGT |
| 14F | cctgCACAACGACTCACTATAGGGG |
| 14R | agctCCCCTATAGTGAGTCGTTGTG |
| 15F | cctgATAGACGACTCACTATAGGGG |
| 15R | agctCCCCTATAGTGAGTCGTCTAT |
| 16F | cctgCGCAACGACTCACTATAGGGG |
| 16R | agctCCCCTATAGTGAGTCGTTGCG |
| 17F | cctgCGCACCGACTCACTATAGGGG |
| 17R | agctCCCCTATAGTGAGTCGGTGCG |
| 18F | cctgTGCAGCGACTCACTATAGGGG |
| 18R | agctCCCCTATAGTGAGTCGCTGCA |
| 19F | cctgGGTCGCGACTCACTATAGGGG |
| 19R | agctCCCCTATAGTGAGTCGCGACC |
| 20F | cctgCTACACGACTCACTATAGGGG |
| 20R | agctCCCCTATAGTGAGTCGTGTAG |
| 21F | cctgTTTGTCGACTCACTATAGGGG |
| 21R | agctCCCCTATAGTGAGTCGACAAA |
| 22F | cctgTATCGCGACTCACTATAGGGG |
| 22R | agctCCCCTATAGTGAGTCGCGATA |
| 23F | cctgCACTACGACTCACTATAGGGG |
| 23R | agctCCCCTATAGTGAGTCGTAGTG |
| 24F | cctgAAAGCCGACTCACTATAGGGG |
| 24R | agctCCCCTATAGTGAGTCGGCTTT |
| 25F | cctgGCATTCGACTCACTATAGGGG |
| 25R | agctCCCCTATAGTGAGTCGAATGC |
| 26F | cctgAGCCACGACTCACTATAGGGG |
| 26R | agctCCCCTATAGTGAGTCGTGGCT |
| 27F | cctgCTCATCGACTCACTATAGGGG |
| 27R | agctCCCCTATAGTGAGTCGATGAG |
| 28F | cctgCCGTTCGACTCACTATAGGGG |
| 28R | agctCCCCTATAGTGAGTCGAACGG |
| 29F | cctgCGAAACGACTCACTATAGGGG |
| 29R | agctCCCCTATAGTGAGTCGTTTCG |
| 30F | cctgAGGAACGACTCACTATAGGGG |

| | |
|---|---|
| 30R | agctCCCCTATAGTGAGTCGTTCCT |
| 31F | cctgCATCACGACTCACTATAGGGG |
| 31R | agctCCCCTATAGTGAGTCGTGATG |
| 32F | cctgGACAACGACTCACTATAGGGG |
| 32R | agctCCCCTATAGTGAGTCGTTGTC |
| 33F | cctgAATCACGACTCACTATAGGGG |
| 33R | agctCCCCTATAGTGAGTCGTGATT |
| 34F | cctgGGCTACGACTCACTATAGGGG |
| 34R | agctCCCCTATAGTGAGTCGTAGCG |
| 35F | cctgAGTTACGACTCACTATAGGGG |
| 35R | agctCCCCTATAGTGAGTCGTAACT |
| 36F | cctgTAAAACGACTCACTATAGGGG |
| 36R | agctCCCCTATAGTGAGTCGTTTTA |
| 37F | cctgATAAACGACTCACTATAGGGG |
| 37R | agctCCCCTATAGTGAGTCGTTTAT |
| 38F | cctgATCTACGACTCACTATAGGGG |
| 38R | agctCCCCTATAGTGAGTCGTAGAT |
| 39F | cctgACAAACGACTCACTATAGGGG |
| 39R | agctCCCCTATAGTGAGTCGTTTGT |
| 40F | cctgATACACGACTCACTATAGGGG |
| 40R | agctCCCCTATAGTGAGTCGTGTAT |
| 41F | cctgTAAACCGACTCACTATAGGGG |
| 41R | agctCCCCTATAGTGAGTCGGTTTA |
| URA amplification | |
| URA-F | ccgctgagggttgccgccggggcttcggtgtgtccgtcaggcaccacgcttttcaattcaattcatcatt |
| URA-R | cgccctgggtgggttacacgacgcccctctatggcccgtagggtaataactgatataattaaattgaagctcta |

**Table S5. The list of promoter cassettes and URA marker**

| Promoter | Sequence (5′—3′) [a] | Insertion position |
|---|---|---|
| bipromoter P1-P2 | ggaagcgcggtgatggtcatacggccacggtcgcccggtTggccacgACTTTAcacc<br>catgcccagtgagcTGTCAAactgccagctTTGACAaggtcccggtcttaggtgT<br>AAAGTcgtggccAgagtgggcgctgccaggctcgtcgccatgagcaccgtga | Between *actVA* and *actVI* (1) |
| P3 | cgacacgtgctcctcatcgtatggcatgaacgggccaccgtgttcacattcgaaccgtctctgct<br>TTGACAacatgctgtgcggtgttgTAAAGTcgatggcAtttcttgacgcgaccaccg<br>ttccattgtagaacggtggtc | Between *actII-orf1* and *actII-orf2* (2) |
| bipromoter P4-P5 | ctccctgcctcgtggtccctcacgcgctcagctttgggcTggccacgACTTTAcaccata<br>gcgcttgtccgTGTCAAcgagaaagccTTGACAacatgctgtgcggtgttgTAA<br>AGTcgggtgaAcacggggccgacgatgacgacgaccaccggacgaacgca | Between *act*III and *act*I-I (3) |
| URA | ccgctgagggttgccgccggggcttcggtgtgtccgtcaggcaccacgcttttcaattcaattca<br>tcatttttttttttattcttttttttgatttcggtttccttgaaatttttttgattcggtaatctccgaacagaag<br>gaagaacgaaggaaggagcacagacttagattggtatatatacgcatatgtagtgttgaagaaa<br>catgaaattgcccagtattcttaacccaactgcacagaacaaaaacctgcaggaaacgaagata<br>aatcatgtcgaaagctacatataaggaacgtgctgctactcatcctagtcctgttgctgccaagct<br>atttaatatcatgcacgaaaagcaaacaaacttgtgtgcttcattggatgttcgtaccaccaagga<br>attactggagttagttgaagcattaggtcccaaaatttgtttactaaaaacacatgtggatatcttga<br>ctgatttttccatggagggcacagttaagccgctaaaggcattatccgccaagtacaattttttact<br>cttcgaagacagaaaatttgctgacattggtaatacagtcaaattgcagtactctgcgggtgtata<br>cagaatagcagaatgggcagacattacgaatgcacacggtgtggtgggcccaggtattgttag<br>cggtttgaagcaggcggcagaagaagtaacaaaggaacctagaggccttttgatgttagcaga<br>attgtcatgcaagggctccctatctactggagaatatactaagggtactgttgacattgcgaaga<br>gcgacaaagattttgttatcggctttattgctcaaagagacatgggtggaagagatgaaggttac<br>gattggttgattatgacacccggtgtgggtttagatgacaagggagacgcattgggtcaacagt<br>atagaaccgtggatgatgtggtctctacaggatctgacattattattgttggaagaggactatttgc<br>aaagggaagggatgctaaggtagagggtgaacgttacagaaaagcaggctgggaagcatatt<br>tgagaagatgcggccagcaaaactaaaaaactgtattataagtaaatgcatgtatactaaactca<br>caaattagagcttcaatttaattatatcagttattaccctacgggccatagaggggcgtcgtgtaa<br>cccacccagggcg | (4) |

[a] The sequences highlighted in red are the homologous sequences corresponding to the target sites. The sequences highlighted in blue are promoter sequences.
